# Cell Communication Network factor 4 promotes tumor-induced immunosuppression in melanoma

**DOI:** 10.1101/2021.02.23.432584

**Authors:** Audry Fernandez, Wentao Deng, Sarah L. McLaughlin, Anika C. Pirkey, Stephanie L. Rellick, David J. Klinke

## Abstract

Immune cell composition within the tumor microenvironment is regulated by tumor-derived factors. Cell Communication Network factor 4 (CCN4/WISP1) is a matricellular protein secreted by cancer cells that promotes metastasis by inducing the epithelial-mesenchymal transition. While metastatic dissemination limits patient survival, the absence of anti-tumor immunity also associates with poor patient outcomes with recent work suggesting these two clinical correlates are linked. Motivated by finding that CCN4 was associated with a dampened anti-tumor immune contexture in patients diagnosed with primary melanoma, we tested for a direct causal link by knocking out CCN4 (CCN4-KO) in the B16F0 and YUMM1.7 mouse models for melanoma. Tumor growth was significantly reduced when CCN4-KO melanoma cells were implanted subcutaneously in immunocompetent C57BL/6 mice but not in immunodeficient NSG mice. Correspondingly, the frequency of total CD45^+^ tumor-infiltrating leukocytes was significantly increased in CCN4-KO tumors, with increased natural killer (NK) and effector CD8^+^ T cells and reduced myeloid-derived suppressor cells (MDSC). Additionally, the absence of tumor-derived CCN4 was associated with an impaired splenic generation of suppressive granulocytic MDSC. Among mechanisms linked to local immunosuppression, we found CCN4 directly suppressed antigen-induced IFN*γ* release by CD8^+^ T cells, promoted glycolysis and consequent lactate release by melanoma cells, and enhanced tumor secretion of MDSC-attracting chemokines like CCL2 and CXCL1. Finally, CCN4-KO in B16F0 and YUMM1.7 melanoma cells complemented the anti-tumor effect of immune checkpoint blockade (ICB) therapy. Overall, our results suggest that CCN4 promotes tumor-induced immunosuppression and is a potential target for therapeutic combinations with ICB.

**Statement of Significance:** Given emerging interest in understanding the interplay between functional plasticity and anti-tumor immunity, Cell Communication Network factor 4, a secreted matricellular protein linked to promoting metastasis in melanoma, also suppresses anti-tumor immunity.

## Introduction

Immune checkpoint blockade (ICB) has transformed the clinical landscape for treating patients diagnosed with melanoma. While these immunotherapies provide a significant benefit to a portion of the patient population, there is still unmet need for identifying targets that can broaden the clinical benefit of immunotherapies (1). As immune checkpoint blockade relies on inhibitory signals that can be present both in the local tumor microenvironment and in secondary lymphoid organs to augment anti-tumor immunity (2, 3), an ongoing anti-tumor immune response is a prognostic indicator of a clinical response (4, 5). Collateral targets that transform the tumor microenvironment from immunologically cold to hot are natural next steps. To date, a variety of targets have been identified that heat-up the anti-tumor immune response, including targets related to developmental pathways like the Wnt signaling pathway (6–8).

In addition to an immunologically cold tumor, metastatic dissemination also predicts limited patient survival (9). Metastatic dissemination is attributed to re-engaging developmental pathways that enable malignant cells to decouple from their tissue niche, migrate via the circulation, extravasate into a peripheral tissue, and establish metastatic colonies (10). In previous work, Cell Communication Network factor 4 (CCN4), a secreted matricellular protein produced by activating the Wnt/*β*-catenin pathway, promotes metastatic dissemination in melanoma by engaging the epithelial-mesenchymal transition (11, 12). Interestingly, we observed slower tumor growth rates of CCN4 knock-out variants in immunocompetent (C57BL/6) compared with immunocompromised (NSG) hosts while the parental cell lines exhibited no difference in growth rates. One explanation for this differential response is that CCN4 inhibits anti-tumor immunity. A secreted protein that promotes metastatic dissemination and simultaneously inhibits anti-tumor immunity is intriguing as neutralizing this protein therapeutically would impact two key limiters for patient survival. While a secretome screen piqued our interest in CCN4 (13), the literature related to CCN4 (a.k.a. WISP1) is not well-developed with less than 500 publications since 1993 listed in PubMed (14). To clarify the potential role that CCN4 plays at the interface of metastasis and anti-tumor immunity, the objective of this study was to test for clinical correlates in data obtained from humans diagnosed with melanoma and to assess a causal role for CCN4 in regulating host anti-tumor immunity more directly using immunocompetent mouse models for melanoma.

## Materials and Methods

### Mice

C57BL/6 mice (6-8 week-old, female) and NOD-scid IL2R*γ*^*null*^ immunodeficient mice (NSG, 6-8 week-old, male) were purchased from Charles River Laboratories and The Jackson Laboratory, respectively. Upon receipt, animals were labeled and randomly assigned to treatment arms/cages, with a density of five mice per cage. All animal experiments were approved by West Virginia University (WVU) Institutional Animal Care and Use Committee and performed at the WVU Animal Facility (IACUC Protocol #1604002138).

### Cell culture

All biochemical reagents were obtained from commercial sources and used according to the suppliers’ recommendations unless otherwise indicated. Mouse melanoma cell lines B16F0 and YUMM1.7 were cultured in supplemented DMEM as previously described (11). B16F0 cells (RRID: CVCL_0604) were obtained from the American Tissue Culture Collection (ATCC, Manassas, VA) in 2008. YUMM1.7 cells (RRID: CVCL_JK16) were a gift from Drs. William E. Damsky and Marcus W. Bosenberg (Yale University) (15) and were received in 2017. CCN4-knockout (CCN4-KO) B16F0 and YUMM1.7 cells were generated using a double nickase-based CRISPR/Cas9 approach as previously described (11). Additionally, for the B16F0 model, DNMT3A-and CCN4-knockout cells were obtained through transfection with a mix of CRISPR/Cas9 KO and Homology-Directed Repair (HDR) plasmids, followed by puromycin selection (12). Tet-on inducible mouse CCN4 expression lentiviral vector (IDmCCN4) was constructed with Gateway cloning using Tet-on destination lentiviral vector pCW57.1 (Addgene Plasmid #41393, a gift from David Root) and pShuttle Gateway PLUS ORF Clone for mouse CCN4 (GC-Mm21303, GeneCopoeia). Lentiviruses were packaged as described (11) to transduce YUMM1.7 cells with *Ccn4* CRISPR knockout (Ym1.7-KO1). Two pools of Tet-on variant cells with inducible mCCN4 (Ym1.7-KO1-IDmCCN4) or vector control (Ym1.7-KO1-IDvector) were obtained after puromycin selection. All cell lines were revived from frozen stock, used within 10-15 passages, and routinely tested for mycoplasma contamination by PCR.

### In vivo tumor growth and ICB immunotherapy

To evaluate the effect of CCN4-KO in B16F0 and YUMM1.7 melanoma tumors, 3 ×10^5^ tumor cells were subcutaneously (s.c.) injected in C57BL/6 and NSG mice. Once palpable, the largest perpendicular diameters of the s.c. tumors were measured unblinded with a caliper twice a week, and the tumor volume was calculated using the formula: *π*/6 x length x width^2^, where the width is the smaller dimension of the tumor. Using the resulting measurements, the growth rate of the tumors was estimated using a log-linear tumor growth model and a Markov Chain Monte Carlo approach to generate the posterior distribution in the rate parameters, as described previously (12). C57BL/6 mice were also injected subcutaneously with Tet-on variants constructed using CCN4 knockout YUMM1.7 cells (KO1) using a 2×2 factor experimental design, with doxycycline (Sigma-Aldrich) treatment and the Tet-on variant cells as the two factors. Doxycycline was delivered by injecting 0.15 mL (10 mg/mL) intraperitoneally at day 0 and orally via consumption (ad lib) of standard mouse chow containing 200 mg dox per 1 kg food (Bio-Serv). ICB therapy was studied using the InVivoMAbs anti-mouse PD1 (CD279, clone J43) monoclonal antibody with YUMM1.7 cell variants, anti-mouse CTLA-4 (clone UC10-4F10-11) monoclonal antibody with B16F0 cell variants, and a polyclonal Armenian hamster IgG as isotype control (IC) (BioXCell, NH) at doses of 200 *µ*g/mouse. C57BL/6 mice were s.c. inoculated with 5 ×10^5^ CCN4-KO and wild type (WT) YUMM1.7 tumor cells. The anti-PD1 and IC antibodies were administered intraperitoneally (i.p.) on days 0, 4 and 9, considering as day 0 the day when the tumors reached a volume of 100 mm^3^. All in vivo studies were repeated at least twice with two independent cohorts and with n ≥ 3 in each experimental group.

### In vitro suppression of CD8^+^ T cell function

To generate YUMM1.7-reactive CD8^+^ T cells, healthy C57BL/6 mice were inoculated subcutaneously with irradiated YUMM1.7 cells (10^5^/mouse), followed by live YUMM1.7 cells (3× 10^5^/mouse) 3 weeks later. The mice without tumors in the following five weeks were maintained. Three days before the assay, the mice were injected again with live YUMM1.7 cells (10^5^/mouse). On the day of assay, the YUMM1.7-reactive cells were isolated from mouse splenocytes using mouse CD8a+ T Cell Isolation Kit (Miltenyi Biotec, Germany), resuspended at 10^6^ cells/ml. 50*µ*l (5 ×10^4^) of the YUMM1.7-reactive CD8+ T cells were aliquoted into 96-well plates for ELISpot assay using Mouse IFN*γ*/TNF*α* Double-Color ELISpot kit (Cellular Technology Limited) following manufacturer’s instructions. Briefly, target tumor cells were stimulated with IFN*γ* (200U/ml, or, 20ng/ml) for 24 hours, harvested and resuspended at 2 × 10^6^ cells/ml. 50*µ*l (10^5^) of indicated tumor cells were aliquoted in triplicate, with or without doxycycline (Dox, final 0.5*µ*g/ml). The reactions were incubated at 37^*o*^C for 24 hours and colored spots were developed, imaged using an Olympus MVX10 Microscope, and counted.

### Flow cytometry

Tumors were surgically removed after euthanasia, weighted and processed into single cell suspensions using the Tumor Dissociation Kit, mouse (Miltenyi Biotec) and the manufacturer’s instructions. Single cell suspensions from tumors and spleens were stained with specific antibodies or IC using conventional protocols. Live and dead cells were discriminated with Live/Dead Fixable Violet Dead Cell Stain Kit (Thermo Fisher Scientific, MA). Fc receptors were blocked with purified rat anti-mouse CD16/CD32 (Mouse BD Fc Block, BD Biosciences, CA). Anti-mouse antibodies were used to characterize the lymphoid populations: CD45/BB515 (clone 30-F11, BD Biosciences), CD3*ϵ*/Alexa Fluor 700 (clone 500A2, BioLegend, CA), CD3*ϵ*/PE (clone 17A2, Miltenyi Biotec), CD8a/APC (clone 53-6.7, Miltenyi Biotec), CD4/APC-Cy7 (clone GK1.5, BD Biosciences), CD45R/B220/APC (clone RA3-6B2, BioLegend), NK-1.1/APC-Cy7 (clone PK136, BioLegend), CD49b/PerCP-Cy5.5 (clone DX5, BioLegend), CD25/PerCP-Cy5.5 (clone PC61.5, eBioscience), CD279 (PD-1)/PE (clone REA802, BioLegend), and FOXP3/PE (clone FJK-16s, eBioscience). The following antibodies were used to detect the myeloid populations: CD45/BB515 (clone 30-F11, BD Biosciences), CD11b/PerCP-Cy5.5 (clone M1/70, Thermo Fisher Scientific), Ly-6G/Ly-6C (Gr-1)/APC (clone RB6-8C5, BioLegend), CD11c/PE (clone N418, Thermo Fisher Scientific), F4/80/APC (clone BM8, BioLegend), I-A/I-E/Alexa Fluor 700 (clone M5/114.15.2, BioLegend), Ly-6G/APC (clone 1A8, BD Biosciences), and Ly-6C/PE (clone AL-21, BD Biosciences). Total number of cells in tumors and spleens were determined using SPHERO AccuCount Fluorescent Particles (Spherotech, IL). Expression of H-2K^*b*^ and PD-L1 were assayed in the CD45^−^ fraction generated from WT and CCN4-KO YUMM1.7 tumors using the antibodies: CD274 (PD-L1)/PE (clone 10F.9G2, BioLegend) and H-2K^*b*^/APC (clone AF6-88.5, BioLegend). For comparison, H-2K^*b*^ and PD-L1 expression was assayed in WT and CCN4-KO YUMM1.7 tumor cells conditioned in the presence or absence of IFN*γ* (200U/ml, or, 20ng/ml) for 24 hours, with unstained cells as controls. Events were acquired using a BD LSRFortessa (BD Biosciences) flow cytometer with FACS-Diva software, where the fluorescence intensity for each parameter was reported as a pulse area with 18-bit resolution. Flow cytometric data were exported as FCS3.0 files and analyzed with FCS Express 6.0 (DeNovo Software, CA) and FlowJo 5.7.2 (Tree Star Inc., OR). The typical gating strategies for lymphoid and myeloid cells are shown in Figures S1 and S2, respectively.

### T cell proliferation and MDSC-mediated suppression assay

Splenocytes from mice bearing CCN4-KO and YUMM1.7-WT tumors, as well as from tumor-free mice, were used as effector cells. CD45^+^ cells isolated from tumor-bearing (TB) mice using the CD45 MicroBeads, mouse (Miltenyi Biotec) were also used as an effector population. All effector cells (1 × 10^7^/mL) were stained with CellTrace Violet Cell Proliferation Kit (Thermo Fisher Scientific) following the manufacturer protocol. Once stained, 5 × 10^5^ effector cells/well were stimulated for 72 h with the T Cell Activation/Expansion Kit, mouse (Miltenyi Biotec) at a 1:1 ratio with anti-CD3/anti-CD28-loaded beads. To evaluate the suppressive function, granulocytic MDSC (G-MDSC) were isolated using the Myeloid-Derived Suppressor Cell Isolation Kit, mouse (Miltenyi Biotec) from the spleens of CCN4-KO and YUMM1.7-WT TB mice and tumor-free mice. G-MDSC (20% of effector cells) were then co-incubated for 72 h with 5 × 10^5^ stained naïve splenocytes in the presence of anti-CD3/anti-CD28-loaded beads at 1:1 ratio with effector cells. Proliferation diluted the CellTrace Violet dye, as assayed by flow cytometry. Live CD8^+^ effector cells were identified with Live/Dead Fixable Green Dead Cell Stain Kit (Thermo Fisher Scientific) and anti-mouse CD8a/APC (clone 53-6.7, Miltenyi Biotec).

### Tumor-conditioned media collection, cytokine array and ELISA assays

To collect tumor-conditioned media (TCM), cells were grown in complete DMEM medium until 80% confluency, washed with PBS (Cellgro/Corning, NY) and incubated for 48 h in FBS-free DMEM medium. TCM were then centrifuged at 3000xg and 4^*°*^C for 15 min with the supernatant collected and filtered. The cytokines, chemokines and growth factors in TCM were detected with the Proteome Profiler Mouse XL Cytokine Array (R&D Systems, MN), following the manufacturers instructions. CCN4 was assayed in TCM using the mouse WISP-1/CCN4 DuoSet ELISA Kit (R&D Systems).

CCL2 and CXCL1 were also quantified with Mouse CCL2/JE/MCP-1 and Mouse CXCL1/KC DuoSet ELISA kits (R&D Systems), respectively. These chemokines were measured in TCM, obtained from the cell lines in vitro as described above, and from CD45^−^ cells isolated from WT and CCN4-KO tumors by negative selection using a mouse CD45 MicroBeads kit (Miltenyi Biotec). Conditioned media were obtained by culturing 1 ×10^4^ CD45^−^ cells/well for 36 h in DMEM supplemented with 10% FBS. Blood was collected from the submandibular vein of CCN4-KO and YUMM1.7-WT TB mice and the serum was obtained after letting the blood to clot for 1 h at room temperature and performing a 10 min. centrifugation at 2000xg and 4^*°*^C and assayed for CCL2 and CXCL1.

### Metabolic function assays

WT and CCN4-KO melanoma cells were cultured overnight (1 × 10^4^ cells/well) in Seahorse XFe96 cell culture microplates (Agilent, CA) with complete DMEM medium. The Extracellular Acidification Rate (ECAR) was measured using a Seahorse XFe96 Analyzer (Agilent) according to the manufacturer’s instructions, which allowed calculating glycolysis and glycolytic capacity. To compare the lactate secretion, live CD45^−^ cells were isolated by negative selection from digested WT and CCN4-KO tumors using the mouse CD45 (TIL) MicroBeads kit (Miltenyi Biotec). A total of 1 ×10^4^ cells/well were cultured for 36 h in DMEM supplemented with 10% dialyzed FBS (Gibco, Thermo Fisher Scientific). Lactate was measured in the conditioned media using the Lactate-Glo Assay (Promega, WI).

### Statistical Analysis

Gene expression and clinical profiles for patients diagnosed with stage I to III melanoma (SKCM) from TCGA were downloaded using the “TCGAbiolinks” (V2.8.2) package in R (V3.5.1). Single-cell RNA sequencing data obtained from tumor samples of patients diagnosed with melanoma that were naïve and resistant to immune checkpoint therapy were used with Gene Expression Omnibus accession numbers GSE72056 and GSE115978, where non-zero counts in CCN4 expression was used to designate a CCN4-positive malignant cell. Statistical enrichment of CCN4-positive malignant cells was assessed by a binomial test where the observed frequency was compared against a null hypothesis represented by a binomial distribution with a baseline frequency of 1% CCN4-positive cells. A p-value represents the probability of the observed or greater frequency being drawn from null distribution. The immune contexture was estimated from the SKCM data obtained from primary melanoma tissue samples using CIBERSORTx and the LM22 immune cell gene signatures (16). Statistical differences in the posterior distributions in tumor growth rate parameters were assessed using a Pearson’s Chi-squared test. Kaplan-Meier analysis, Cox proportional hazards modeling, and Mann-Whitney U tests were performed using the “survival” (V2.42-6), “survminer” (V0.4.2), and “stats” (V3.5.1) packages in R. Unless otherwise specified, quantitative results were summarized as mean ±standard error of measurement (SEM) and overlaid on individual results. Unpaired Student’s t-test (two-tailed) or one-way ANOVA followed by Tukey’s multiple comparison ad hoc post-test were performed with GraphPad Prism (version 5). A p-value of < 0.05 was considered statistically significant and denoted as follows: * = 0.01<p<0.05, ** = 0.001<p<0.01, and *** = p<0.001.

## Results

### CCN4 is associated with a reduced anti-tumor immune contexture in primary melanoma patients

To assess the clinical context, we first tested for possible connection between *CCN4* mRNA expression and overall survival of patients diagnosed with primary melanoma and reported in skin cutaneous melanoma (SKCM) arm of the Cancer Genome Atlas (TCGA). Using data from samples obtained at diagnosis from patients with primary melanoma and with complete survival histories for statistical analysis (n = 95), we stratified patients based on *CCN4* expression and summarized their overall survival using Kaplan-Meier survival curves (Fig. 1A). A Cox proportional hazards model was used to assess covariance of overall survival with *CCN4* expression, tumor stage, age at diagnosis, and gender (Fig. S3). *CCN4* expression was the only statistically significant covariate (HR 2.24, p-value = 0.022).

**Figure 1.**
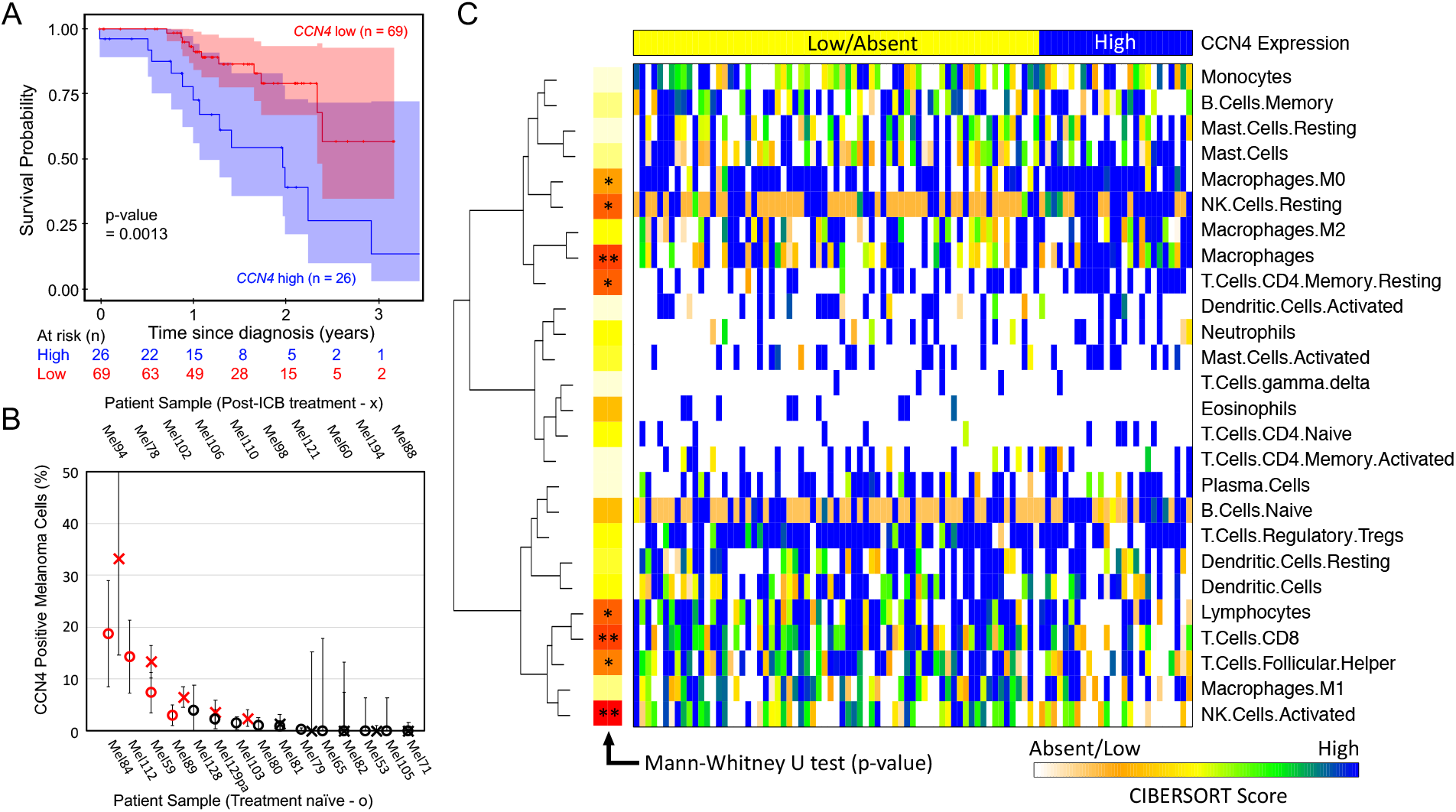
CCN4 is associated with reduced overall survival of patients diagnosed with primary melanoma and a shift in immune contexture. (A) Kaplan-Meier estimate of overall survival of patients diagnosed with primary melanoma stratified by CCN4 transcript abundance, with patients at risk tabulated below graph, as similarly shown in (11). Original data obtained from SKCM arm of TCGA and stratified based on CCN4 mRNA expression (CCN4 high/positive > 1 FPKM: blue, CCN4 low/negative < 1 FPKM: red). P-value calculated using the Peto & Peto modified Gehan-Wilcoxon test. (B) The proportion of CCN4 positive melanoma cells obtained from patients with melanoma obtained prior to treatment (o, n = 15) and from patients that did not respond to immune checkpoint blockade (x, n = 10). Values shown as the proportion of CCN4 positive melanoma cells of the sample ± SE of the sample proportion, given a binomial distribution. A binomial test assessed significance between the proportion of CCN4 positive cells in the sample relative to a null proportion of 1% or less CCN4 positive melanoma cells. Samples with significantly enriched CCN4 positive cells are indicated in red. (C) Immune contexture in corresponding primary SKCM tissue samples estimated from bulk RNAseq data using CIBERSORTx deconvolution. Columns ordered from low (left) to high (right) CCN4 expression. Rows hierarchically clustered based on a Euclidian distance metric in R (Ward.D). A non-parametric Mann-Whitney U test assessed significance of difference in immune subset signature between CCN4 high/positive and low/negative groups. The log10 p-values were color coded.

As the TCGA data are derived from a bulk tissue assay that averages across malignant, stromal, and immune cells present within the tissue sample, we used single cell RNA-seq data obtained from melanoma biopsied prior to treatment (n = 15) and following non-response to immune checkpoint blockade (ICB, n = 10) to assess the frequency of melanoma cells producing *CCN4* (Fig. 1B). Assuming a null hypothesis of 1% or less of CCN4 positive cells could be observed by random chance, a binomial test was used to assess whether the proportion of CCN4 positive cells in each patient sample is greater than the null hypothesis. The samples with a frequency of CCN4-positive cells greater than 1% with 95% confidence were considered CCN4 positive. In the treatment naïve cohort, 4 of 15 samples had CCN4 positive tumors. In the ICB resistant cohort, 5 of 10 samples had CCN4 positive tumors. In comparison, the TCGA SKCM primary melanoma cohort had 26 of 95 samples with CCN4 positive tumors (CCN4 reads greater than 1 FPKM). Using a Fisher Exact test, the difference in CCN4 positive samples in the treatment naïve group (4 of 15) compared to the TCGA SKCM cohort (26 of 95) was not statistically different (p-value = 1, odds ratio = 0.965). While the prevalence of CCN4 positive samples in the ICB resistant cohort (5 of 10) seemed to be higher compared to treatment naïve and TCGA SKCM cohorts (p-value = 0.1535, odds ratio = 2.64), we are unable to conclude with confidence that these samples come from different populations as n = 10 is not sufficiently powered to detect a difference in prevalence between 27% and 50%. Collectively, the results suggest that (1) the frequency of CCN4 positive tissue samples is similar among all of the cohorts, (2) patients with higher levels of CCN4 expression have a worse outcome, and (3) separating the population into high and low CCN4 expression subsets is not masking for other common latent variables that may influence overall survival, such as age, sex, and tumor stage.

Given that the clinical data suggest that melanoma patients with high *CCN4* expression have a worse outcome, we used digital cytometry to infer whether changes in immune contexture also corresponded with changes in *CCN4* expression (Fig. 1C). In the SKCM TCGA dataset, gene signatures associated with CD8 T cells (p-value < 0.01), activated NK cells (p-value < 0.01), follicular T helper cells (p-value < 0.05), and lymphocytes (p-value < 0.05) were reduced while M0 macrophages (p-value < 0.05), resting NK cells (p-value < 0.05), resting memory CD4^+^ T cells (p-value < 0.05), and macrophages (p-value < 0.01) were increased in the high *CCN4* cohort. Collectively, characterizing the immune contexture in human melanoma using digital cytometry suggests that increased *CCN4* expression corresponds with a shift in immune response from active anti-tumor immunity to a dampened immune response with enhanced resting and undifferentiated immune cell phenotypes. Animal models may help clarify mechanistic underpinnings of these clinical observations.

### CCN4 knockout in mouse melanoma reduces subcutaneous tumor growth

To analyze the importance of CCN4 for subcutaneous (s.c.) tumor growth and anti-tumor immune response, we used two mouse melanoma models: the spontaneous B16F0 model and the more clinically relevant YUMM1.7 model displaying Braf^*V* 600*E/W T*^ Pten^−*/*−^Cdkn2^−*/*−^ genotype. B16F0 and YUMM1.7 parental cells secreted 605±15 pg/ml and 451± 25 pg/ml, respectively, of CCN4 in 2D culture media. CCN4-KO variants of the B16F0 and YUMM1.7 cell lines were generated through CRISPR/Cas9 methodology and produced undetectable levels of CCN4 under similar culture conditions (Fig. 2A). A control for puromycin selection (B16F0-Ctr) produced CCN4 similar to the parental cell line.

**Figure 2.**
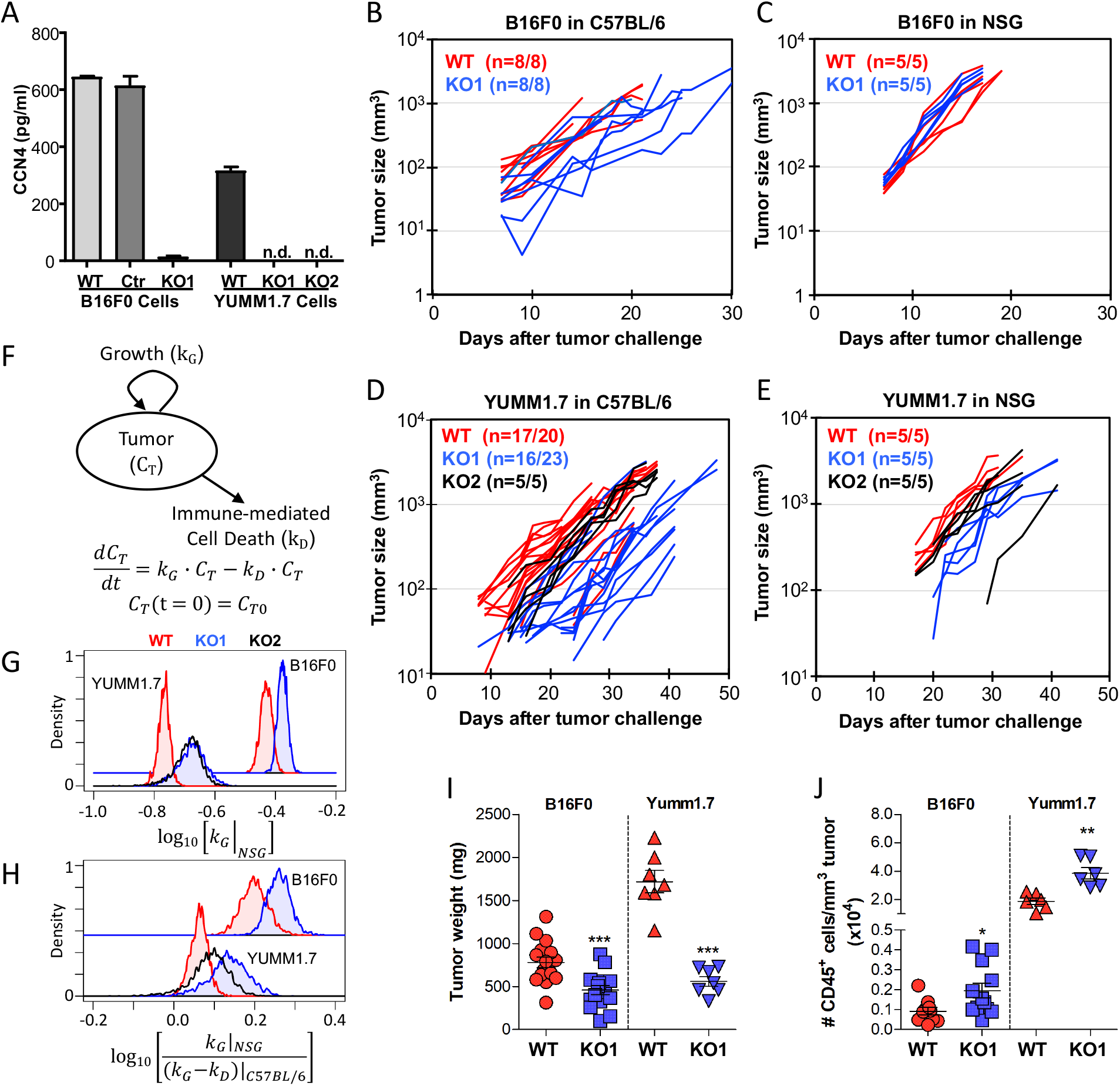
CCN4 knockout suppressed melanoma tumor growth in immunocompetent mice. (A) CCN4 secretion after CRISPR/Cas9 knockout in B16F0 and YUMM1.7 cell lines. Cell culture media conditioned for 48 hours by the indicated cell line were tested by ELISA (n.d. = not detected). B16F0 knockout created with Homology-Directed Repair approach with controls for puromycin selection (B16F0-Ctr) created with pBabe-puro retrovirus. YUMM1.7 knockout created with double nickase approach with KO1 and KO2 indicating two different clones. Wildtype (WT: red) B16F0 (B,C) and YUMM1.7 (D,E) cells and CCN4 knockout variants (KO1: blue, KO2: black) were subcutaneously injected into (B,D) C57BL/6 immunocompetent and (C,E) NSG immunocompromised mice. Tumor volumes measured by caliper as a function of time after tumor challenge, with n expressed as the number of mice with tumors over the number of mice injected. (F) The tumor growth model used to interpret the log-linear tumor growth curves. Posterior distributions in the growth rate constant of tumors in NSG mice (G) and in the log-ratio of the net growth rate constants of tumors in NSG relative to C57BL/6 mice (H). (I) In a time-matched experiment, average tumor weights of both B16F0 and YUMM1.7 were significantly decreased following CCN4 knockout (KO1). (J) Significant increases in the number of CD45^+^ cells were observed in both B16F0 and YUMM1.7 cell lines following CCN4 knockout (KO1).

Following s.c. implantation, we compared the growth trajectories of CCN4-KO variants with parental B16F0 (Fig. 2B and C) and YUMM1.7 (Fig. 2D and E) cells in immunocompetent C57BL/6 (Fig. 2B and D) and in severely immuno-compromised NSG (Fig. 2C and E) mice. Tumor growth consistently followed a log-linear growth trajectory, which implies that tumor size at any time point depends on the initial bolus of tumor-initiating cells (*C*_*T* 0_) and the net growth rate constant (*k* = *k*_*G*_ − *k*_*D*_) (Fig. 2F). Between these two parameters, the log growth rate constants were more consistent among replicates than the initial bolus of tumor-initiating cells, despite injecting the same number of cells. As genetic editing of the cells can alter the intrinsic growth rate, we used tumor growth in NSG mice to estimate the intrinsic growth rate constants (*k*_*G*_) for WT and CCN4 KO variants (Fig. 2G). We noted that CCN4 KO increased *k*_*G*_ for both cell lines (p-value < 1E-5). In immunocompetent mice, we assumed that the net growth rate constant reflects both the intrinsic growth rate constant and a loss in tumor size due to immune-mediated cell death (*k*_*D*_), which lumps all mechanisms for anti-tumor immunity that are present in C57BL/6 mice and absent in NSG mice into a single constant. Of note, any residual anti-tumor immune function present in NSG mice, albeit nearly negligible, is included in the intrinsic growth rate constant. We can infer the impact of CCN4 KO on immune-mediated cell death by comparing the difference in net growth rate constant for a given cell line in NSG versus C57BL/6 mice, which here is expressed as a log-ratio (Fig. 2H). Generally, YUMM1.7 variants exhibited lower log-ratios compared to B16F0 variants (e.g., median log-ratio B16F0 WT = 0.1991 versus YUMM1.7 WT = 0.063), which is consistent with genetically engineered mouse models being less immunogenic compared to spontaneous tumor models. More importantly, CCN4 KO variants of both cell lines exhibited greater log-ratios relative to WT cell lines (p-values all < 1E-5). KO1 variants for both cell lines were used for subsequent experiments. Of note, B16F0 variants with a knock-out of DNA (cytosine-5)-methyltransferase 3A (DNMT3A-KO), which served as a CRISPR/Cas9 editing control, exhibited no difference in tumor growth and overall survival (Fig. S4) compared with parental B16F0 cells. After 14 days following tumor challenge in the B16F0 model, tumors were surgically removed and weighted, with CCN4-KO tumors being 1.7-fold smaller than WT tumors (Fig. 2I, p-value=0.0003). After 28 days in the YUMM1.7 model, excised CCN4-KO tumors were 3-fold lighter than WT tumors (Fig. 2I; p-value<0.0001). Given the different growth response to CCN4 knockout in C57BL/6 versus NSG mice, the results suggest that CCN4 plays a role in modulating the immune system to favor tumor development. Supporting this idea, a 2-fold increase of infiltrating CD45^+^ leukocytes was detected in CCN4-KO tumors when compared to B16F0-WT (p-value=0.0297) and YUMM1.7-WT (p-value=0.0017) tumors (Fig. 2J). Subsequent experiments focused on clarifying CCN4’s role in modulating anti-tumor immunity.

### CCN4 knockout increases CTL and NK effector cells frequency while reducing MDSC infiltration in melanoma tumors

Next, we used flow cytometry to characterize tumor-infiltrating leukocytes using three different antibody panels that focused on T cells, NK cells, and myeloid cell subsets (Fig. 3). In resolving lymphoid subsets, we found that CD3e^+^ T cells (Live CD45^+^ CD3e^+^ events, Fig. 3A and B; p-value=0.0046), CD4^+^ T cells (Live CD45^+^ CD3e^+^ CD4^+^ CD8a^−^ events, Fig. 3A and C; p-value=0.0006), CD8^+^ T cells (Live CD45^+^ CD3e^+^ CD8a^+^ CD4^−^ events, Fig. 3A and D; p-value=0.0004), and NK cells (Live CD45^+^ CD3e^−^ NK1.1^+^ B220^−*/lo*^ events, Fig. 3A and E; p-value=0.0104) were increased in CCN4-KO tumors compared to YUMM1.7-WT tumors. In terms of the myeloid compartment, a subset of macrophages (Live CD45^+^ CD11b^+^ CD11c^+^ Gr1^−^ F480^+^ MHCII^+^ events, Fig. 3F; p-value<0.0001) and neutrophils (Live CD45^+^ CD11b^*lo*^ CD11c^−^ Gr1^+^ events, Fig. 3A and G; p-value=0.0004) were significantly expanded in the CCN4-KO tumors. Interestingly, these macrophages were CD11c^+^ (Fig. 3A), a marker associated with pro-inflammatory M1 TAM (17). Compared to CD11c^+^ subset, CD11c^−^ macrophages were 100-times less abundant and not statistically different. No significant differences were detected in the number of dendritic cells (DC: Live CD45^+^ CD11b^−^ CD11c^+^ Gr1^−^ F480^−^ events, Fig. 3H; p-value=0.0699). We also noted the number of CD8a^+^ DCs (Live CD45^+^ CD3e^−^ CD8a^+^ events) was negligible. Conversely, putative myeloid-derived suppressor cells (MDSC) were reduced (Live CD45^+^ CD11b^+^ Gr1^+^ events, Fig. 3A and I; p-value=0.0078), which resulted in the CCN4-KO tumors having 4.4-fold (p-value<0.0001) and 3.5-fold (p-value=0.0013) higher CD8^+^ T cells/MDSC and NK/MDSC ratios, respectively (Fig. S5). Similar results were obtained with the B16F0 tumor model (Fig. S5).

**Figure 3.**
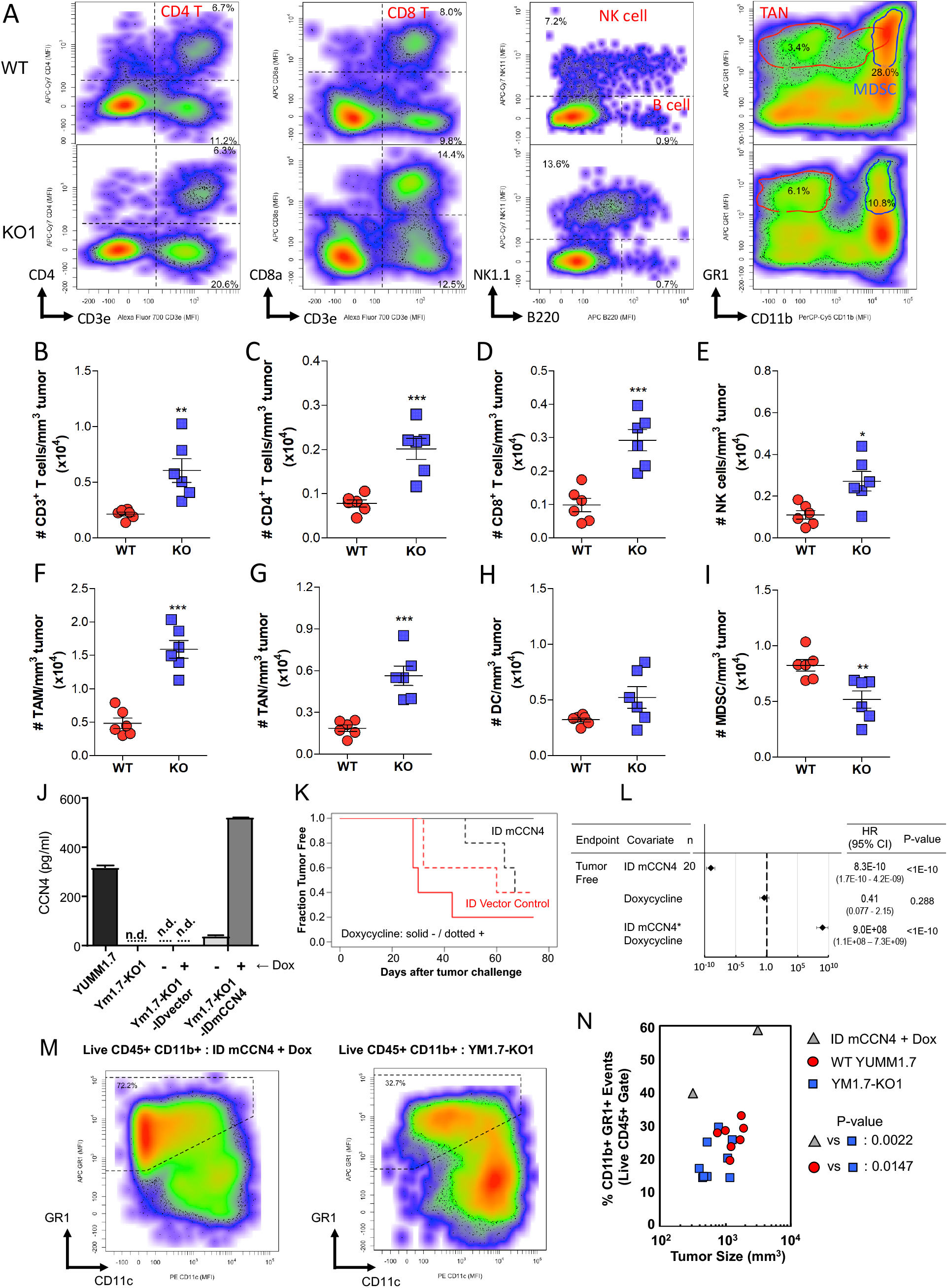
CCN4 knockout increased CTL and NK effector cells and decreased MDSC in the tumor microenvironment. (A) Representative flow cytometry data of the frequency of tumor-infiltrating CD4^+^ and CD8^+^ T cells, NK effector cells, tumor-associated neutrophils (TAN) and MDSC from the live CD45^+^ compartment in WT and CCN4 knockout (KO1) YUMM1.7 tumors. In the right panel, the contour lines enclose 90% of TANs (red line) and MDSC (blue) that were determined based on gating shown in Fig. S2 but here backgated along the GR1/CD11b axes. (B-I) Graphs summarizing the change in frequency of immune cells in the YUMM1.7 tumor microenvironment following CCN4 knockout (n=6 /group). (B) CD3^+^, (C) CD4^+^, (D) CD8^+^, (E) NK cells, (F) CD11c^+^ TAMs, (G) TANs, (H) DCs, and (I) MDSC. (J) CCN4 secretion from CCN4-inducible cells and associated control cell lines in conditioned media with or without 0.5 *µ*g/ml doxycycline, as measured by ELISA (n.d. = not detected). (K) Kaplan-Meier summary of the fraction tumor free in a 2×2 factor experimental design (n = 5 / group), where Tet-on vector control (red) versus inducible mCCN4 vector (black) and in the presence (dotted lines) or absence (solid lines) of doxycycline were the two factors. (L) Summary of a Cox multivariate proportional hazards analysis. (M) Representative flow cytometry data of the frequency of tumor-infiltrating MDSC (GR1^+^ CD11c^−*/lo*^ : dotted box) from the live CD45^+^ CD11b^+^ compartment isolated from CCN4-induced rescue (ID mCCN4 + Dox) and CCN4 knockout (YM1.7-KO1) YUMM1.7 tumors. (N) Frequency of MDSC for similarly sized tumors generated from CCN4-induced rescue (gray triangle), CCN4 knockout (blue square), and WT YUMM1.7 cells (red circles).

To test whether reintroducing CCN4 into CCN4-KO cells rescues the phenotype, we created a CCN4-inducible variant of CCN4 KO YUMM1.7 cells under the control of doxycycline and a vector control. CCN4 expression was under stringent control of doxycycline, with induced levels similar to wild-type YUMM1.7 cells (Fig. 3J). C57BL/6 mice were challenged with the Tet-on inducible CCN4 (ID mCCN4) and vector control variants of the YUMM1.7 CCN4 KO cells (Ym1.7-KO1) and doxycycline using a 2×2 factor experimental design (Fig. 3K). Tumor free survival of the mouse cohort was regressed using a Cox proportional hazards model jointly to the presence of the ID mCCN4 expression vector, treatment with DOX, and the combination (Fig. 3L). As expected, genetic editing of the Ym1.7-KO1 cells to include the ID mCCN4 vector reduced the hazard ratio (p-value <1e-10) while the treating with doxycycline had no significant effect (p-value = 0.288). Re-expression of CCN4 significantly increased tumor development (p-value < 1e-10). The presence of putative tumor-infiltrating MDSC were increased by CCN4 re-expression compared with similarly sized WT and CCN4 KO tumors (Fig. 3M-N, p-value=0.0022). Collectively, these data suggest that CCN4 skews the immune contexture within the melanoma microenvironment by reducing the infiltration of cells with potential anti-tumor cytolytic activity, namely CD8^+^ T cells and NK cells, and by enhancing the prevalence of cells with immunosuppressive capacity, namely MDSC.

### Melanoma-produced CCN4 promotes the splenic expansion of G-MDSC

Considering the spleen’s role in tumor-induced immunosuppression and particularly for MDSC extramedullary generation (18, 19), we analyzed the frequency of MDSC in the spleen of CCN4-KO and YUMM1.7-WT tumor-bearing (TB) mice. Splenic MDSC were first compared on day 28 after tumor challenge (time-matched, Fig. 4A), where both the percentage (Fig. 4B, p-value=0.0074) and number of MDSC per gram of spleen (Fig. 4C, p-value=0.0019) were significantly reduced in CCN4-KO TB mice. Splenomegaly was also notable in mice bearing YUMM1.7-WT tumors (Fig. S6A-C). Moreover, the percentage of CD45+ cells and total number per gram spleen of CD11b^+^Gr1^+^ cells in mice with CCN4-KO tumors were similar to the normal values detected in tumor-free mice (Fig. 4B and C, p-value=0.8886 and p-value=0.5042, respectively). However in matching the time point, the results raised an additional concern as CCN4-KO tumors were 3.7-fold smaller than YUMM1.7-WT (Fig. 4A, p-value=0.0277) and MDSC prevalence commonly correlates with tumor burden.

**Figure 4.**
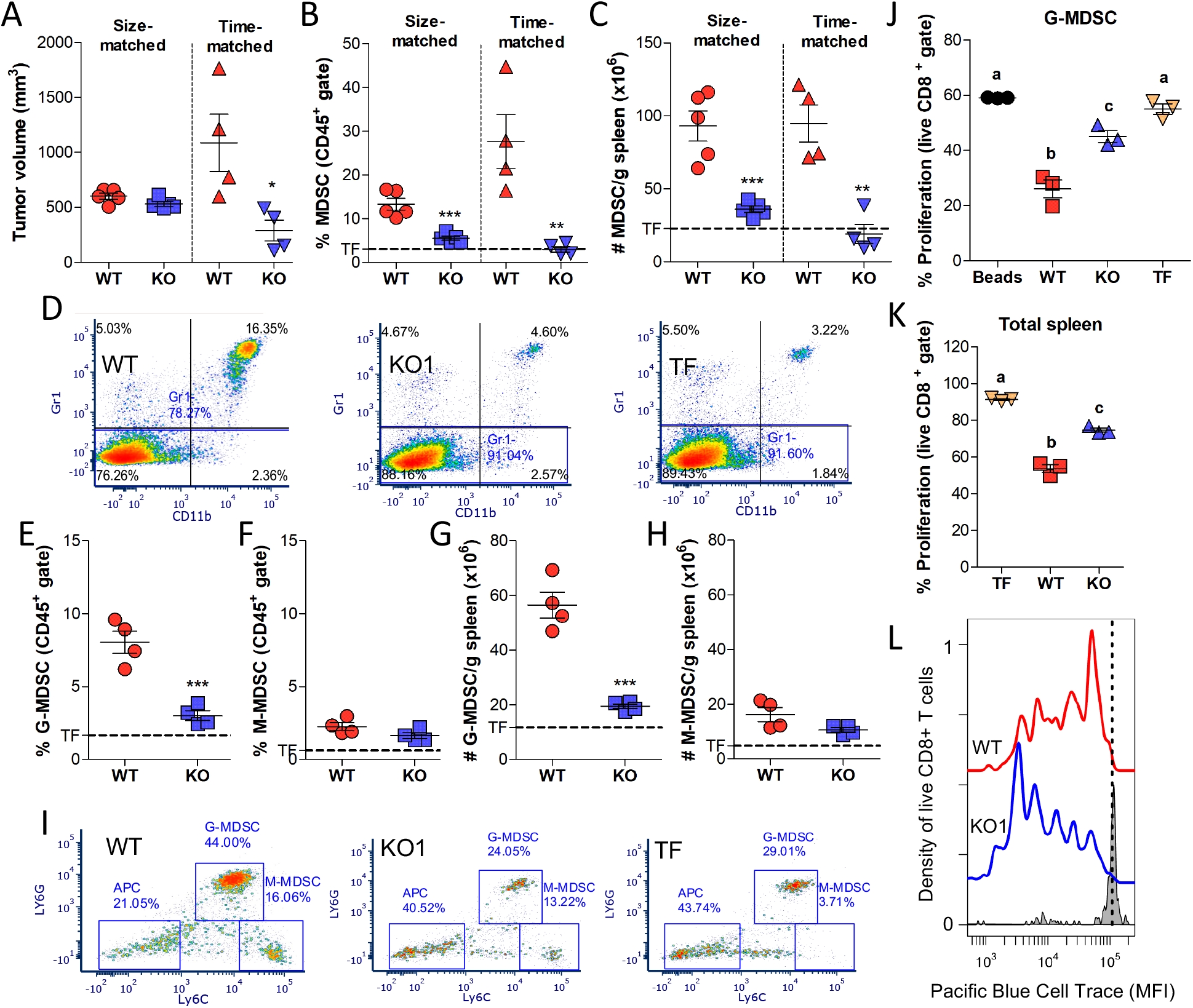
CCN4 expanded MDSC in the spleens of tumor bearing mice. (A) Tumor volume of YUMM1.7-WT and CCN4 KO (KO1) compared when size-matched and time-matched on day 28 after tumor challenge. (B-C) MDSC in the spleens of tumor bearing mice expressed as percentage (B) or weight-basis (C) observed in size-matched versus time-matched experimental designs (n = 5 for size-matched, 4 for time-matched experiments). (D) Representative flow cytometry data of CD11b^+*/*−^/GR-1^+*/*−^ cells found in WT, KO1, and tumor free (TF) groups. (E-H) Graphs summarizing G-MDSCs (E,G) and M-MDSCs (F,H) in the spleens of time-matched YUMM1.7-WT and CCN4 KO (KO1) tumor bearing mice (n = 4/group) expressed as percentage of CD45^+^ (E,F) and total number per gram spleen (G,H). In panels A-C and E-H, dotted line represents average value found in tumor free mice. (I) Representative flow cytometry data of CD11b^+^/Ly6C^*hi/low*^/Ly6G^+*/*−^ cells found in WT, KO1, and TF groups. (J,K) Suppression of naïve CD8^+^ T cell proliferation by (J) freshly isolated G-MDSCs and (K) total splenocytes from WT, KO1, and TF groups (n = 3/group) on a per cell basis. ANOVA with post-hoc Tukey tests were used to assess statistical significance, where different letter (a, b, or c) denotes statistically different groups. (L) Cell trace profiles of live CD8^+^ T cells contained within CD45^+^ cells extracted from WT (red) and KO1 (blue) tumors in a time-matched experiment (TME) and stimulated for three days in vitro with *α*CD3/*α*CD28-loaded beads (representative of n = 3/group), with unstimulated cells as a staining control (gray).

To test whether CCN4-KO impairs splenic MDSC expansion independent of tumor burden, we analyzed MDSC in mice bearing CCN4-KO and YUMM1.7-WT tumors of similar size (size-matched, Fig. 4A, p-value=0.0843). Once again, significant decreases in the percentage (p-value=0.0006) and number per gram of spleen (p-value=0.0007) of MDSC were observed in CCN4-KO TB mice compared to YUMM1.7-WT counterparts (Fig. 4B-D). Given that CCN4-KO reduced MDSC in both time-matched and size-matched TB mice, we asked whether CCN4 had a differential effect on the two main MDSC subpopulations. Using a size-matched experimental design, CD11b^+^Ly6C^*low*^Ly6G^+^ granulocytic MDSC (G-MDSC) were the majority of the splenic MDSC in mice with YUMM1.7-WT tumors (Fig. 4E-I). Interestingly, a 2.7-fold and 2.9-fold reduction was observed in the percentage (Fig. 4E, p-value=0.0009) and the number (Fig. 4G, p-value=0.0003) of G-MDSC per gram of spleen, respectively. In addition, no differences in the frequency of splenic CD11b^+^Ly6C^*hi*^Ly6G^−^ monocytic MDSC (M-MDSC) were found in the absence of tumor-derived CCN4 (Fig. 4F and H). We also noted that G-MDSC (CXCR2^+^ CCR2^−^ CD11b^+^ GR1^+^) were predominantly captured in our initial MDSC gating based on GR1/CD11b staining (Fig. S6D).

Next, we tested whether splenic G-MDSC, isolated from CCN4-KO and YUMM1.7-WT TB mice, could suppress naïve CD8^+^ T cell proliferation stimulated by *α*CD3/*α*CD28-loaded beads (Fig. 4J). In a time-matched design, fresh G-MDSC induced by CCN4-KO tumors were significantly less suppressive on a per cell basis than those isolated from YUMM1.7-WT TB mice. In addition, CD11b^+^Ly6C^*low*^Ly6G^+^ cells isolated from tumor-free (TF) mice were unable to suppress CD8^+^ T cell proliferation. While G-MDSC were the predominant MDSC subset in TB mice (7% of splenocytes), we also evaluated CD8^+^ T cell proliferation after stimulating total splenocytes from TB mice with *α*CD3/*α*CD28-loaded beads (Fig. 4K). The precursor frequency of proliferative CD8^+^ T cells (KO1: 0.357± 0.022 versus WT: 0.220 ±0.011, p-value = 0.0007) and the fraction that divided at least once (KO1: 0.700 ±0.020 versus WT: 0.502± 0.028, p-value = 0.0006) were significantly increased when splenocytes from CCN4-KO TB mice were compared with splenocytes from mice with YUMM1.7-WT tumors. Once cell division occurred, CD8^+^ T splenocytes from CCN4-KO TB mice also had higher indices of proliferation (KO1: 1.72 ±0.02 versus WT: 1.52 ±0.05, p-value = 0.0040) and division (KO1: 0.61 ±0.03 versus WT: 0.33 ±0.03, p-value = 0.0002).

When tumor-infiltrating CD45^+^ cells were stimulated in similar fashion (Fig. 4L), live CD8^+^ T cells extracted from CCN4-KO and WT YUMM1.7 tumors had the same precursor frequency of proliferative cells (KO1: 0.884 ±0.051 versus WT: 0.884±0.029, p-value = 0.998) and fraction that divided at least once to in vitro stimulation (KO1: 0.985 ± 0.008 versus WT: 0.971 ± 0.009, p-value = 0.109). While CD8^+^ T cells were less prevalent in WT compared to CCN4-KO tumors, proliferation on a per cell basis was also reduced. Specifically, CD8^+^ T cells extracted from CCN4-KO tumors had higher indices of proliferation (KO1: 2.26 ± 0.08 versus WT: 1.63±0.08, p-value = 0.0006) and division (KO1: 2.00 ± 0.16 versus WT: 1.45 ± 0.11, p-value = 0.009). While we observed a similar effect on CD8^+^ T cell proliferation in WT versus CCN4 KO systems when using isolated G-MDSC as with splenocytes or TILs, the presence of T regulatory cells is a potential additional suppressive cell type in the mixed cell assays. We do note, though, that MDSC were 10-times more abundant than CD4^+^ T cells in analyzing TIL populations in WT YUMM1.7 tumors. Moreover, we observed no difference in T regulatory cell fraction (Live CD45+ CD4+ FOXP3+ CD25+ events) within the CD4+ TIL compartment upon CCN4 KO (p-value = 0.566, Fig. S7). Collectively, these results suggest that melanoma-derived CCN4 contributes to the splenic expansion of immunosuppressive G-MDSC.

### CCN4 stimulates the secretion of lactate and MDSC-attracting chemokines by melanoma cells and directly inhibits CD8^+^ T cells

To identify mechanisms associated with CCN4-mediated immunomodulation, we analyzed the cytokines, chemokines and growth factors produced by CCN4-KO and parental YUMM1.7-WT cells in vitro using R&D Systems Mouse XL Cytokine Array (Fig. 5A). Interestingly, CCL2 and CXCL1 chemokines were among the most down-regulated proteins in the tumor-conditioned media (TCM) from CCN4-KO tumor cells with high basal expression. These chemokines have been previously associated with MDSC recruitment to the tumor and their splenic accumulation (20–22). The reduction in CCL2 and CXCL1 were confirmed by ELISA in the TCM of CCN4-KO cells compared to YUMM1.7-WT counterparts (Fig. 5B and C, p-value=0.0001), as well as in media conditioned by CD45^−^ cells that were isolated from CCN4-KO and YUMM1.7WT tumors, after 36 h of culture ex vivo (Fig. 5D and E, p-value=0.0151 and p-value<0.0001, respectively). Additionally, CCL2 and CXCL1 were significantly diminished in the serum from CCN4-KO TB mice compared to mice with YUMM1.7-WT tumors (Fig. 5F and G). In fact, CCL2 and CXCL1 serum concentrations in CCN4-KO TB mice were not different from normal levels detected in tumor-free mice (Fig. 5F and G). We also observed that G-MDSC expressed a receptor for CXCL1: CXCR2 (Fig. S6D).

**Figure 5.**
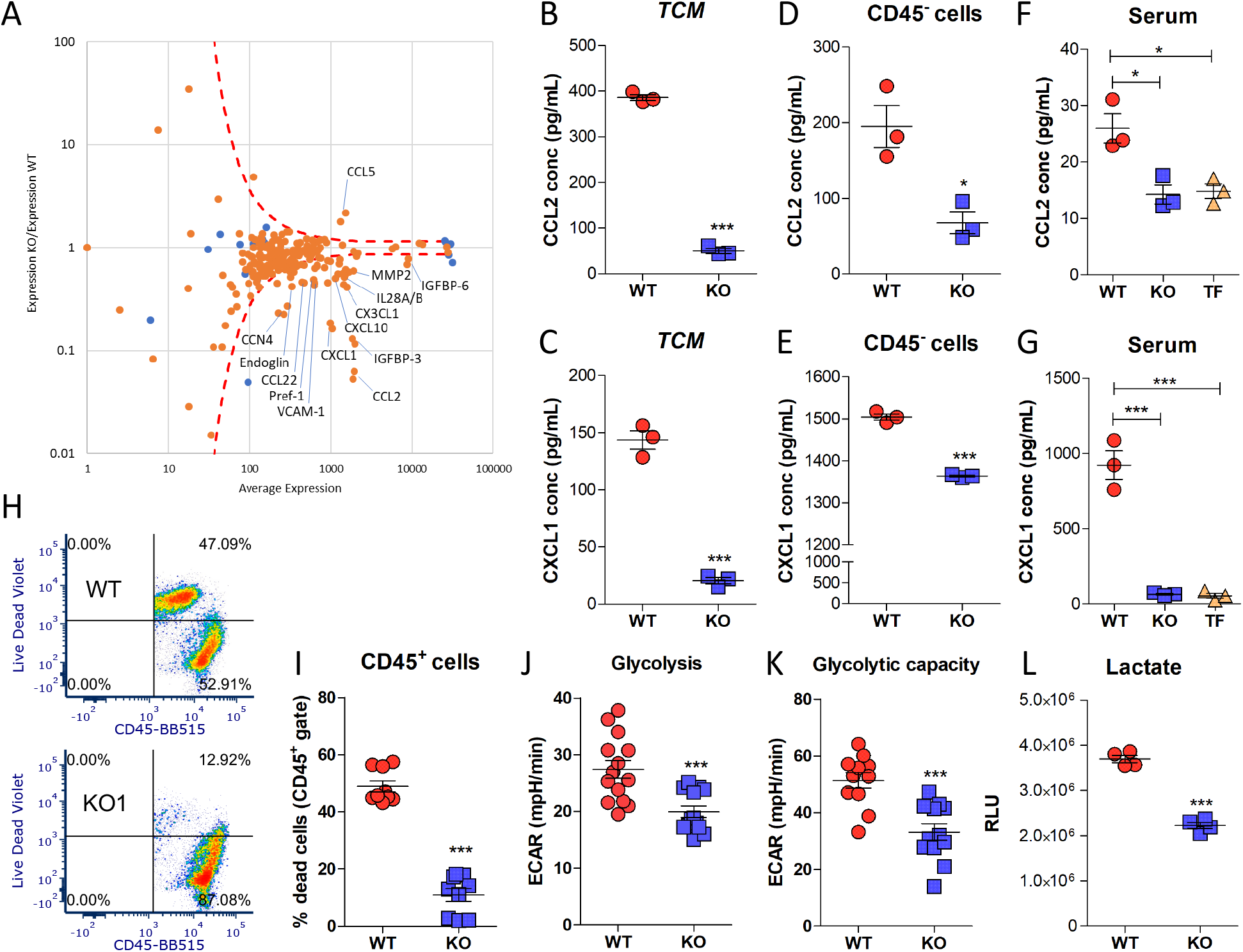
Knockout of CCN4 down-regulated CCL2 and CXCL1 expression and decreased glycolysis and glycolytic capacity in YUMM1.7 melanoma cells. (A) Cytokine, chemokine, and growth factor expression by YUMM1.7-WT and CCN4-KO cells in vitro assayed using R&D Systems’ Mouse XL Cytokine Array. Orange dots represent results for specific cytokine probes while blue dots represent positive and negative controls. Dotted lines enclose a null distribution estimated from positive and negative controls. (B-G) ELISA results for CCL2 and CXCL1 secretion after 36 h ex vivo culture (n = 3/group). CCL2 assayed in (B) tumor-conditioned medium (TCM), (D) CD45^−^ medium, and (F) serum. CXCL1 assayed in (E) TCM, (E) CD45^−^ medium, and (G) serum. (H-I) Live-dead staining of CD45^+^ cells isolated from WT and CCN4-KO tumors. (J-K) Analysis of the extracellular acidification rate (ECAR) associated with (J) glycolysis and (K) glycolytic capacity assayed by Seahorse Analyzer in WT and CCN4-KO tumors (n ≥ 12 WT, KO) after 36 hours ex vivo. (L) Lactate production by isolated CD45^−^ cells after 36 h ex vivo culture (n = 4).

IGFBP-3 (Insulin Like Growth Factor Binding Protein 3) was another protein showing a significant change between CCN4-KO and YUMM1.7-WT conditioned media with high basal expression (Fig. 5A). Considering that both IGFBP-3 and CCN4 play a role in glycolysis regulation (23–26) and that the viability of tumor-infiltrating CD45^+^ cells was reduced in YUMM1.7-WT tumors when compared to CCN4-KO counterparts (Fig. 5H and I, p-value <0.0001), we measured the extracellular acidification rate (ECAR) associated with glycolysis and glycolytic capacity in WT and CCN4-KO YUMM1.7 cells using a Seahorse Analyzer (Fig. 5J and K and Fig. S8A). Of note, glycolysis (p-value = 0.0006) and glycolytic capacity (p-value = 0.0002) were significantly reduced in the absence of CCN4. Similar reduction in glycolysis parameters in CCN4-KO cells was observed in the B16F0 tumor model (Fig. S8B-D).

One consequence of aerobic glycolysis in tumor cells is the release of lactate that acidifies the tumor microenvironment (27), suppresses or induces apoptosis of tumor-infiltrating lymphocytes (27, 28) and expands MDSC (29). Therefore, lactate production from CD45^−^ cells, isolated from CCN4-KO and YUMM1.7-WT tumors, was evaluated after 36 h of ex vivo culture. As shown in Fig 5L, CCN4-KO tumor cells secreted less lactate into the extracellular milieu (p-value <0.0001), which is consistent with reduced glycolysis and differences in TIL viability.

Given the observed increase in CD8^+^ T cells upon CCN4-KO, we next tested whether CCN4 had a direct impact on CD8^+^ T cell function through quantifying target-specific ex vivo cytokine release. To generate YUMM1.7-reactive CD8^+^ T cells, we immunized C57BL/6 mice using subcutaneous injection of irradiated YUMM1.7 cells and boosted with live YUMM1.7 cells three days before isolating CD8^+^ T cells. As target cells, we used the CCN4-inducible variant of CCN4 KO YUMM1.7 cells under the control of doxycycline and the corresponding vector control. IFN*γ* ELISpots were used to quantify the CD8^+^ T cell functional response to the different tumor targets in the absence or presence of tumor-produced CCN4 (Fig. 6A-C). As expected, the highest IFN*γ* and lowest TNF*α* responses were against WT and CCN4 KO YUMM1.7 cells, with a seemingly higher IFN*γ* response to WT YUMM1.7 targets (Fig. 6C, p-value = 0.098). Interestingly, re-expression of CCN4 by CCN4 KO YUMM1.7 cells following doxycycline induction significantly reduced both IFN*γ* and TNF*α* production (p-value < 0.001), which suggests that CCN4 directly inhibits CD8^+^ T cell function.

**Figure 6.**
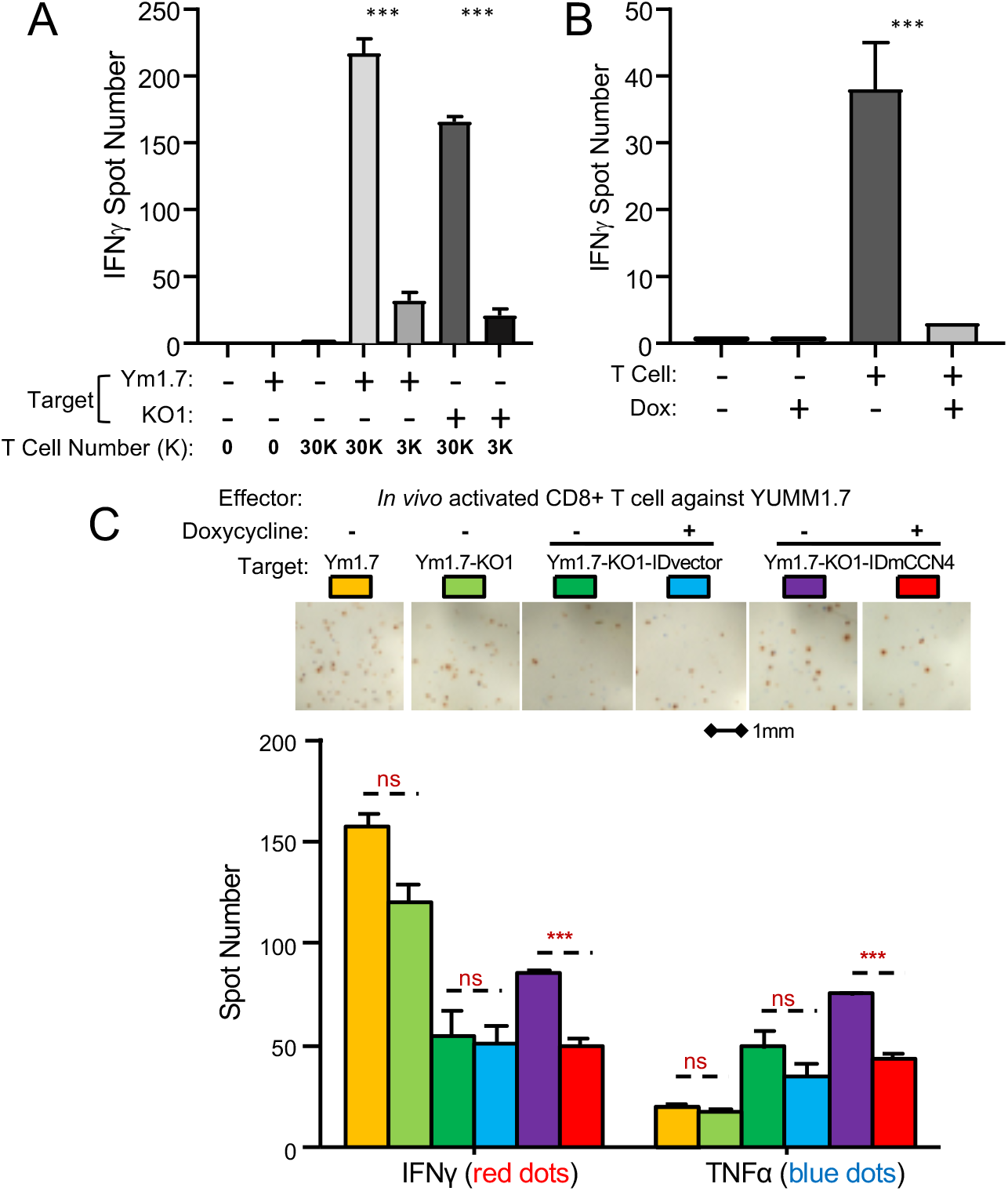
CCN4 directly inhibited CD8^+^ T cell function. (A) ELISpot for IFN*γ* release by CD8^+^ T cells using parental YUMM1.7 and CCN4-KO YUMM1.7 (KO1) cells as targets and different amount of in vivo activated CD8^+^ T cells. (B) ELISpot for IFN*γ* release by in vivo activated CD8^+^ T cells with CCN4-inducible cells as targets in the presence or absence of 0.5 mg/ml doxycycline. (C) CD8+ T cells isolated from the spleens of C57BL/6 mice that rejected YUMM1.7 tumors were assayed by in vitro ELISpot using variants of the YUMM1.7 cell line as targets (WT YUMM1.7 (Ym1.7) -yellow, CCN4-KO YUMM1.7 (Ym1.7-KO1)-light green, CCN4-KO YUMM1.7 with a blank inducible expression vector (Ym1.7-KO1-IDvector) -dark green and blue, CCN4-KO YUMM1.7 with a CCN4 inducible expression vector (Ym1.7-KO1-IDmCCN4) -purple and red). Variants containing the inducible expression vector were also cultured in the absence (dark green and purple) or presence of 0.5 *µ*g/ml doxycycline (blue and red). CD8+ T cells expressing IFN*γ* and TNF*α* were quantified following 24 hour coculture (bar graph). Results shown as mean ± S.D. for three biological replicates.

Taken together these results indicate that melanoma-derived CCN4 directly inhibits CD8^+^ T cell function but also has autocrine effects to produce CXCL1 and CCL2 and to stimulate glycolysis and lactate secretion by tumor cells. While cited studies describe the functional implications of these observations, our data suggest, to us, that the autocrine effects of CCN4 contribute to MDSC expansion and recruitment, as well as to the apoptosis of tumor-infiltrating CD45^+^ cells. A direct inhibitory effect of CCN4 on DC differentiation and maturation were not observed (Fig. S9). As CCN4 is expressed heterogeneously among human melanoma cells (Fig. 1B), additional work will be required to parse how autocrine versus paracrine effects propagate from CCN4-expressing to non-CCN4-expressing cells within the tumor microenvironment.

### CCN4 knockout complements the anti-tumor effect of immune checkpoint blockade therapy

As CCN4-KO increased T cell infiltration, we next compared the anti-tumor effect of an *α*PD1 antibody on CCN4-KO and YUMM1.7-WT tumors. Administering *α*PD1 and isotype control (IC) antibodies started when the tumors reached approximately 100 mm^3^ in size for all experimental groups. As shown in Fig. 7A, CCN4-KO delayed the beginning of *α*PD1 therapy for 11 days. Of note, a significant reduction in tumor growth was observed for both CCN4-KO and YUMM1.7-WT tumors after three doses of the *α*PD1 antibody when compared to mice receiving the IC antibody. Interestingly, mice with CCN4-KO tumors treated with *α*PD1 antibody had a tumor volume of 413.9 ± 94.99 mm^3^ on day 32, whereas YUMM1.7-WT TB mice receiving IC antibody reached a tumor volume of 1540 ± 300.7 mm^3^ on day 21, four days after the last antibody dose for each group. Similar results were obtained in the B16F0 model, where administering an *α*CTLA4 antibody significantly delayed the growth of CCN4-KO but not B16F0-WT tumors (Fig. S10).

**Figure 7.**
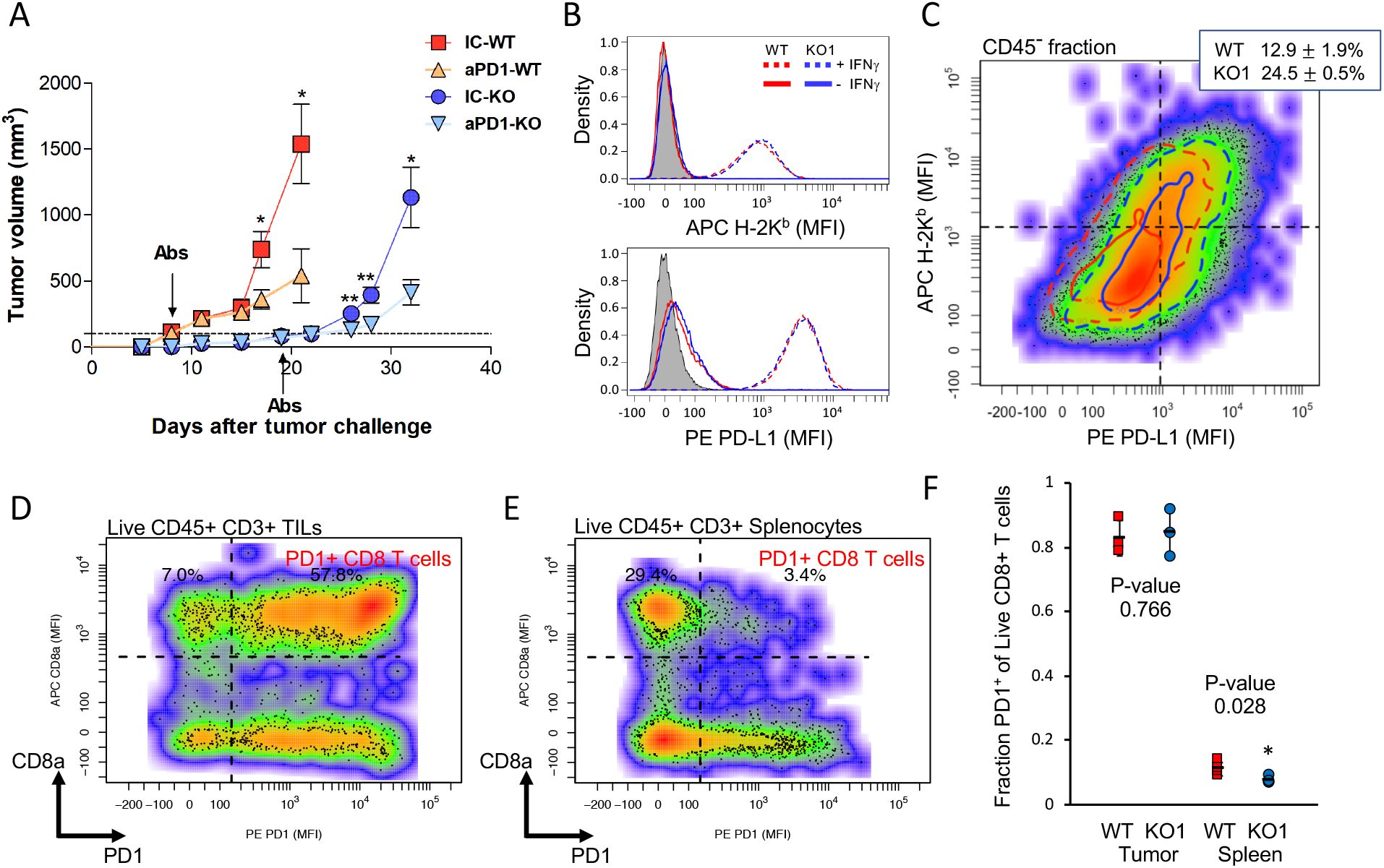
CCN4 knockout further promoted the anti-tumor effect of immune checkpoint blockade therapy. (A) Average tumor volumes of mice bearing YUMM1.7-WT (squares and triangles) or CCN4-KO (KO1, circles and inverted triangles) tumors (n = 4/group). Groups were treated with either *α*PD1 (triangles and inverted triangles) or isotype control (squares and circles) antibodies when the tumors reached 100 mm^3^. (B) Expression of H-2K^*b*^ (top panel) and PD-L1 (bottom panel) were assayed by flow cytometry in WT (red curves) and CCN4-KO (KO1 blue curves) YUMM1.7 cells with (dotted curves) and without (solid curves) preconditioning with IFN*γ*. Unstained cells were used as a negative control (shaded curve). (C) CD45^−^ fraction isolated from WT and CCN4-KO YUMM1.7 tumors in a time-matched experiment were assayed for H-2K^*b*^ and PD-L1 expression by flow cytometry. Contour curves enclose 90% (dotted curve) and 50% (solid curves) of CD45^−^ events obtained from WT (red) and CCN4-KO (blue) YUMM1.7 tumors. CD8+ T cells expressing PD1 within the tumor (D) and spleen (E) were assayed by flow cytometry in mice bearing WT and CCN4-KO YUMM1.7 tumors (F). Results representative of at least three biological replicates.

While generally the effect of CCN4 KO was consistent between B16F0 and YUMM1.7 models, subtle differences in immunosuppression mechanisms seemed to change the susceptibility of these two systems to treatment with a specific immune checkpoint. For instance in ELISpot assays, we found that CCN4 inhibited antigen-specific release by CD8^+^ T cells of IFN*γ*. Separately, we observed that IFN*γ* upregulated the MHC class I molecule H-2K^*b*^ and PD-L1 (CD274) in both WT and CCN4-KO YUMM1.7 cells (Fig. 7B). In assaying these two proteins on the CD45^−^ fraction isolated from WT and CCN4-KO YUMM1.7 tumors (Fig. 7C), cells with increased expression of both H-2K^*b*^ and PD-L1 increased from 12.9 ± 1.9% in WT to 24.5 ± 0.5% in CCN4-KO tumors (n = 3, p-value = 0.0005). While tumor-infiltrating CD8+ T cells predominantly expressed PD1 compared to splenocytes (Fig. 7D-F), we observed no difference in PD1^+^ CD8^+^ TILs upon CCN4 KO (p-value = 0.766, Fig. 7F). It follows then that, by knocking out CCN4, the PD1-PDL1 axis plays a more dominant role in the YUMM1.7 model, likely through IFN*γ* cross-talk, in restraining the T cell response and is more sensitive to therapeutic intervention. We also noted that infiltrating T cells and NK cells were almost 10-times lower per gram tumor in the WT B16F0 model compared with the WT YUMM1.7 model. Those differences coupled with B16F0 sensitivity to *α*CTLA4 suggest that pathways associated with initiating an anti-tumor response, such as the extent of clonal T cell expansion within secondary lymphoid organs, are key limiters in the B16F0 model. Overall, these data indicate that absence of tumor-derived CCN4 complements the anti-tumor effect of immune checkpoint blockade therapy.

## Discussion

Recent literature connects the epithelial-mesenchymal transition (EMT) with tumor immune escape (22, 30, 31). Interestingly, CCN4 activates EMT-associated genes in melanoma cells thus increasing metastatic potential (11, 12). Motivated by encouraging clinical correlates, we report that melanoma-derived CCN4 also stimulated tumor-induced immunosuppression in mice, particularly by directly suppressing antigen-induced IFN*γ* release by CD8^+^ T cells and by expanding and recruiting G-MDSC. IFN*γ* action within the tumor microenvironment is a key promoter of anti-tumor immunity (32). In addition, tumor-induced MDSC suppress the proliferation and effector function of T cells and NK cells and interfere with the migration and viability of lymphocytes (33). Correspondingly, increased anti-tumor T cells (CD4^+^ and CD8^+^) and NK cells were observed in CCN4-KO tumors, concomitant with a reduced frequency of MDSC. Moreover, the ability of G-MDSC to suppress T cell proliferation was diminished in mice bearing CCN4-KO tumors. Similar shifts in immune contexture with *CCN4* expression were observed in human data. While CCN4 is a downstream effector of the Wnt/*β*-catenin pathway, no significant difference in tumor-infiltrating DC was observed. This is in contrast to previous work from Gajewski et al. where deficient recruitment of CD103^+^ DC in melanoma tumors with intrinsic *β*-catenin signaling was associated with T cell exclusion (6). Of note, *Ccn4* was not significantly increased in tumors with a constitutively active *β*-catenin compared with Braf^*V* 600*E*^/Pten^−*/*−^ control mice based on transcriptional profiling using Illumina microarrays, which suggests that transcriptional co-activators play a role here in shaping cell-to-cell communication.

MDSC generation is a complex process mediated by tumor-derived soluble factors (34). Glycolysis exacerbation expands MDSCs, where a high glycolytic rate promotes MDSC generation by increasing granulocyte-macrophage colony-stimulating factor (GM-CSF) and granulocyte-colony stimulating factor (G-CSF) in triple negative breast cancer models (35). Interestingly, the glycolytic rate in CCN4-KO melanoma cells was reduced. Prior work notes that CCN4 promotes glycolysis in laryngeal squamous cell carcinoma (24). CCN4 also interacts with peroxisome proliferator-activated receptor (PPAR*γ*) to inhibit PPAR*γ* activity (23). In turn, PPAR*γ* stimulates adipocyte differentiation (23) and inhibits glycolysis (36). Thus, a model where CCN4 stimulates glycolysis by repressing PPAR*γ* could apply to our results with melanoma cells. However, we found no differences in GM-CSF and G-CSF secretion in the TCM from CCN4-KO and YUMM1.7-WT cells. Conversely, and associated with impaired glycolysis, lactate secretion was reduced in the absence of CCN4 in melanoma cells. Of note, Husain et al. observed a role for tumor-derived lactate in generating MDSC (29), suggesting that the reduced MDSC frequency in CCN4KO TB mice is related to a diminished lactate secretion in the tumor microenvironment. Lactate release by the tumor cells also acidifies the tumor microenvironment (27), which reinforces tumor-induced immunosuppression by impairing CD8+ T cell proliferation, cytokine production, and lytic activity (27, 28). In addition, lymphocytes apoptose at the low pH values of the tumor microenvironment (37). Correspondingly, we observed that half of CD45^+^ leukocytes in the tumor microenvironment of YUMM1.7-WT tumors were dead compared with around 90% viability in CCN4-KO counterparts.

CCL2 and CXCL1 secretion was also significantly reduced by CCN4-KO YUMM1.7 cells in vitro, as well as in the serum from TB mice and when analyzing the CD45^−^ cells isolated from the s.c. tumors. Of note, CCL2-CCR2 signaling plays a key role in recruiting MDSC into the tumor microenvironment in glioma, renal and colon carcinomas, lung cancer, and melanoma (38–43). CXCL1 interaction with CXCR2 also helps recruit G-MDSC into ovarian cancer microenvironment, where a high CXCL1 serum concentration correlates with increased tumor-infiltrating G-MDSC and a poor prognosis (22). Interestingly, this process is mediated by Snail (SNAI1), a relevant transcription repressor regulating EMT, which stimulates CXCL1 gene expression through a direct binding to the promoter and via NF-*κ*B activation (22). Additionally, Snail regulates CCL2 production in epithelial cells (44). Of note, CCN4 activates Snail in YUMM1.7 melanoma cells and Snail overexpression in CCN4-KO cells rescues the metastatic potential of B16F10 cells in vivo (11). While additional experiments would help clarify the conditional dependence among these changes in immune contexture elicited by CCN4, our results are consistent with a model where CCN4-induced Snail activation promotes CXCL1 and CCL2 secretion directly or through NF-*κ*B activation, which in turn stimulates G-MDSC expansion and suppressive function.

Antibodies targeting CTLA4 (ipilimumab) and PD1 (nivolumab and pembrolizumab) have proven the potential of ICB immunotherapy in patients with different malignancies, like melanoma. However, even when the response rate in melanoma is high compared to other treatments, many patients still do not receive clinical benefit (45–47). Since ICB treatment relieves inhibitory signals controlling T cell function, combination therapies that increase T cell infiltration in the tumor can improve the anti-tumor effect of ICB. Our results demonstrated that knocking out CCN4 in the tumor cells complemented the anti-tumor effect of anti-PD1 and anti-CTLA4 antibodies in YUMM1.7 and B16F0 mouse melanoma models, respectively. Thus, targeting CCN4 can enhance ICB therapy not only by increasing T cell infiltration in the tumor and enhancing local IFN*γ* production, but also through ameliorating MDSC-mediated suppression. Others have shown that targeting MDSCs, for example by blocking CCL2-CCR2 interactions, enhances the anti-tumor effect of ICB therapy even in resistant tumors like glioblastoma (48). Considering these elements, targeting CCN4 is immunotherapeutic strategy that can potentially impair tumor-induced immunosuppression and enhance ICB therapy while inhibiting EMT and metastasis formation.

## Supporting information

PeerReview_CancerResearch

## ACKNOWLEDGEMENTS

This work was supported by National Science Foundation (NSF CBET-1644932 to DJK) and National Cancer Institute (NCI 1R01CA193473 to DJK). The content is solely the responsibility of the authors and does not necessarily represent the official views of the NSF or NCI. We also used equipment from the WVU Flow Cytometry & Single Cell core, which was supported by the National Institutes of Health Grants GM103488/RR032138, GM104942, GM103434, and OD016165.

## AUTHOR CONTRIBUTIONS

Conception and design: AF and DJK; Data Acquisition: AF, WD, SLM, and SR; Analysis and interpretation of data: AF, WD, AP, SR, and DJK; Funding acquisition: DJK; Methodology: AF and DJK; Project administration: DJK; Software: DJK; Supervision: DJK; Writing -original draft: AF, AP, and DJK; Writing -review & editing: all authors.

## Supplementary Information

**Fig. S1.**
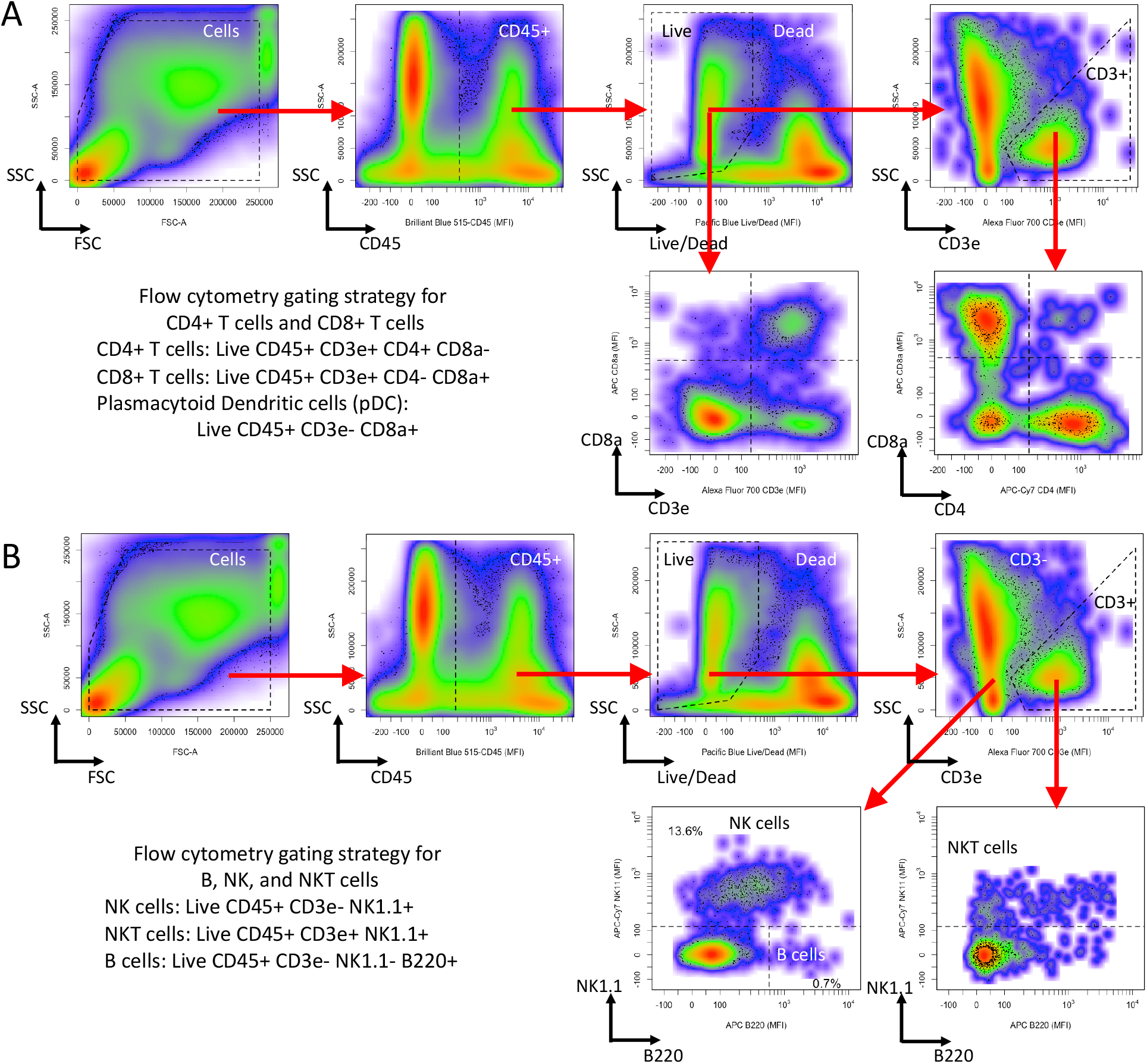
Flow cytometry gating strategy for T cells (A) and B, NK, and NKT cells (B). (A) CD45 staining versus side scatter area was used to gate for CD45+ cells. Live Dead Pacific Blue staining versus side scatter area was used to then gate for Live CD45^+^ cells, which were then gated based on CD3e^+^ expression. Live CD45^+^ CD3e^+^ cells were further subdivided into CD8^+^ T cells (live CD8a^+^ CD4^−^ CD3e^+^ CD45^+^ cells), CD4 T cells (live CD4^+^ CD8a^−^ CD3e^+^ CD45^+^ cells), plasmacytoid dendritic cells (pDC: live CD8a^+^ CD3e^−^ CD45^+^ cells), and double negative T cells (live CD8a^−^ CD4^−^ CD3e^+^ CD45^+^ cells). (B) CD45 staining versus side scatter area was used to gate for CD45+ cells. Live Dead Pacific Blue staining versus side scatter area was used to gate for Live CD45^+^ cells, which were then subdivided into B cells (live NK1.1^−^ B220^+^ CD3e^−^ CD45^+^ cells), NK cells (live NK1.1^+^ CD3e^−^ CD45^+^ cells), and NKT cells (live NK1.1^+^ CD3e^+^ CD45^+^ cells).

**Fig. S2.**
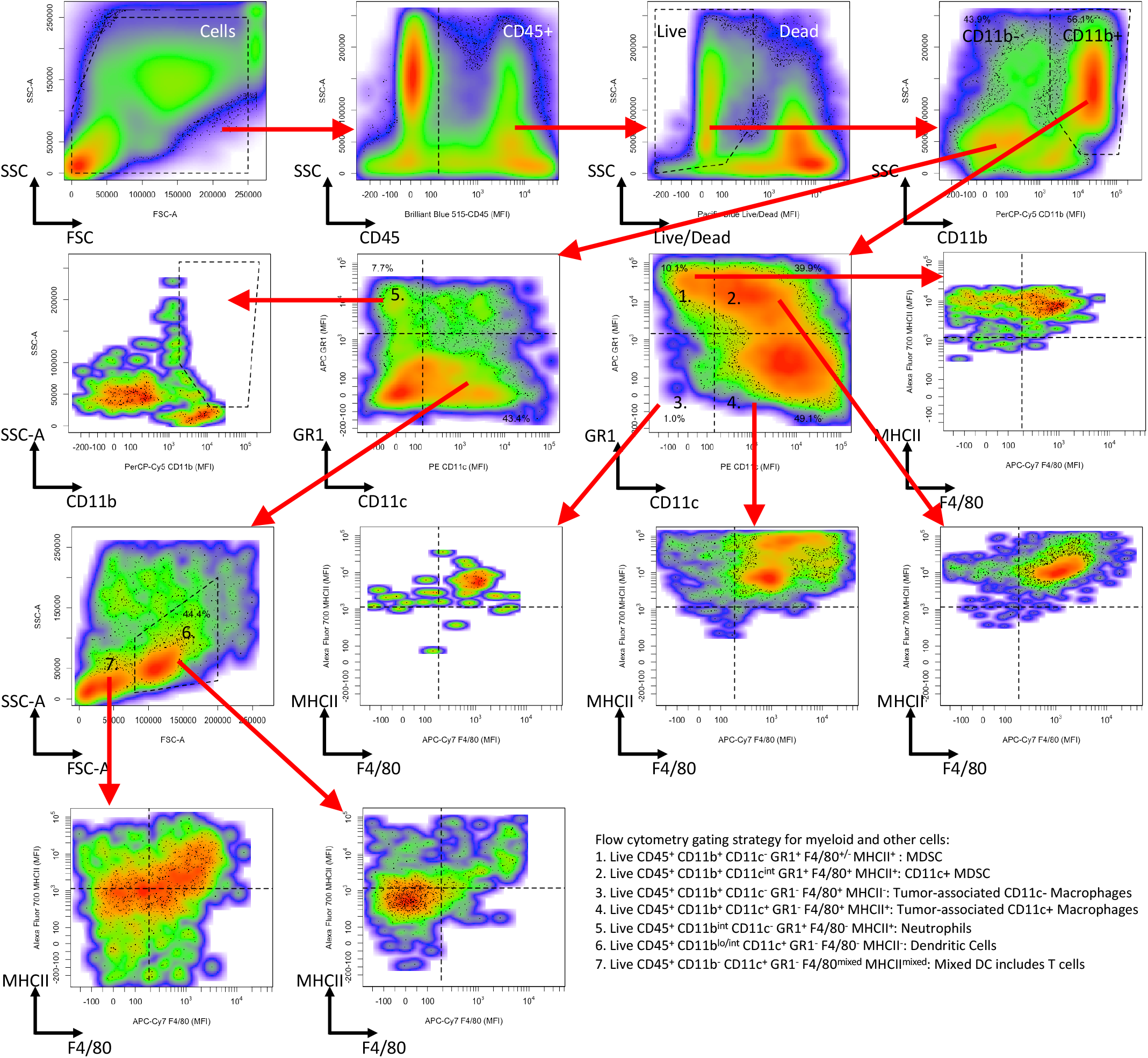
Flow cytometry gating strategy for Tumor associated neutrophils (TAN) and myeloid cell subsets. CD45 staining versus side scatter area was used to gate for CD45^+^ cells. Live Dead Pacific Blue staining versus side scatter area was used to gate for Live CD45^+^ cells, which were then subdivided into subsets based on CD11b staining followed by Gr1 versus CD11c staining. From the CD11b^+^ gate, myeloid-derived suppressor cells (MDSC) (live CD45^+^ CD11b^+^ Gr1^+^ cells) were subdivided into CD11c^*int/*+^ MDSC (F4/80^+^ MHCII^+^) and CD11c^−^ MDSC (F4/80^*mixed*^ MHCII^+^). Also from the CD11b^+^ gate, macrophages (live Gr1^−^ F4/80^+^ CD11b^+^ CD45^+^ cells) were subdivided into tumor-associated CD11c^+^ (CD11c^*int/*+^ MHCII^*hi*^) and CD11c^−^ (CD11c^−^ MHCII^*lo*^)subsets. The CD11b^−^ subset included tumor-associated neutrophils (TAN) (Gr1^+^ CD11c^−^ CD11b^*int*^ MHCII^*hi*^ F4/80^−^) and dendritic cells (Gr1^−^ CD11c^+^ CD11b^*lo/int*^ FSC-A^*hi*^ MHCII^*lo*^ F4/80^−^).

**Fig. S3.**
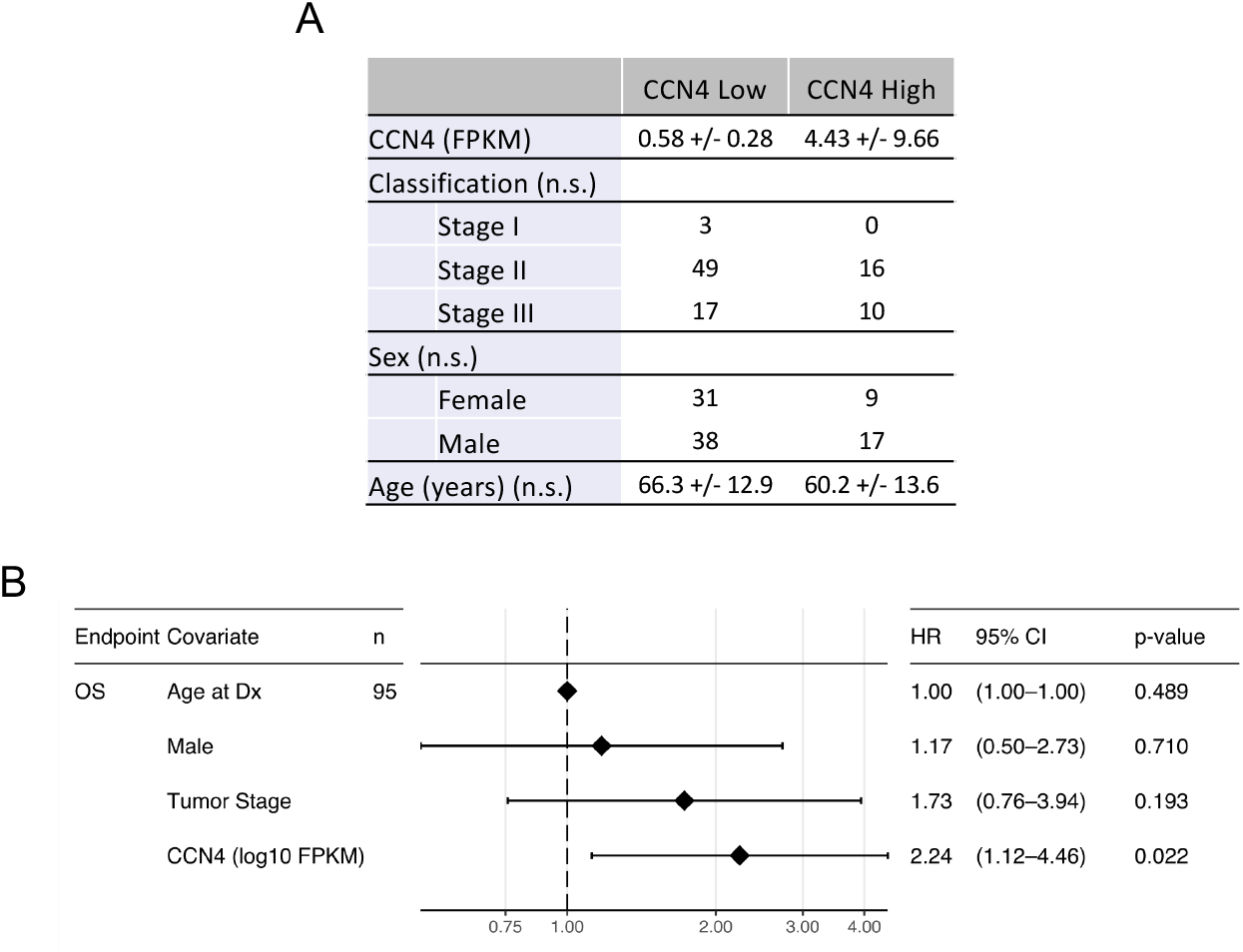
*CCN4* expression is associated with an increased hazard ratio for the overall survival of patients diagnosed with primary melanoma. (A) The characteristics of the patient population stratified by *CCN4* expression (*CCN4* high versus *CCN4* low). Statistical differences among categorical data and age at diagnosis were assessed using Fisher’s exact test and Student’s t test, respectively (n.s., p > 0.05). (B) Graphical summary of a multivariate Cox proportional hazards model for overall survival (OS) that was regressed to the indicated population covariates. TCGA data accessed 09/19/2019.

**Fig. S4.**
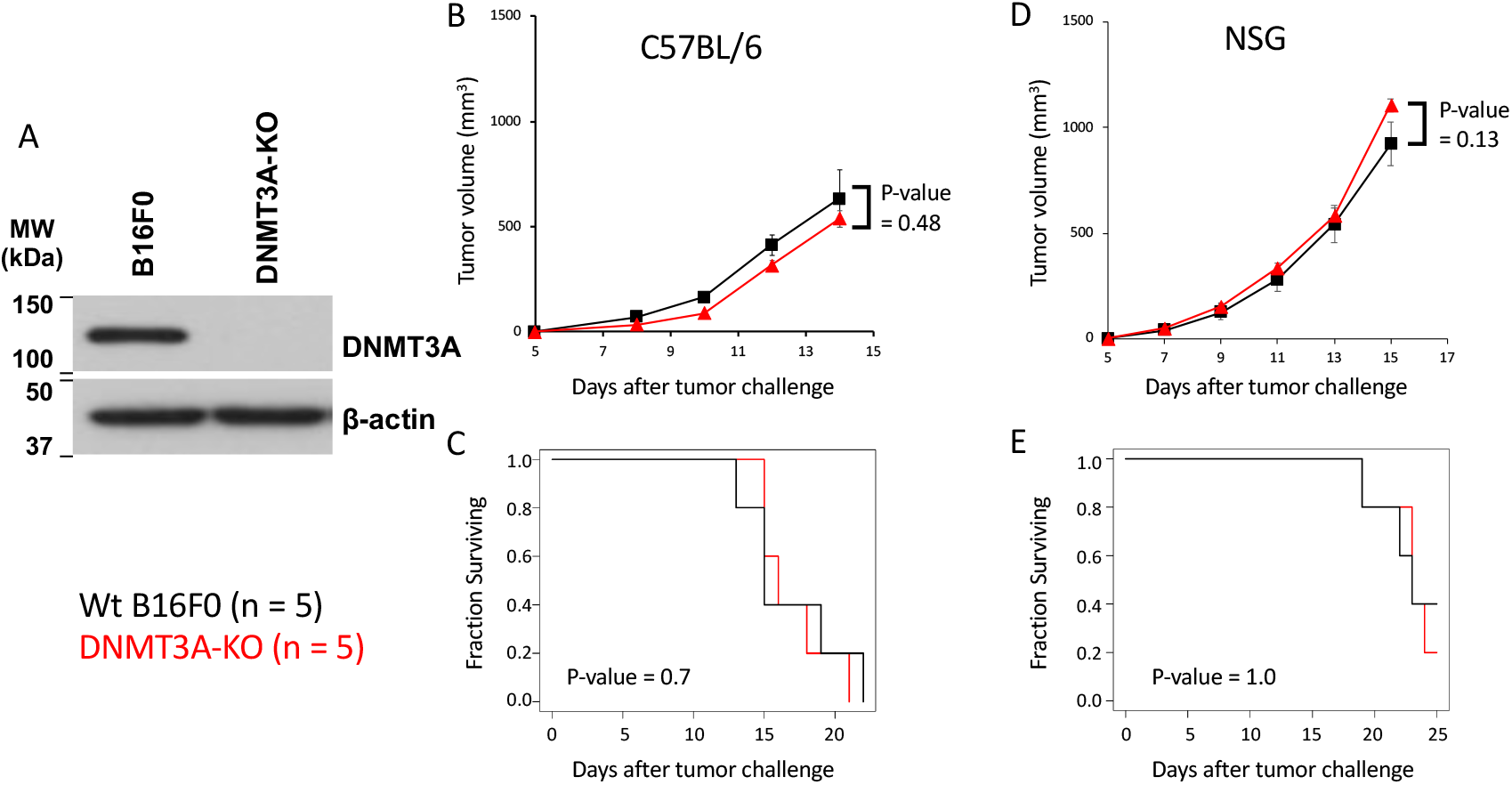
DNMT3A knockout had no significant effect on B16F0 melanoma growth in C57BL/6Ncrl and NSG mice. DNMT3A-KO variants of the B16F0 cell line were obtained through transfection with a mix of CRISPR/Cas9 KO and Homology-Directed Repair(HDR) plasmids (Santa Cruz BT SC-420035-NIC, SC-420035-NIC2), followed by puromycin selection. (A) Knockout of DNMT3A was confirmed by western blot of whole cell lysates (DNMT3A rabbit mAb Cell Signaling cat 3598) with beta-actin serving as loading control (SCBT mouse mAb clone C4). C57BL/6Ncrl (B,C) and NSG (D,E) mice were challenged with a subcutaneous injection of 3e5 WT B16F0 (black) and DNMT3A-KO variants cells (red) (n = 5/group). Tumor size (B, D), which was measured by calipers (mean +/-SEM), and overall survival (C,E) were used as outcome measures. Statistical significance associated with differences in tumor size was assessed using a two-sided homoscedastic Student’s t-test while differences in overall survival were assessed using the Peto and Peto modification of a Gehan-Wilcoxon test.

**Fig. S5.**
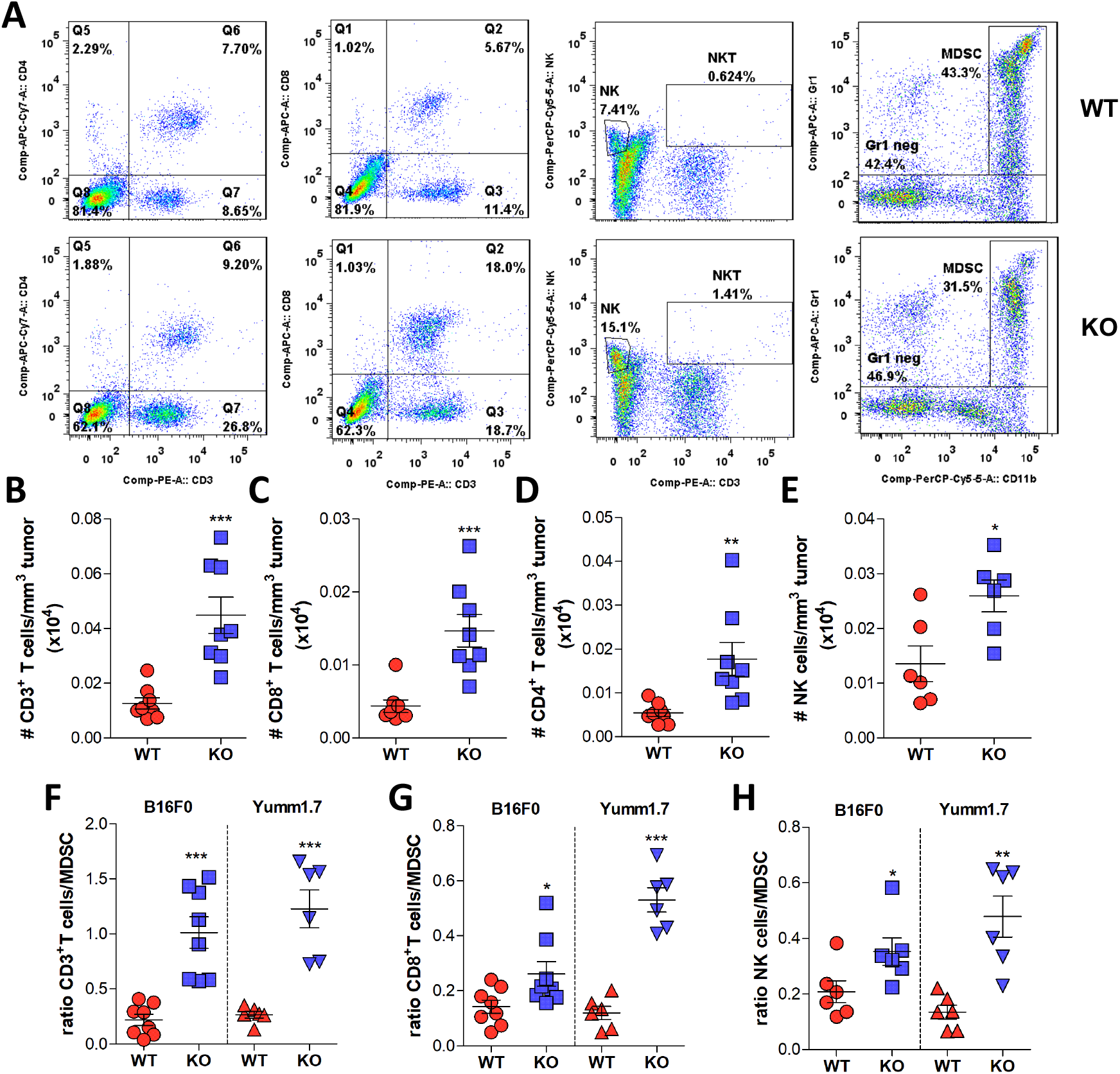
Comparison of TILs in mice receiving wt versus CCN4 KO B16F0 cells. (A) Representative flow cytometry panels comparing CD4^+^ T cells (Live CD45^+^ CD3^+^ CD8^−^ CD4^+^ events), CD8^+^ T cells (Live CD45^+^ CD3^+^ CD8^+^ CD4^−^ events), NK (Live CD45^+^ CD3^−^ NK1.1^+^ events) and NKT (Live CD45^+^ CD3^+^ NK1.1^+^ events) cells, and MDSC (Live CD45^+^ CD11b^+^ Gr1^+^ events) infiltration in tumors derived from WT and CCN4 KO B16F0 cells (left to right columns). (B-E) Summary figures for the number of CD3^+^ (B), CD8^+^ (C), and CD4^+^ (D) T cells and NK cells (E) per mm^3^ tumor (n ≥ 5/group). (F-H) Comparison in the change of CD3^+^ T cell:MDSC (F), CD8^+^ T cell:MDSC (G), NK cell:MDSC (H) ratios upon CCN4 KO between the B16F0 and YUMM1.7 models. * 0.01<p<0.05, ** 0.001<p<0.01, *** p<0.001. Error bars represent SEM.

**Fig. S6.**
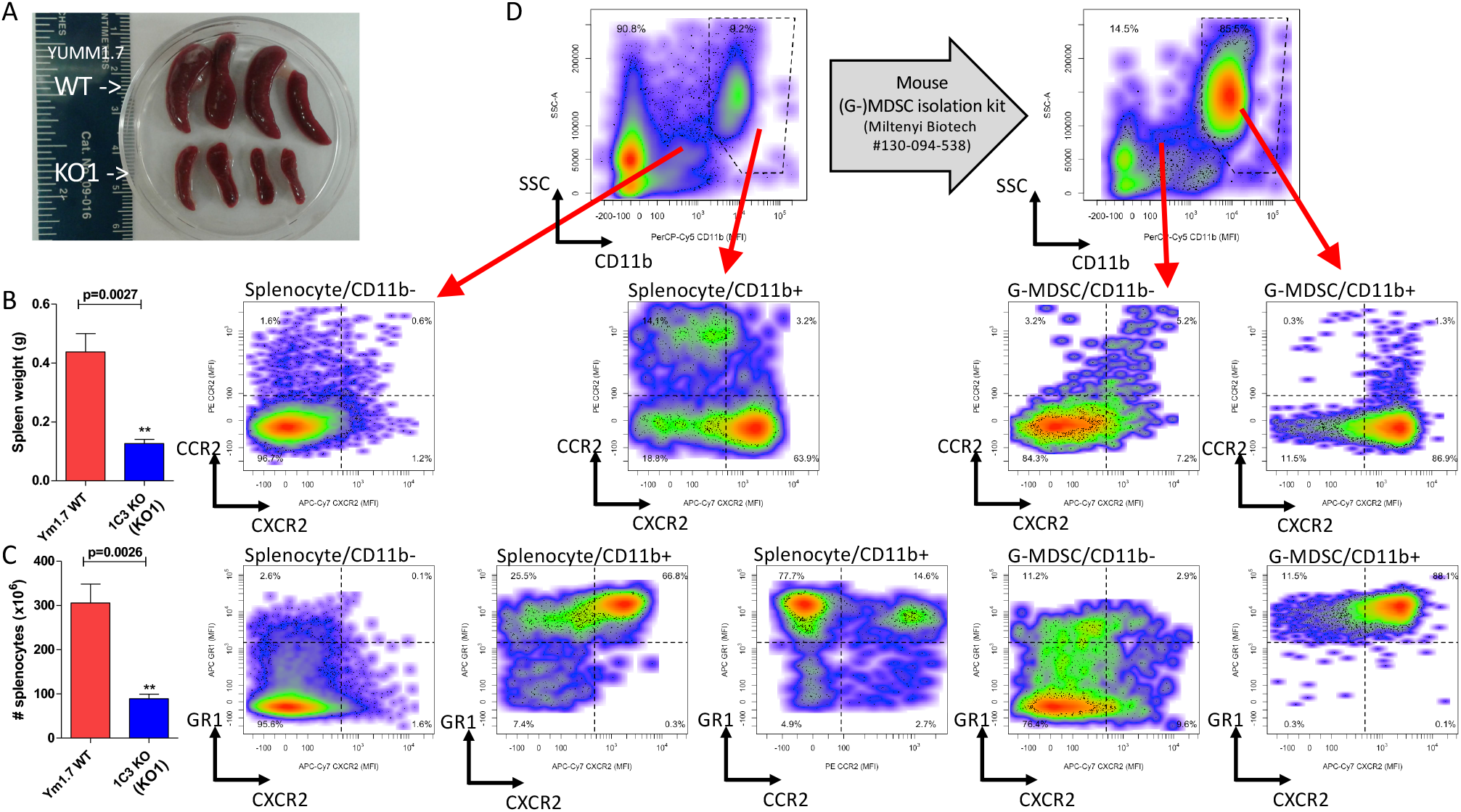
The presence of MDSCs in the spleens of mice bearing WT YUMM1.7 tumors was associated with splenomegaly. In a time-matched experiment, the spleens from mice bearing WT YUMM1.7 tumors were larger (A,B) and contained more splenocytes (C) than mice bearing YUMM1.7-KO (KO1) tumors. Error bars represent SEM. (D) To validate the MDSC gating strategy, we characterized splenocytes (left panels) and G-MDSC isolated from splenocytes (right panels, mouse G-MDSC isolation kit, Miltenyi Biotec) by flow cytometry using anti-mouse antibodies GR1 APC (Clone: RB6-8C5, BioLegend), CD11b PerCP-Cy5 (Clone: M1/70, ThermoFisher), CXCR2 APC-Cy7 (Clone: REA942, Miltenyi Biotec), and CCR2 PE (Clone: REA538, Miltenyi Biotec).

**Fig. S7.**
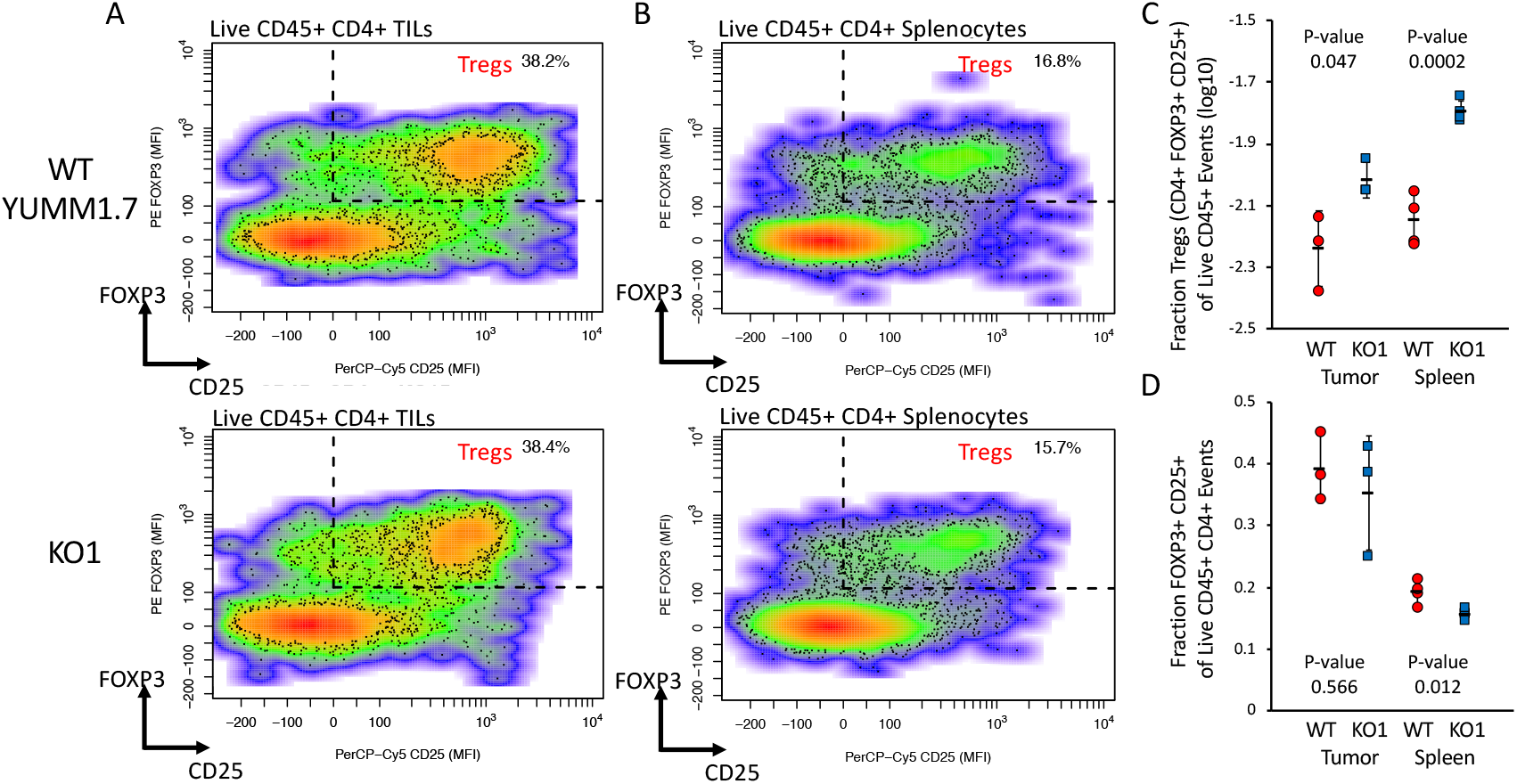
The fraction of T regulatory cells (Live CD45+ CD4+ FOXP3+ CD25+) within the CD4+ TIL compartment was not changed upon CCN4 KO. In a time-matched experiment, TILs (A) and splenocytes (B) obtained from mice bearing WT (n = 3, top panels) and CCN4 KO (n = 4, KO1 -bottom panels) YUMM1.7 tumors were analyzed for the presence of T regulatory cells (Tregs) by flow cytometry. Results are summarized in terms of the fraction of Tregs within total live CD45+ events (C) and the fraction of Tregs within the total live CD45+ CD4+ events (D). Statistical significance was assessed using a Student’s t-test and error bars represent the standard deviation.

**Fig. S8.**
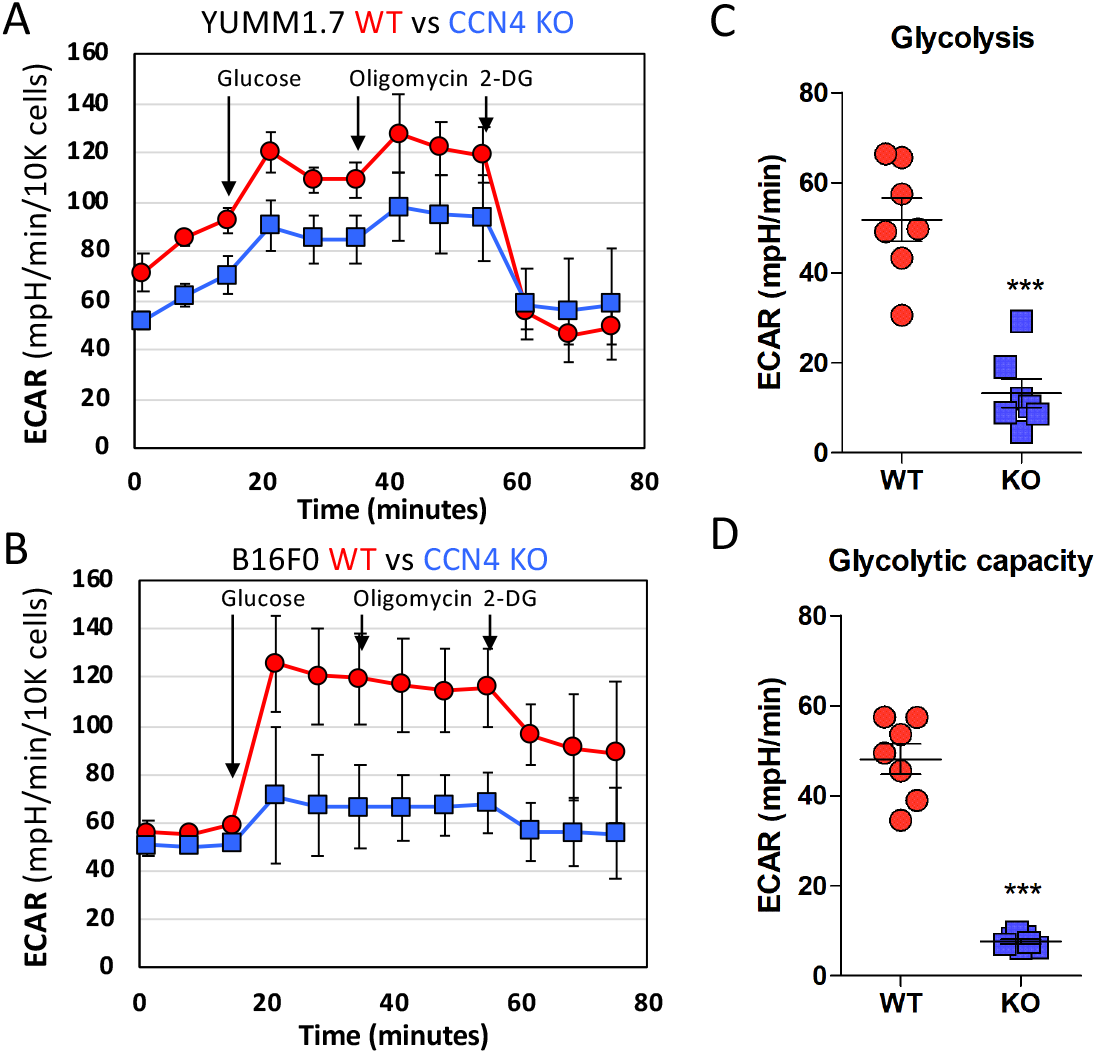
Glycolysis and glycolytic capacity in YUMM1.7 and B16F0 cells were reduced upon CCN4 KO. Time course of ECAR analysis using XFe96 Seahorse Analyzer in WT and CCN4-KO tumors derived from YUMM1.7 (A) and B16F0 cells (B) after 36 hours ex vivo. (n = 3, results representative of one of three independent experiments). Curves represent average *±* standard deviation. Time course profiles were analyzed to estimate (C) glycolysis and (D) glycolytic capacity for WT and CCN4-KO tumors derived from B16F0 cell variants. *** p<0.001. Error bars represent SEM.

**Fig. S9.**
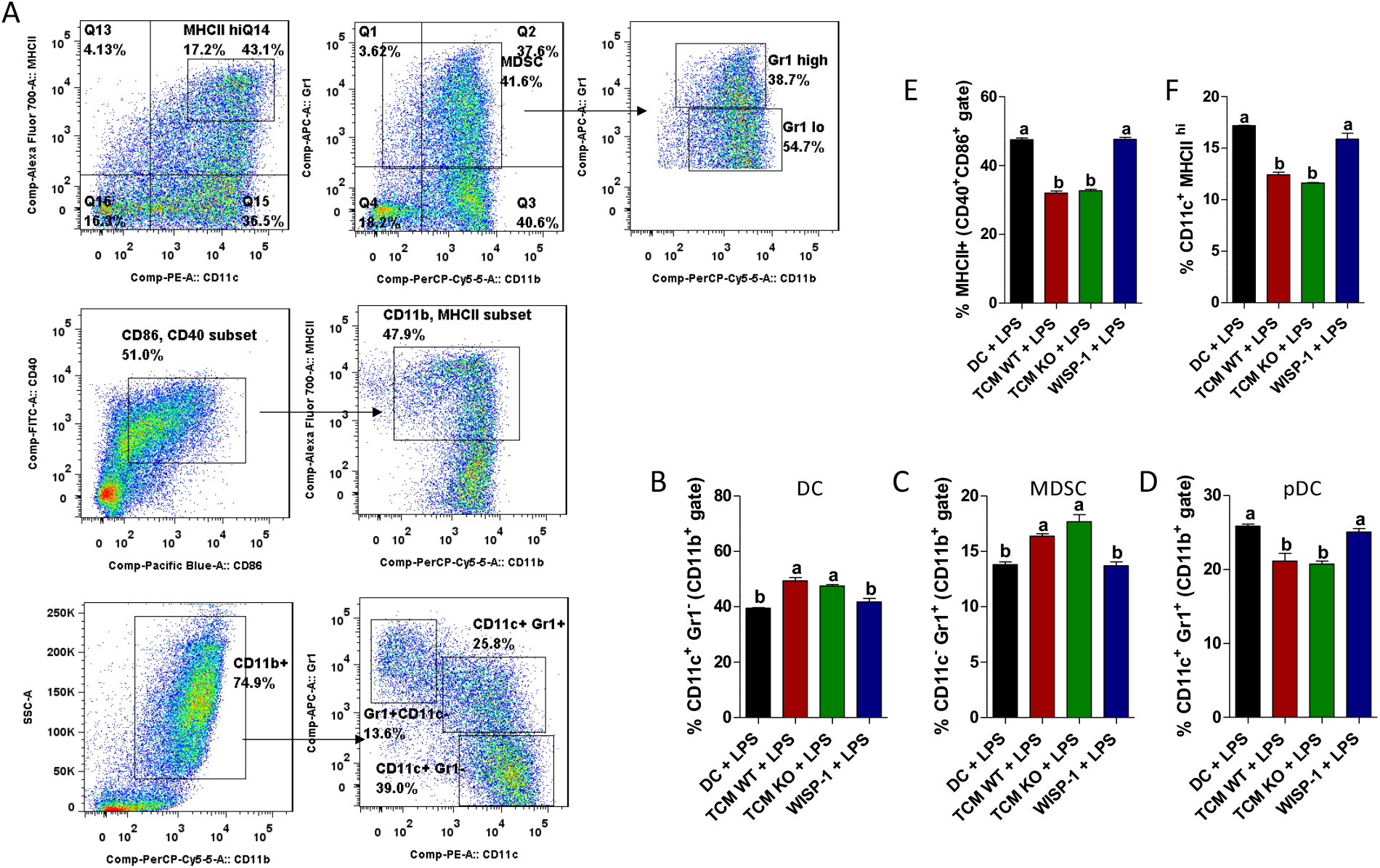
CCN4 has no direct inhibitory effect on dendritic cell maturation and differentiation. Bone marrow cells were harvested from femurs and tibias of C57BL/6 mice. For dendritic cell (DC) preparation, 6 × 10^5^ cells/well were cultured for 6 days with 20 ng/mL of GM-CSF (eBioscience, ThermoFisher) in 6 well plates. On day 6, the media was removed and DC maturation was induced using 1 *µ*g/mL of LPS (Sigma). Medium conditioned by WT (TCM WT) and CCN4/WISP1 KO (TCM KO) melanoma B16F0 cells was added at 50% final volume during differentiation and maturation of the DC, whereas rmCCN4 (WISP1, R&D) was added at a final concentration of 10 ng/mL. After 24h of LPS treatment, DC were extracted, washed, and incubated with Mouse BD Fc Block (BD Biosciences). Using the gating strategy illustrated in (A), the following antibodies were used to analyze by flow cytometry the efficiency of the DC generation (B-D) and the maturation of these cells (E-F): anti-mouse CD11c/PE (eBioscience, ThermoFisher), anti-mouse IA/IE (MHCII)/AlexaFluor 700 (BioLegend), anti-mouse CD11b/PerCP-Cy5.5 (eBioscience, ThermoFisher), anti-mouse Gr1/APC (BioLegend), anti-mouse CD40/FITC (eBioscience, ThermoFisher), and anti-mouse CD86/V450 (BD Biosciences). b indicates p<0.01 assessed by ANOVA with Tukey’s ad hoc post-test. Error bars represent SEM.

**Fig. S10.**
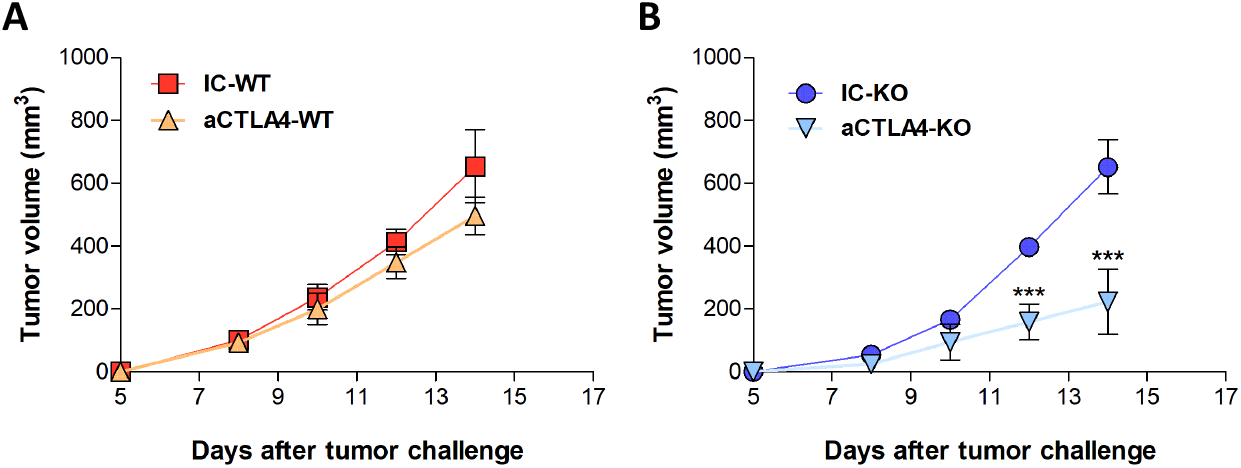
Tumor growth profiles of tumors derived from wt and CCN4 KO B16F0 cells in response to *α*CTLA4 mAb treatment. Average tumor volumes of C57BL/6 mice bearing B16F0-WT (squares and triangles) or CCN4-KO (circles and inverted triangles) tumors inoculated with s.c. injection of 3 × 10^5^ cells. Groups were treated with either *α*CTLA4 (triangles and inverted triangles) or isotype control (IC: squares and circles). The anti-CTLA4 and IC antibodies were administered intraperitoneally (i.p.) at a dose of 200 *µ*g/mouse on days 3, 7 and 10 following inoculation (n = 8/group, combined from two independent experiments). *** p<0.001. Error bars represent SEM.

## References

1. P. Darvin, S. M. Toor, V. Sasidharan Nair, and E. Elkord. Immune checkpoint inhibitors: recent progress and potential biomarkers. Exp. Mol. Med., 50(12):1–11, 12 2018.

2. M. Binnewies, E. W. Roberts, K. Kersten, V. Chan, D. F. Fearon, M. Merad, L. M. Coussens, D. I. Gabrilovich, S. Ostrand-Rosenberg, C. C. Hedrick, R. H. Vonderheide, M. J. Pittet, R. K. Jain, W. Zou, T. K. Howcroft, E. C. Woodhouse, R. A. Weinberg, and M. F. Krummel. Understanding the tumor immune microenvironment (TIME) for effective therapy. Nat. Med., 24(5):541–550, May 2018.

3. M. F. Fransen, M. Schoonderwoerd, P. Knopf, M. G. Camps, L. J. Hawinkels, M. Kneilling, T. van Hall, and F. Ossendorp. Tumor-draining lymph nodes are pivotal in PD-1/PD-L1 checkpoint therapy. JCI Insight, 3(23), Dec 2018.

4. A. I. Daud, K. Loo, M. L. Pauli, R. Sanchez-Rodriguez, P. M. Sandoval, K. Taravati, K. Tsai, A. Nosrati, L. Nardo, M. D. Alvarado, A. P. Algazi, M. H. Pampaloni, I. V. Lobach, J. Hwang, R. H. Pierce, I. K. Gratz, M. F. Krummel, and M. D. Rosenblum. Tumor immune profiling predicts response to anti-PD-1 therapy in human melanoma. J. Clin. Invest., 126(9):3447– 3452, 09 2016.

5. O. Hamid, H. Schmidt, A. Nissan, L. Ridolfi, S. Aamdal, J. Hansson, M. Guida, D. M. Hyams, H. Gómez, L. Bastholt, S. D. Chasalow, and D. Berman. A prospective phase II trial exploring the association between tumor microenvironment biomarkers and clinical activity of ipilimumab in advanced melanoma. J Transl Med, 9:204, Nov 2011.

6. S. Spranger, R. Bao, and T. F. Gajewski. Melanoma-intrinsic β-catenin signalling prevents anti-tumour immunity. Nature, 523(7559):231–235, Jul 2015.

7. M. Luo, H. Wang, Z. Wang, H. Cai, Z. Lu, Y. Li, M. Du, G. Huang, C. Wang, X. Chen, M. R. Porembka, J. Lea, A. E. Frankel, Y. X. Fu, Z. J. Chen, and J. Gao. A STING-activating nanovaccine for cancer immunotherapy. Nat Nanotechnol, 12(7):648–654, July 2017.

8. T. Zhan, N. Rindtorff, and M. Boutros. Wnt signaling in cancer. Oncogene, 36(11):1461– 1473, Mar 2017.

9. R. L. Siegel, K. D. Miller, and A. Jemal. Cancer statistics, 2020. CA Cancer J Clin, 70(1): 7–30, Jan 2020.

10. X. Ye and R. A. Weinberg. Epithelial-Mesenchymal Plasticity: A Central Regulator of Cancer Progression. Trends Cell Biol., 25(11):675–686, Nov 2015.

11. W. Deng, A. Fernandez, S. L. McLaughlin, and D. J. Klinke. WNT1-inducible signaling pathway protein 1 (WISP1/CCN4) stimulates melanoma invasion and metastasis by promoting the epithelial-mesenchymal transition. J. Biol. Chem., 294(14):5261–5280, 04 2019.

12. W. Deng, A. Fernandez, S. L. McLaughlin, and D. J. Klinke. Cell Communication Network Factor 4 (CCN4/WISP1) Shifts Melanoma Cells from a Fragile Proliferative State to a Resilient Metastatic State. Cell Mol Bioeng, 13(1):45–60, Feb 2020.

13. Y. M. Kulkarni, E. Chambers, A. J. McGray, J. S. Ware, J. L. Bramson, and D. J. Klinke. A quantitative systems approach to identify paracrine mechanisms that locally suppress immune response to Interleukin-12 in the B16 melanoma model. Integr Biol (Camb), 4(8): 925–936, Aug 2012.

14. A. Leask. Conjunction junction, what’s the function? CCN proteins as targets in fibrosis and cancers. Am. J. Physiol., Cell Physiol., 318(6):C1046–C1054, 06 2020.

15. K. Meeth, J. X. Wang, G. Micevic, W. Damsky, and M. W. Bosenberg. The YUMM lines: a series of congenic mouse melanoma cell lines with defined genetic alterations. Pigment Cell Melanoma Res, 29(5):590–597, Sept 2016.

16. A. M. Newman, C. B. Steen, C. L. Liu, A. J. Gentles, A. A. Chaudhuri, F. Scherer, M. S. Khodadoust, M. S. Esfahani, B. A. Luca, D. Steiner, M. Diehn, and A. A. Alizadeh. Determining cell type abundance and expression from bulk tissues with digital cytometry. Nat. Biotechnol., 37(7):773–782, July 2019.

17. H. Jeong, I. Hwang, S. H. Kang, H. C. Shin, and S. Y. Kwon. Tumor-Associated Macrophages as Potential Prognostic Biomarkers of Invasive Breast Cancer. J Breast Cancer, 22(1):38–51, Mar 2019.

18. K. R. Jordan, P. Kapoor, E. Spongberg, R. P. Tobin, D. Gao, V. F. Borges, and M. D. McCarter. Immunosuppressive myeloid-derived suppressor cells are increased in splenocytes from cancer patients. Cancer Immunol. Immunother., 66(4):503–513, Apr 2017.

19. S. Ugel, E. Peranzoni, G. Desantis, M. Chioda, S. Walter, T. Weinschenk, J. C. Ochando, A. Cabrelle, S. Mandruzzato, and V. Bronte. Immune tolerance to tumor antigens occurs in a specialized environment of the spleen. Cell Rep, 2(3):628–639, Sep 2012.

20. E. Chun, S. Lavoie, M. Michaud, C. A. Gallini, J. Kim, G. Soucy, R. Odze, J. N. Glickman, and W. S. Garrett. CCL2 Promotes Colorectal Carcinogenesis by Enhancing Polymorphonuclear Myeloid-Derived Suppressor Cell Population and Function. Cell Rep, 12(2):244–257, Jul 2015.

21. C. Wu, H. Ning, M. Liu, J. Lin, S. Luo, W. Zhu, J. Xu, W. C. Wu, J. Liang, C. K. Shao, J. Ren, B. Wei, J. Cui, M. S. Chen, and L. Zheng. Spleen mediates a distinct hematopoietic progenitor response supporting tumor-promoting myelopoiesis. J. Clin. Invest., 128(8):3425–3438, Aug 2018.

22. M. Taki, K. Abiko, T. Baba, J. Hamanishi, K. Yamaguchi, R. Murakami, K. Yamanoi, N. Horikawa, Y. Hosoe, E. Nakamura, A. Sugiyama, M. Mandai, I. Konishi, and N. Matsumura. Snail promotes ovarian cancer progression by recruiting myeloid-derived suppressor cells via CXCR2 ligand upregulation. Nat Commun, 9(1):1685, 04 2018.

23. N. Ferrand, V. B/’ereziat, M. Moldes, M. Zaoui, A. K. Larsen, and M. Sabbah. WISP1/CCN4 inhibits adipocyte differentiation through repression of PPARγ activity. Sci Rep, 7(1):1749, 05 2017.

24. L. Wang, J. Sun, P. Gao, K. Su, H. Wu, J. Li, and W. Lou. Wnt1-inducible signaling protein 1 regulates laryngeal squamous cell carcinoma glycolysis and chemoresistance via the YAP1/TEAD1/GLUT1 pathway. J. Cell. Physiol., Feb 2019.

25. R. H. Muzumdar, X. Ma, S. Fishman, X. Yang, G. Atzmon, P. Vuguin, F. H. Einstein, D. Hwang, P. Cohen, and N. Barzilai. Central and opposing effects of IGF-I and IGF-binding protein-3 on systemic insulin action. Diabetes, 55(10):2788–2796, Oct 2006.

26. M. Mireuta, M. A. Hancock, and M. Pollak. Binding between insulin-like growth factor 1 and insulin-like growth factor-binding protein 3 is not influenced by glucose or 2-deoxy-D-glucose. J. Biol. Chem., 286(19):16567–16573, May 2011.

27. V. Huber, C. Camisaschi, A. Berzi, S. Ferro, L. Lugini, T. Triulzi, A. Tuccitto, E. Tagliabue, C. Castelli, and L. Rivoltini. Cancer acidity: An ultimate frontier of tumor immune escape and a novel target of immunomodulation. Semin. Cancer Biol., 43:74–89, 04 2017.

28. A. Calcinotto, P. Filipazzi, M. Grioni, M. Iero, A. De Milito, A. Ricupito, A. Cova, R. Canese, E. Jachetti, M. Rossetti, V. Huber, G. Parmiani, L. Generoso, M. Santinami, M. Borghi, S. Fais, M. Bellone, and L. Rivoltini. Modulation of microenvironment acidity reverses anergy in human and murine tumor-infiltrating T lymphocytes. Cancer Res., 72(11):2746–2756, Jun 2012.

29. Z. Husain, Y. Huang, P. Seth, and V. P. Sukhatme. Tumor-derived lactate modifies antitumor immune response: effect on myeloid-derived suppressor cells and NK cells. J. Immunol., 191(3):1486–1495, Aug 2013.

30. A. Dongre, M. Rashidian, F. Reinhardt, A. Bagnato, Z. Keckesova, H. L. Ploegh, and R. A. Weinberg. Epithelial-to-Mesenchymal Transition Contributes to Immunosuppression in Breast Carcinomas. Cancer Res., 77(15):3982–3989, 08 2017.

31. S. Terry, P. Savagner, S. Ortiz-Cuaran, L. Mahjoubi, P. Saintigny, J. P. Thiery, and S. Chouaib. New insights into the role of EMT in tumor immune escape. Mol Oncol, 11(7):824–846, 07 2017.

32. E. Alspach, D. M. Lussier, and R. D. Schreiber. Interferon γ and its important roles in promoting and inhibiting spontaneous and therapeutic cancer immunity. Cold Spring Harb Perspect Biol, 11(3):a028480, Mar 2019.

33. D. I. Gabrilovich, S. Ostrand-Rosenberg, and V. Bronte. Coordinated regulation of myeloid cells by tumours. Nat. Rev. Immunol., 12(4):253–268, Mar 2012.

34. T. Condamine and D. I. Gabrilovich. Molecular mechanisms regulating myeloid-derived suppressor cell differentiation and function. Trends Immunol., 32(1):19–25, Jan 2011.

35. W. Li, T. Tanikawa, I. Kryczek, H. Xia, G. Li, K. Wu, S. Wei, L. Zhao, L. Vatan, B. Wen, P. Shu, D. Sun, C. Kleer, M. Wicha, M. Sabel, K. Tao, G. Wang, and W. Zou. Aerobic Glycolysis Controls Myeloid-Derived Suppressor Cells and Tumor Immunity via a Specific CEBPB Isoform in Triple-Negative Breast Cancer. Cell Metab., 28(1):87–103, Jul 2018.

36. B. Guo, X. Huang, M. R. Lee, S. A. Lee, and H. E. Broxmeyer. Antagonism of PPAR-γ signaling expands human hematopoietic stem and progenitor cells by enhancing glycolysis. Nat. Med., 24(3):360–367, 03 2018.

37. L. Lugini, P. Matarrese, A. Tinari, F. Lozupone, C. Federici, E. Iessi, M. Gentile, F. Luciani, G. Parmiani, L. Rivoltini, W. Malorni, and S. Fais. Cannibalism of live lymphocytes by human metastatic but not primary melanoma cells. Cancer Res., 66(7):3629–3638, Apr 2006.

38. A. L. Chang, J. Miska, D. A. Wainwright, M. Dey, C. V. Rivetta, D. Yu, D. Kanojia, K. C. Pituch, J. Qiao, P. Pytel, Y. Han, M. Wu, L. Zhang, C. M. Horbinski, A. U. Ahmed, and M. S. Lesniak. CCL2 Produced by the Glioma Microenvironment Is Essential for the Recruitment of Regulatory T Cells and Myeloid-Derived Suppressor Cells. Cancer Res., 76(19):5671– 5682, 10 2016.

39. M. Hale, F. Itani, C. M. Buchta, G. Wald, M. Bing, and L. A. Norian. Obesity triggers enhanced MDSC accumulation in murine renal tumors via elevated local production of CCL2. PLoS ONE, 10(3):e0118784, 2015.

40. H. Liang, L. Deng, Y. Hou, X. Meng, X. Huang, E. Rao, W. Zheng, H. Mauceri, M. Mack, M. Xu, Y. X. Fu, and R. R. Weichselbaum. Host STING-dependent MDSC mobilization drives extrinsic radiation resistance. Nat Commun, 8(1):1736, 11 2017.

41. T. Hartwig, A. Montinaro, S. von Karstedt, A. Sevko, S. Surinova, A. Chakravarthy, L. Taraborrelli, P. Draber, E. Lafont, F. Arce Vargas, M. A. El-Bahrawy, S. A. Quezada, and H. Walczak. The TRAIL-Induced Cancer Secretome Promotes a Tumor-Supportive Immune Microenvironment via CCR2. Mol. Cell, 65(4):730–742, Feb 2017.

42. B. Huang, Z. Lei, J. Zhao, W. Gong, J. Liu, Z. Chen, Y. Liu, D. Li, Y. Yuan, G. M. Zhang, and Z. H. Feng. CCL2/CCR2 pathway mediates recruitment of myeloid suppressor cells to cancers. Cancer Lett., 252(1):86–92, Jul 2007.

43. D. A. Arenberg, M. P. Keane, B. DiGiovine, S. L. Kunkel, S. R. Strom, M. D. Burdick, M. D. Iannettoni, and R. M. Strieter. Macrophage infiltration in human non-small-cell lung cancer: the role of CC chemokines. Cancer Immunol. Immunother., 49(2):63–70, May 2000.

44. D. S. Hsu, H. J. Wang, S. K. Tai, C. H. Chou, C. H. Hsieh, P. H. Chiu, N. J. Chen, and M. H. Yang. Acetylation of snail modulates the cytokinome of cancer cells to enhance the recruitment of macrophages. Cancer Cell, 26(4):534–548, Oct 2014.

45. R. W. Jenkins, D. A. Barbie, and K. T. Flaherty. Mechanisms of resistance to immune checkpoint inhibitors. Br. J. Cancer, 118(1):9–16, 01 2018.

46. D. Schadendorf, F. S. Hodi, C. Robert, J. S. Weber, K. Margolin, O. Hamid, D. Patt, T. T. Chen, D. M. Berman, and J. D. Wolchok. Pooled Analysis of Long-Term Survival Data From Phase II and Phase III Trials of Ipilimumab in Unresectable or Metastatic Melanoma. J. Clin. Oncol., 33(17):1889–1894, Jun 2015.

47. A. Ribas, O. Hamid, A. Daud, F. S. Hodi, J. D. Wolchok, R. Kefford, A. M. Joshua, A. Patnaik, W. J. Hwu, J. S. Weber, T. C. Gangadhar, P. Hersey, R. Dronca, R. W. Joseph, H. Zarour, B. Chmielowski, D. P. Lawrence, A. Algazi, N. A. Rizvi, B. Hoffner, C. Mateus, K. Gergich, J. A. Lindia, M. Giannotti, X. N. Li, S. Ebbinghaus, S. P. Kang, and C. Robert. Association of Pembrolizumab With Tumor Response and Survival Among Patients With Advanced Melanoma. JAMA, 315(15):1600–1609, Apr 2016.

48. J. A. Flores-Toro, D. Luo, A. Gopinath, M. R. Sarkisian, J. J. Campbell, I. F. Charo, R. Singh, T. J. Schall, M. Datta, R. K. Jain, D. A. Mitchell, and J. K. Harrison. CCR2 inhibition reduces tumor myeloid cells and unmasks a checkpoint inhibitor effect to slow progression of resistant murine gliomas. Proc. Natl. Acad. Sci. U.S.A., 117(2):1129–1138, Jan 2020.

