## Supplementary material for "Cell Communication Network factor 4 promotes tumor-induced immunosuppression in melanoma": PeerReview_CancerResearch

The following Supplement contains the peer-review documents associated with the submission of the pre-print:

To Cancer Research. Items include:

- Decision letter 1.
- Rebuttal letter 1.
- Decision letter 2 and associated peer-review (major revision)
- Rebuttal letter 2.
- Rebuttal letter 3.
- Decision letter 3 and associated peer-review (reject)

**Subject:** RE: CANCER RESEARCH CAN-20-3478 -- Decision Rendered
**Date:** Friday, October 30, 2020 at 2:26:50 PM Eastern Daylight Time
**From:** Echikson, Eileen
**To:** David Klinke
**CC:** journalemails
**Attachments:** image925000.jpg, image833001.png, image955002.png, image872003.png, image729004.png, image576005.png, image892006.png

Hi Dr. Klinke,

It sounds like Dr. Dang is entertaining an appeal if you can provide the information he has requested in the decision letter.

I will forward your appeal letter to him, and let you know what he decides.

Thank you,
Eileen

**Eileen Echikson**
EDITORIAL ASSOCIATE
Cancer Research

American Association for Cancer Research
FINDING CURES TOGETHER

American Association for Cancer Research
615 Chestnut Street, 17th Floor | Philadelphia, PA 19106-4404
215-446-7233 Direct
 | www.AACR.org

DONATE

Please note that this e-mail and any files transmitted with it may be privileged, confidential, and protected from disclosure under applicable law. This information is intended only for the person or entity to which it is addressed and may contain confidential or privileged material. Any review, retransmission, dissemination, or other use of, or taking of any action in reliance upon, this information by persons or entities other than the intended recipient is prohibited. If you received this in error, please contact the sender and delete the material from any computer.

**From:** David Klinke <>
**Sent:** Friday, October 30, 2020 12:38 PM
**To:** Echikson, Eileen <>
**Subject:** Re: CANCER RESEARCH CAN-20-3478 -- Decision Rendered

Hi
Can you clarify? Are they rejecting this – because there is no link to resubmit - or would they consider a revision that addresses the reproducibility of our animal studies?

I'm attaching a PDF of our rebuttal.

Sincerely,
David Klinke
—

David J. Klinke II, Ph.D.
Professor - Department of Chemical and Biomedical Engineering
Adjunct Associate Professor - Department of Microbiology, Immunology & Cell Biology
West Virginia University, Morgantown, WV 26506-6102
(304) 293-9346 | ESB 517 | BMRF 219 | Website: http://klinke.openwetware.org

---

**From:** Cancer Research <>
**Reply-To:** "" <>
**Date:** Monday, October 19, 2020 at 9:15 AM
**To:** David Klinke <>
**Subject:** CANCER RESEARCH CAN-20-3478 -- Decision Rendered

Re: CAN-20-3478
Cell Communication Network factor 4 promotes tumor-induced immunosuppression in melanoma.

Dear Dr. Klinke:

Thank you for submitting your manuscript to Cancer Research.

Our Editors have now completed pre-review of your study and I am sorry to say that the manuscript was not judged to reach the overall priority that would be needed for further consideration. **Therefore, I am returning it to you with suggestions.**

In principle, we are interested in your work, which potentially provides additional insights into Wisp1/CCN4. As you may know, we are quite concerned about the reproducibility of tumor animal studies and this was reflected in my recent CR editorial (PMID: 32878863) aiming to improve the reporting of animal studies. In this regard, the animal studies reported in Figure 2 appear to have inadequately described cohort sizes. For example, the study of B16F0 control vs CRISPR-mediated Wisp1 KO tumors indicates that clone KO1 had "significantly decreased" tumor weights in C57BL/6 mice with animal numbers greater or equal to 5. Please describe precisely the numbers of animals in each group and address why the results in Figure S2b for clone B16F0-KO2 (n=6 each group control vs. KO) in your BioRxiv manuscript (https://www.biorxiv.org/content/10.1101/427088v1.full.pdf) did not show a difference in tumor growth in C57BL/6. This seems to counter the claim in your current manuscript. Further the study reported in Figure 2B seems to have imbalanced cohort groups (n=3 versus n=8) construed to control for puromycin resistance. The small number n=3 seems inadequate to conclude that puromycin resistance did not affect survival. **Overall, please indicate the reproducibility of your animal studies in an adequately revised manuscript.**

We regret we could not render a more favorable decision on your manuscript. However, we appreciate your consideration of the concerns raised regarding your current submission. Thank you very much for your interest in Cancer Research.

Sincerely,
Chi Van Dang, MD, PhD
Editor-in-Chief
(On behalf of the Senior Editorial Board)

**AACR Manuscript Transfer Service**

The AACR offers authors the option to easily transfer a declined Research Article to a second AACR journal of your choice for editorial consideration. At your request, all files, manuscript information, and reviews (if applicable) will be transferred immediately. For details of this service, please visit the AACR Manuscript Transfer FAQ page.

Click the link below to view your options and (if desired) begin the transfer process.

https://can.msubmit.net/cgi-bin/main.plex?el=DELETED

**External email. Please do not click links or open attachments unless you recognize the source.**

**Response to Reviewer's Comments**

**Cell Communication Network factor 4 promotes tumor-induced immunosuppression in melanoma.**

Audry Fernandez, Wentao Deng, Stephanie Rellick, Sarah McLaughlin, Anika C. Pirkey, and David J. Klinke II

Submitted for review in Cancer Research (Manuscript ID: CAN-20-3478)

Our responses to the specific comments are listed below in blue font.

Comments from Editor-in-Chief:

1) In principle, we are interested in your work, which potentially provides additional insights into Wisp1/CCN4. As you may know, we are quite concerned about the reproducibility of tumor animal studies and this was reflected in my recent CR editorial (PMID: 32878863) aiming to improve the reporting of animal studies. In this regard, the animal studies reported in Figure 2 appear to have inadequately described cohort sizes. For example, the study of B16F0 control vs CRISPR-mediated Wisp1 KO tumors indicates that clone KO1 had "significantly decreased" tumor weights in C57BL/6 mice with animal numbers greater or equal to 5. Please describe precisely the numbers of animals in each group.

   Author Response (AR): We agree that reproducibility is important. To clarify our results, we have now included a revised Figure 2 that shows the overall numbers included in each cohort and the full tumor growth curves. Following from our recent publication (Deng et al. Cell Mol Bioeng 2020 (bioRxiv 487751; doi: https://doi.org/10.1101/487751) which we published after the BioRxiv manuscript mentioned below), we have also included a quantitative analysis of these data. As is clear in these data figures, tumor growth of these models follow a log-linear growth profile. Among replicates, the initial bolus of tumor initiating cells varies despite injecting the same number of cells but the tumor growth rate is fairly consistent.

2) Please address why the results in Figure S2b for clone B16F0-KO2 (n=6 each group control vs. KO) in your BioRxiv manuscript (https://www.biorxiv.org/content/10.1101/427088v1.full.pdf) did not show a difference in tumor growth in C57BL/6. This seems to counter the claim in your current manuscript.

   AR: Couple of comments here. **First,** Figure S2 (panel A and B) shows the results generated using B16F10 cells – these were selected originally for metastatic potential – and not B16F0 cells using in this manuscript. **Second,** while there were n = 6 in the cohort, the average tumor size for each time point is not comprised of six measurements. For instance, the Day 7 and Day 15 time points have less than six as some of the tumors appeared earlier in some of the mice and thus needed to be euthanized earlier than mice that has tumors appear later. The variability among mice of a cohort can be seen in the new Figure 2.

**Third,** the figures are annotated with the slope of the linear regression line that is fit to the averages calculated at each time point. Performing linear regression to averages is a bit unrefined as it lumps the number of tumor-initiating cells and the growth rates together and thus can flatten the curve giving an inaccurate estimate of the tumor growth rate. Moreover, the slopes are heavily influenced by extrema. Those limitations aside, the slope of the KO in Figure S2B appears lower than the WT slope (0.139 versus 0.161). In addition, the slope of the growth curve of KO tumors implanted in NSG mice appears greater than WT tumors (0.248 versus 0.243). This implies that CCN4 KO increases the intrinsic growth rate of the tumors, which is quantified by the NSG experiments, but also increases immune-mediated control of tumor growth, which is quantified by the difference in the rate of tumor growth between C57BL/6 and NSG experiments (for comparison with new Figure 2H: log10 [ kG|NSG / (kG- kD)|C57BL/6] WT: log10(0.243/0.161) = 0.179; CCN4 KO: log10(0.248/0.139) = 0.251). The observations shown in Fig S2b are consistent with what we report with the YUMM1.7 and B16F0 lines.

**Fourth,** a follow-on paper developed a more refined approach to analyze these data (Deng et al. Cell Mol Bioeng 2020 (bioRxiv 487751; doi: https://doi.org/10.1101/487751)) that we employ in this new Figure 2. We note that all of the CCN4 KO variants of the YUMM1.7 and B16F0/B16F10 generated using either a double nickase or Homology Directed Repair (HDR) CRISPR/Cas9 strategy exhibited a consistent response. One experiment using a pool of CCN4 KO variants of B16F10 cells generated using a HDR strategy was different, however (shown for transparency in Supplemental Figure S9B). It turned out that the cells that we injected had lost the CCN4 KO as they were cultured for a period of time without puromycin selection prior to injection. That is why we didn't proceed to test these cells in NSG mice.

3) Further the study reported in Figure 2B seems to have imbalanced cohort groups (n=3 versus n=8) construed to control for puromycin resistance. The small number n=3 seems inadequate to conclude that puromycin resistance did not affect survival. Overall, please indicate the reproducibility of your animal studies in an adequately revised manuscript.

AR: In the interest of brevity, we have removed that panel as the control included in Figure S2 – DNMT3A KO – serves as a more comprehensive control for CRISPR/Cas9 gene editing instead of just puromycin selection.

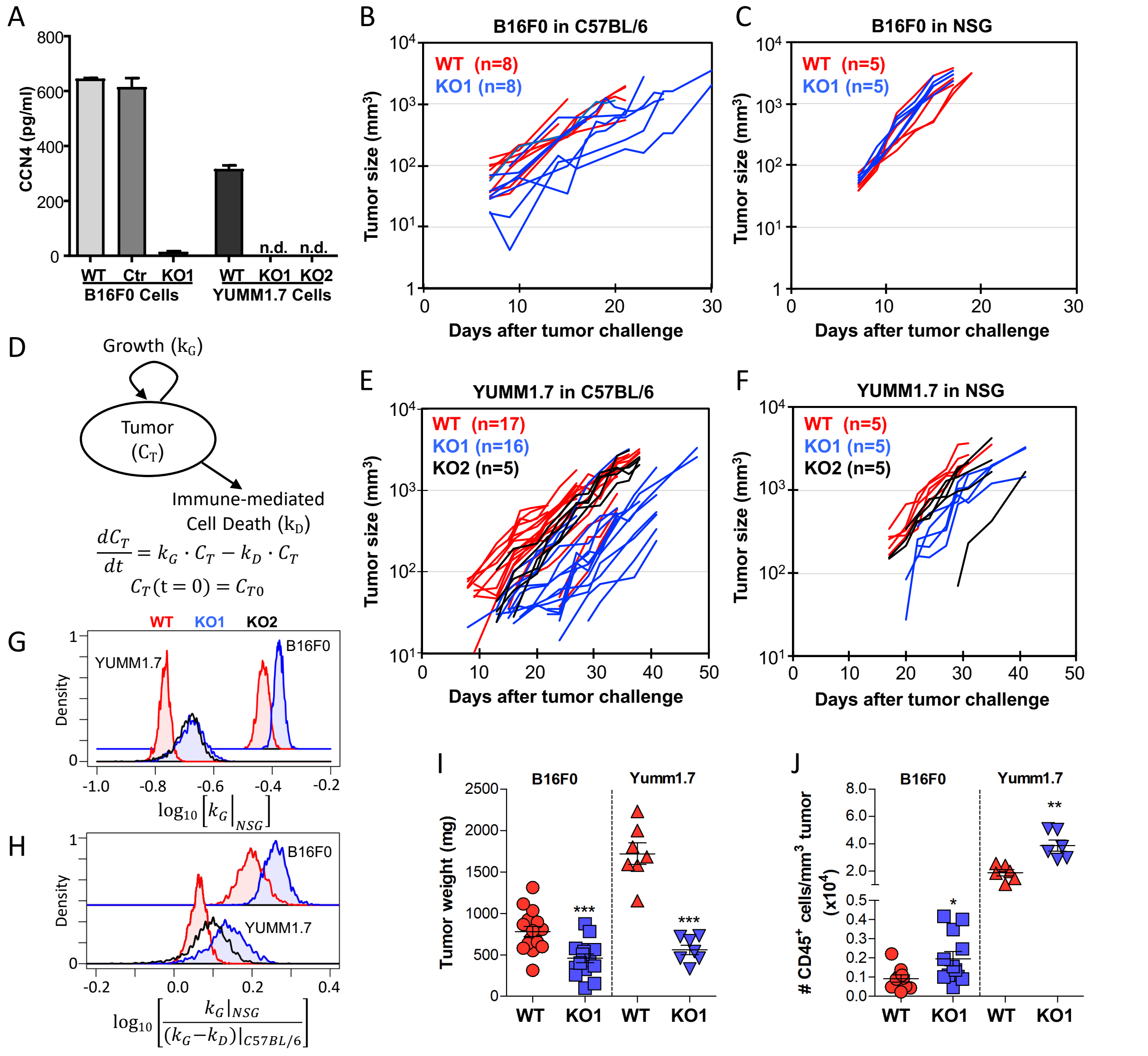


$$\frac{dC_T}{dt} = k_G \cdot C_T - k_D \cdot C_T$$
$$C_T(t=0) = C_{T0}$$

$$\log_{10}\left[k_G|_{NSG}\right]$$

$$\log_{10}\left[\frac{k_G|_{NSG}}{(k_G - k_D)|_{C57BL/6}}\right]$$

Figure 2 - CCN4 knockout suppressed melanoma tumor growth in immunocompetent mice} (A) CCN4 secretion after CRISPR/Cas9 knockout in mouse spontaneous B16F0 and YUMM1.7 $Braf^{V600E/WT}$ $Pten^{-/}$ $Cdkn2^{-/-}$ cell lines. Cell culture media conditioned for 48 hours by the indicated cell line were tested by ELISA (n.d. = not detected). All knockouts were generated through CRISPR/Cas9 methodology. B16F0 knockout created with Homology-Directed Repair approach with controls for puromycin selection (B16F0-Ctr) created with pBabe-puro retrovirus. YUMM1.7 knockout created with double nickase approach with KO1 and KO2 indicating two different clones. Wildtype (WT: red) B16F0 (B,C) and YUMM1.7 (E,F) cells and CCN4 knockout variants (KO1: blue, KO2: black) were subcutaneously injected into (B,E) C57BL/6 immunocompetent and (C,F) NSG immunocompromised mice. Average tumor volume measured by caliper as a function of time after tumor challenge. (D) Schematic of the tumor growth model used to interpret the log-exponential tumor growth curves. Posterior distributions in (G) the proliferation rate of tumors in NSG mice and in (H) the log-ratio of the proliferation rate in NSG relative to the net growth rate of tumors in C57BL/6 mice. (I) In a time-matched experiment, average tumor weights of both B16F0 and YUMM1.7 were significantly decreased following CCN4 knockout (KO1). (J) Significant increases in the number of CD45+ cells were observed in both B16F0 and YUMM1.7 cell lines following CCN4 knockout (KO1). * $0.01<p<0.05$, ** $0.001 <p<0.01$, *** $p<0.001$.

**Subject:** CANCER RESEARCH CAN-20-3478R-A -- Decision Rendered
**Date:** Tuesday, December 29, 2020 at 11:13:33 AM Eastern Standard Time
**From:** Cancer Research
**To:** David Klinke

Re: CAN-20-3478R-A
Cell Communication Network factor 4 promotes tumor-induced immunosuppression in melanoma.

Dear Dr. Klinke:

Thank you for submitting your manuscript to Cancer Research. Please see the reviewer comments appended below. Based on these comments as well as the confidential recommendations and priority scores, we would potentially be willing to evaluate a revised revision. Any such revision would likely need to be returned to the reviewers for additional advice.

Please carefully review the Instructions for Resubmission appended below. As you will see, the reviewers feel that significant additional work is necessary before it can be further considered for publication.

Please note that the processing of manuscripts not prepared in accordance with the Journal's criteria may be delayed and/or returned to you for correction prior to receiving additional consideration.

We look forward to receiving your properly revised manuscript and ask that you return it within three months. If you require additional time, please contact the Cancer Research Editorial Office via email at.

We appreciate the opportunity to review your work.

You may resubmit your manuscript at the following link:
https://can.msubmit.net/cgi-bin/main.plex?el=DELETED

Sincerely,
Chi Van Dang, MD, PhD
Editor-in-Chief

**CONFIDENTIAL TO AUTHORS**

Reviewer #1 (Reviewer Comments to the Author):

In this manuscript presented by Fernandez et al entitled "Cell Communication Network factor 4 promotes tumor-induced immunosuppression in melanoma", the authors demonstrate the immunomodulatory properties of the matricellular protein CCN4. Using CRISPR to knockout CCN4 in two melanoma cell lines B16F0 and YUMM1.7. The authors show that CCN4 expression is correlated with survival in human melanoma using TCGA data, although the sample size is low for this study and the correlation should be interpreted with caution. Further, the authors provide evidence in murine models that knockout of CCN4 results in reduced tumor growth, increased T and NK cell infiltration and reduced MDSC infiltration in tumors. Chemokines CCL2 and CXLC1 are regulated by CCN4 which recruit said MDSC which subsequently reduce T cell proliferation. Lastly, the authors demonstrate that PD-1 targeted checkpoint blockade is more efficacious in the CCN4 knockout tumor models than the wildtype tumors.
1. Throughout the manuscript, the authors make several sweeping statements while lacking evidence to support

their claims as follows:
a) Implanting tumors into NSG mice, the authors state that (P6 L228) "In immunocompetent mice, we assumed that the net growth rate reflects both the intrinsic growth rate and a loss in tumor size due to immune-mediated cell death (kD), which lumps all mechanisms for anti-tumor immunity into a single constant" (Fig 1). The authors derived an equation assuming the loss of tumor is due to the immune system and attribute all possible mechanisms of antitumor immunity into a single constant. While NSG mice lack lymphocytes, they still possess myeloid cells and alternative immune-related tumor surveillance mechanisms, albeit impaired. The authors therefore cannot ascribe all potential mechanisms of antitumor immunity to one single factor/mechanism.
b) When describing data in Figure 3, the authors state (P6 L272) "Collectively, these data suggest that CCN4 skews the immune contexture within the melanoma microenvironment by reducing the infiltration of cells with potent anti-tumor cytolytic activity, namely CD8+ T cells and NK cells, and by enhancing the prevalence of cells with immunosuppressive capacity, namely MDSC." The authors, however, have not demonstrated the cytolytic antitumor capacity of infiltrating CD8 T cells or NK cells. Indeed, these cell type are generally thought to be cytotoxic, yet no direct tumor-killing ability nor by what potential lytic mechanism, has been demonstrated. While they do later in the manuscript perform experiments demonstrating the ability of MDSC to suppressive T cell proliferation, this is not shown within the context of Figure 3.
c) In discussing Figure 7 and the potential synergy between CCN4 knockout and checkpoint blockade therapy, the authors state (P8 L379) "It follows then that, by knocking out CCN4, the PD1-PDL1 axis plays a more dominant role in the YUMM1.7 model, likely through IFNγ cross-talk, in restraining the T cell response and is more sensitive to therapeutic intervention. We also noted that the infiltration of T cells and NK cells were almost 10-times lower per gram tumor in the WT B16F0 model compared with the WT YUMM1.7 model. Those differences coupled with B16F0 sensitivity to αCTLA4 suggests that pathways associated with initiating an anti-tumor response, such as the extent of clonal T cell expansion within secondary lymphoid organs, are key limiters in the B16F0 model. Overall, these data indicate that absence of tumor-derived CCN4 potentiates the anti-tumor effect of immune checkpoint blockade therapy." Particularly, the authors do not demonstrate a role for IFNγ in mediating tumor clearance during PD-1 inhibition (e.g. Does IFNγ increase after PD-1 blockade in your model? Further, while the literature suggests that checkpoint blockade may play a role in clonal expansion, no such data was demonstrated within either of the models presented in this manuscript.

2. The gating hierarchy of immune cell populations Figure 3 gating hierarchy does not reflect markers used in text. For instance, in the text, NK cell gating is described as "Live CD45+ CD3e− NK1.1+ B220− events" (P6 L250), yet the biaxial gating demonstrates that NK1.1+B220- and NK1.1+B220+ cells were gated. Similarly, the gate for CD11blowGr-1+ TANs and CD11bhighGr-1+ MDSC overlaps.

3. In Figure 3N, the authors correlate the tumor size with infiltration of CD11b+Gr1+ events in various tumor models. However, the authors only use 2 ID mCCN4 + Dox tumors and the large YM1.7 KO tumors overlap with the small WT Yum1.7 tumors, making it difficult to interpret how tumor size influences MDSC recruitment.
4. In Figure 5, the authors chose to pursue expression of CXCL1 and CCL2 out of several of the differentially expressed cytokines in the Cytokine Array. Can they provide further rationale for choosing those specific two chemokines out of the entire chemokine set in the array? Further, it would be helpful to provide the ECAR and OCAR stress test profile provided by the Seahorse Assay to fully interpret the glycolytic flux.

5. In Figure 7, the authors seek to combine checkpoint blockade therapy with CCN4 knockout. They start treatment when tumors reach 100mm3 which happens to be later in the knockout model due to slowed growth kinetics. Given that PD-1 targeted monotherapy does not appear to be curative, if treatment was initiated earlier, might monotherapy be curative? Is CTLA-4/PD-1 combination synergistic?

6. Can the authors comment on effects of PD1 therapy in the knockout model? In the wild type tumor model, a clear difference in tumor size is seen between untreated and PD-1 treated mice. In the knockout model, the differences in tumor size are much smaller between the untreated and treated groups. In this case a significant difference is demonstrated 2 days after treatment, and while by day 3 the difference in tumor volume is still significant, the magnitude of the differences appears smaller than in the wild type setting.

Reviewer #2 (Reviewer Comments to the Author):

The current manuscript by Fernandez and colleagues describes a role for Cell Communication Network Factor 4 (CCN4) in promoting tumor-induced immunosuppression. This follows on from previous data from this group identifying this factor as a promotor EMT transition (Refs 11 and 12 from current manuscript). EMT transition has previously been shown to promote immunosuppression through a number of mechanisms and thus in the current study the authors explore the hypothesis that CCN4 directly regulates the immune response. The study is novel and of interest but there are technical limitations to the study that should be addressed to strengthen the conclusions.

1) Throughout the paper, the authors utilise a DNMT3a KO tumor as a control. What is the justification for this over a non-targeting control/ vector only control? Presuming the authors are referring to DNA (cytosine-5)-methyltransferase 3A, knockout of this gene would likely lead to significant transcriptional and epigenetic changes, thus making comparisons with this control irrelevant. This should be clearly explained and justified and experiments performed to ensure this "control" has no phenotype.

2) In Figure 2E none of the mice challenged with YUMM1.7 tumors achieve long-term survival but in Figure 4 this appears to be 20-40% (without or with Dox respectively). How do the authors explain this? One explanation is that the rtta element used for the inducible expression of CCN4 is immunogenic in wild-type mice (refer to PMID 20155386). If this is the case it would be a significant caveat for these experiments.

3) Several of the data are interesting but quite descriptive e.g. reduced expression of chemokines associated with immunosuppression, reduced MDSCs, reduced T cell activation in vitro. However, the mechanism by which CCN4 leads to enhanced tumor growth is not defined. As a minimum the author should present data with T cell depletion/ targeting one of these factors to identify the key driver of the in vivo phenotype.

4) In Figure 1 the authors present data suggesting that Knockout of CCN4 leads to greater tumor control. Whilst it is useful to have the individual growth kinetics from each mouse, this should be supported by data indicating the mean +/- SEM for these experiments to understand whether these differences are statistically significant. The presented formula attempts to do this, but is not clear why such an equation is needed to confirm this important point.

5) The authors suggest that targeting CCN4 enhances the effect of immune checkpoint blockade. However this is not borne out by the data where the magnitude of response is similar. I suggest this data is removed or conclusions modified.

Minor Points
1) Figure 1A is reproduced from one of the author's previous publications. Whilst this is not a major issue given it is a bioinformatics analysis, this should be acknowledged in the figure legend.

2) Several of the figure panels are called out non sequentially.

Reviewer #3 (Reviewer Comments to the Author):

This is an elegant paper on the effects of CCN4 in melanoma. The manuscript is clearly written, and the figures are clearly displayed to illustrate particular points made in the manuscript. The authors show that CCN4 acts as an immune-suppressor in melanoma. The authors propose that the main activity of CCN4 is through the inhibition of MDSC function. They demonstrate both direct effects, such as release by melanoma cells of chemokines to recruit MDSCs, and, inhibition of IFNg release by T cells. Checkpoint inhibitor therapy potentiates the effects of CCN4 knock-out.
Criticisms: the effect of knock-down in B16FO cells seems to be marginal at best, and is therefore not explored further. Does this really make a significant contribution to the manuscript. If not, I suggest to remove these data, as they reduce the impact of all other data shown.
Th ultimate mechanism by which a matricellular protein affects the release of chemokines is not explored herein, and

can safely be left for a future paper. Nevertheless, the authors are encouraged to speculate on this potential mechanism.

**RESUBMISSION INSTRUCTIONS**

Cancer Research holds revised manuscripts to a higher standard than initial submissions. Please review the condensed resubmission instructions below to ensure that your revised paper is prepared and submitted in accordance with journal policies. Failure to do so may result in a delay of consideration, with the paper being returned for correction prior to potential re-review.

For full details on publication policies, format requirements, and more, please consult the Editorial Policies section of the AACR's Information for Authors: https://aacrjournals.org/content/authors/editorial-policies

Please direct any questions to the Cancer Research Editorial Office:

**Cover Letter**

With your revision, you may supply a cover letter to the editor, but please provide your response to the reviewers separately as detailed below.

**Response to Reviewer Comments**

Include a detailed point-by-point response to **all** reviewer comments. Fully explain how and where you addressed each comment in the revised manuscript. You may also include, for review only, a tracked-changes, marked or highlighted manuscript that indicates where changes were made.

**Author List**

Please review the current author list from your submission below and, if applicable, add to the online submission form any new authors who contributed to the study during its revision or any authors who were inadvertently excluded during initial submission. As the corresponding author, you are ultimately responsible for ensuring the accuracy of the list as it appears in SmartSubmit:

Audry Fernandez, Wentao Deng, Sarah L. McLaughlin, Anika C Pirkey, Stephanie L Rellick

If any authorship changes are made during revision, you must submit a signed authorship change form signed by every author, including any author(s) who have been added or removed. Note that authorship changes constitute the addition or removal of contributors; the modification of the author listing order; and the modification of co-authorship.

A blank authorship change form is available here:
https://www.aacrjournals.org/sites/default/files/downloads/authors/Authorship_Change_Form.pdf

Please upload the signed form to SmartSubmit as a "Supplementary Data (for Review only)" file upon resubmission.

**Manuscript Text and Length**

Provide an unmarked version of your manuscript text with all pages numbered and organized into the following sections and order: title page, abstract, materials and methods, results, discussion, acknowledgments, references, tables, and figures with legends.

Please familiarize yourself with journal requirements on manuscript length and figure/table number restrictions. As necessary, we ask that you shorten your presentation wherever achievable and revise the manuscript as succinctly as possible for the field's benefit. Word and display item limits are detailed on the journal information page at:

https://cancerres.aacrjournals.org/site/misc/journal_ifora.xhtml

**Title and Abstract**
Abbreviations, unless commonly used in the literature, must be spelled out in the title. The abstract must be in an unstructured format and may not contain references or exceed 250 words.

**Title Page**
The title page should include the title, author list, running title, key words, financial support, and the corresponding author's full name, mailing address, and email address. A conflict of interest disclosure statement must be included, even if the authors have no potential conflicts of interest to disclose; more information is available at: https://aacrjournals.org/content/authors/editorial-policies#coi

**Figures and Tables**
Figures must be submitted in their final color format: i.e., figures to be printed in black and white must be submitted in black and white, and figures to be printed in color must be submitted in color. We do not offer the option to publish a figure online in color and in print in black and white. Color figures are published at a cost to the corresponding author of US$625 (members) or US$675 (non-members) per figure.

Tables may not exceed one page in length. Tables are not permitted within figures.

Formatting and best practices guidelines for tables and figures are available at: https://aacrjournals.org/content/authors/article-style-and-format#display

**Supplementary Data and Methods**
Supplementary data should not be included in the main manuscript file. Any supplementary data uploaded as "Supplementary Data (for Review only)" will not be published but will be available to the editor and reviewers.

It is unacceptable to move the majority of the Materials and Methods section to the supplement in order to shorten the manuscript. Additional details such as instrumentation settings, recombinant constructions, and oligonucleotide or protein sequences may be included in the supplement, but the main Materials and Methods section alone should provide readers with sufficient information to be able to reproduce the paper's results. This policy is strictly enforced. If necessary, the manuscript length guidelines can be relaxed upon request to allow appropriate reporting of the methods in the main Materials and Methods section.

Supplementary data files must not exceed 8 in number or 50 MB for any individual file. PDF is the preferred format for all supplementary data, except for tables created in Excel.

When uploading supplementary data files, please provide brief captions in the "Description" fields on the submission form. These captions should describe each file's contents without reiterating the title (i.e., it is acceptable to provide "Figure S1," for example, as a title but NOT as a caption; instead, the caption should read, "Figure S1 shows x, y, and z...").

**Studies that describe RNA interference (RNAi)**
**Studies that describe RNA interference (RNAi) must contain one non-specific control and two specific siRNAs for each knockdown target of interest. Additional data is required that demonstrates knockdown achievement at the level of protein expression (not just mRNA).**

**Cell Lines**
**If cell lines were used in the research, the following points must be addressed in the Materials and Methods section of the manuscript:**

**-- From where and when the cells were obtained.**
**-- A description of the cell line authentication and Mycoplasma testing regimen used by the laboratory and, minimally, the latest date the cells were tested. If no testing was performed, then this should be explicitly stated in**

**the text.**
**-- The method(s) by which the cells were tested and authenticated, if applicable.**
**-- The length of time or number of passages between collection or thawing and use in the described experiments.**

**Full details on the AACR's cell line policy can be found here: https://aacrjournals.org/content/authors/editorial-policies#celllines**

**Key Resource Identifiers (RRIDs)**
**All key resources used in the research should be clearly and uniquely identified prior to publication. Please use the information provided in the Key Resources Table contained in the SciScore report you received during the submission process to add RRIDs for all listed reagents and other resources.**

**For more information on adding RRIDs, please consult the RRID section at: https://aacrjournals.org/improv-reprod**

**Data and Materials Availability**
**Data for which community-recognized, structured repositories exist (e.g., natural nucleotide or amino acid sequences, microarray data, small molecule and macromolecular structure data) must be deposited in an appropriate repository no later than submission of a revised manuscript. Failure to do so may result in delayed acceptance and/or publication. For all deposited data, the accession number(s) must be included in the manuscript and the database record must be publicly accessible on or before the first online publication date.**

**Unique materials, such as cells, plasmids, antibodies, animal models, and computer programs that were used in the research reported and that are not available from commercial suppliers, must be made available, either through the authors' laboratories or organizations or through an appropriate collection or repository, with corresponding accession numbers added to the manuscript as necessary.**

**For more details on these policies, please consult: https://aacrjournals.org/content/authors/editorial-policies#availability**

**Patient Studies**
**If the paper describes patient studies, please confirm in the Materials and Methods section of the manuscript that you and your colleagues obtained written informed consent from the patients, that the studies were conducted in accordance with recognized ethical guidelines (e.g., Declaration of Helsinki, CIOMS, Belmont Report, U.S. Common Rule) and that the studies were approved by an institutional review board. It is not acceptable to include this information in the supplementary data.**

**Animal Experiments**
**If your paper describes animal experiments, please confirm in the Materials and Methods section of the manuscript that an IACUC or other review board has approved these studies. It is not acceptable to include this information in the supplementary data.**

**Resubmission Link**
**Please submit your revised manuscript via the link below:**
**https://can.msubmit.net/cgi-bin/main.plex?el=DELETED**

**Response to Reviewer's Comments**

**Cell Communication Network factor 4 promotes tumor-induced immunosuppression in melanoma.**

Audry Fernandez, Wentao Deng, Stephanie Rellick, Sarah McLaughlin, Anika C. Pirkey, and David J. Klinke II

Submitted for review in Cancer Research (Manuscript ID: CAN-20-3478R-A)

Overall, we thank the reviewers for their service and their constructive criticisms on the manuscript. We feel that this revised paper presents, in a more comprehensive way, our findings related to how Cell Communication Network factor 4 (CCN4) creates an immunosuppressive microenvironment in the context of melanoma. Our responses to the specific comments are listed below in blue font. In the tracked changes document, a red font is used to indicate the text that has been changed in this revised manuscript.

**Reviewer 1**
**General Comments:**
In this manuscript presented by Fernandez et al entitled "Cell Communication Network factor 4 promotes tumor-induced immunosuppression in melanoma", the authors demonstrate the immunomodulatory properties of the matricellular protein CCN4. Using CRISPR to knockout CCN4 in two melanoma cell lines B16F0 and YUMM1.7. The authors show that CCN4 expression is correlated with survival in human melanoma using TCGA data, although the sample size is low for this study and the correlation should be interpreted with caution. Further, the authors provide evidence in murine models that knockout of CCN4 results in reduced tumor growth, increased T and NK cell infiltration and reduced MDSC infiltration in tumors. Chemokines CCL2 and CXLC1 are regulated by CCN4 which recruit said MDSC which subsequently reduce T cell proliferation. Lastly, the authors demonstrate that PD-1 targeted checkpoint blockade is more efficacious in the CCN4 knockout tumor models than the wildtype tumors.
Specific Criticisms:
Throughout the manuscript, the authors make several sweeping statements while lacking evidence to support their claims as follows:

1a) Implanting tumors into NSG mice, the authors state that (P6 L228) "In immunocompetent mice, we assumed that the net growth rate reflects both the intrinsic growth rate and a loss in tumor size due to immune-mediated cell death (kD), which lumps all mechanisms for anti-tumor immunity into a single constant" (Fig 1). The authors derived an equation assuming the loss of tumor is due to the immune system and attribute all possible mechanisms of antitumor immunity into a single constant. While NSG mice lack lymphocytes, they still possess myeloid cells and alternative immune-related tumor surveillance mechanisms, albeit impaired. The authors therefore cannot ascribe all potential mechanisms of antitumor immunity to one single factor/mechanism.

Author Response (AR): The NOD.Cg-Prkdc$^{scid}$ Il2rg$^{tim1Wjl}$/SzJ (NSG) mouse is one of the most highly immunodeficient mouse model currently available for research. This has allowed them to become the most widely applicable mouse strain for cancer xenograft modeling as well as a humanized model by engrafting human CD34+ HSCs. While the NSG mice used in this study do still possess myeloid cells, all cell types are considered significantly defective. In addition, the complement system is also absent from these mice (1). These alterations have essentially rendered the immune system in the mouse highly deficient. The single anti-tumor immunity constant then represents the difference in the growth of the same tumor cells in immunocompetent C57BL/6 mice relative to severely immunocompromised NSG mice. Any residual immune function that would reduce the growth of tumors in NSG is implicitly lumped into the intrinsic growth rate. We have edited the text to help clarify this point. We note, though, that this residual immune function is negligible. The ability to engraft non-malignant human HSC into this mouse line speaks to the degree of immunodeficiency in that strain.

(1) The Jackson Laboratory, "NSG Variants Portfolio", 2021. https://www.jax.org/jax-mice-and-services/find-and-order-jax-mice/nsg-portfolio# (accessed 1/12/2021).

1b) When describing data in Figure 3, the authors state (P6 L272) "Collectively, these data suggest that CCN4 skews the immune contexture within the melanoma microenvironment by reducing the infiltration of cells with potent anti-tumor cytolytic activity, namely CD8+ T cells and NK cells, and by enhancing the prevalence of cells with immunosuppressive capacity, namely MDSC." The authors, however, have not demonstrated the cytolytic antitumor capacity of infiltrating CD8 T cells or NK cells. Indeed, these cell type are generally thought to be cytotoxic, yet no direct tumor-killing ability nor by what potential lytic mechanism, has been demonstrated. While they do later in the manuscript perform experiments demonstrating the ability of MDSC to suppressive T cell proliferation, this is not shown within the context of Figure 3.

AR: We have changed the word "potent" to "potential". We do have data showing that pmel1 tgCD8+ T cells can kill YUMM1.7 cells (see Fig C1 below) and that CCN4 KO in YUMM1.7 cells has no impact on that ability. In the interest of brevity (and space limitations of Cancer Research), we decided not to include these data here but will be included in a separate manuscript. These data plus the expression levels of H-2Kb and H-2Db suggest that there isn't a loss of MHC class 1 processing and presentation.

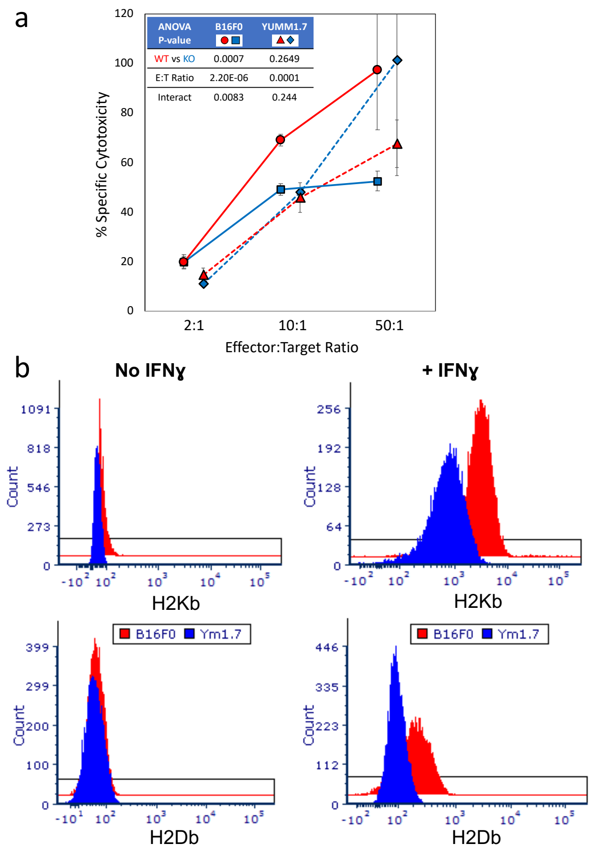


Figure C1. pmel1 tgCD8+ T cells recognize and kill both WT and CCN4 KO B16F0 and YUMM1.7 cells. (a) Splenocytes from pmel1 naïve mice were pre-activated in vitro for two days with 1 µM of the pmel1 CD8+ T cell specific peptide (hgp100$_{25-33}$ (KVPRNQDWL)) in the presence of 100 U/mL of rmIL-2. On days two and four the cells were washed and re-cultured with fresh media containing 100 U/mL of rmIL-2. Pre-activated pmel1 CD8+ T cells were co-cultured for 4 hours with target cells, at the indicated effector to target ratio, to assess cytotoxicity that was quantified using a CyQUANT LDH Cytotoxicity assay (Invitrogen). Wild-type and CCN4 KO B16F0 and YUMM1.7 cell lines were used as targets, after an overnight incubation with 250 U/mL of rmIFNg. Mean +/- sd represent collective results from three biological replicates. Two-factor ANOVA with replication was used to assess statistical differences. (b) Expression of H-2K$^b$ (top panels) and H-2D$^b$ (bottom panels) was assayed by flow cytometry in WT B16F0 (red) and YUMM1.7 (blue) cells without pre-conditioning (left panels) or after an overnight incubation with 250 U/mL of rmIFNg (right panels). In part, differences can be explained by expression levels of H-2Db, as the gp100 peptide is restricted to H-2D$^b$ and YUMM1.7 cells express lower levels of H-2Db after the IFNg treatment compared to B16F10 cells.

1c) In discussing Figure 7 and the potential synergy between CCN4 knockout and checkpoint blockade therapy, the authors state (P8 L379) "It follows then that, by knocking out CCN4, the PD1-PDL1 axis plays a more dominant role in the YUMM1.7 model, likely through IFNγ cross-talk, in restraining the T cell response and is more sensitive to therapeutic intervention. We also noted that the infiltration of T cells and NK cells were almost 10-times lower per gram tumor in the WT B16F0 model compared with the WT YUMM1.7 model. Those differences coupled with B16F0 sensitivity to αCTLA4 suggests that pathways associated with initiating an anti-tumor response, such as the extent of clonal T cell expansion within secondary lymphoid organs, are key limiters in the B16F0 model. Overall, these data indicate that absence of tumor-derived CCN4 potentiates the

anti-tumor effect of immune checkpoint blockade therapy." Particularly, the authors do not demonstrate a role for IFNγ in mediating tumor clearance during PD-1 inhibition (e.g. Does IFNγ increase after PD-1 blockade in your model? Further, while the literature suggests that checkpoint blockade may play a role in clonal expansion, no such data was demonstrated within either of the models presented in this manuscript.
AR: While the experiment using immune checkpoint antibodies was not designed to test for synergy and we didn't explicitly use the term "synergy", we have revised the text to limit that implication. Regarding this paragraph, we are trying to interpret the results that we have obtained and have included adjectives of uncertainty, like "seemed to", "likely", and "suggest", to imply that these statements are interpretations and not clear demonstrations. Yet, the overall statement at the end of this paragraph - that the absence of tumor-derived CCN4 complements the anti-tumor effect of immune checkpoint blockade therapy in two mouse models of melanoma - has been demonstrated. The importance of IFNg in mediating tumor clearance is well-established following from the work of Robert Schreiber. We do show that 1) CCN4 inhibits antigen-induced IFNg production by CD8 T cells (Fig 6), 2) IFNg upregulates PD-L1 and H-2Kb expression by tumor cells (Fig 7 B), 3) CCN4 KO is associated with no difference in the PD-1+ fraction of CD8+ CD3+ TILs (Fig 7D-F), 4) CCN4 KO increases the prevalence of PD-L1+ and H-2Kb+ CD45- cells (predominantly tumor cells) within the tumor, and that 5) prevalence of Treg TILs were not changed upon CCN4 KO (Fig. S7). These data provide some evidence of how aPD1 is working in the YUMM1.7 model.

2) The gating hierarchy of immune cell populations Figure 3 gating hierarchy does not reflect markers used in text. For instance, in the text, NK cell gating is described as "Live CD45+ CD3e− NK1.1+ B220− events" (P6 L250), yet the biaxial gating demonstrates that NK1.1+B220- and NK1.1+B220+ cells were gated. Similarly, the gate for CD11blowGr-1+ TANs and CD11bhighGr-1+ MDSC overlaps.
AR: To clarify, supplemental Figures S1-2 show the specific gating protocol used to identify cell subsets. In short, NK cells are CD45+ CD3e- NK1.1+ based on gating but the backgating in Figure 3 shows that they are also B220-/low, which the text reflects. As for MDSC, binary gating of events along a specific marker axis can't result in overlapping populations. However, backgating the events along a different combination of marker axes can result in overlapping populations, which is pointed out in this critique. One can see from supplemental Figure S2 that the overlapping region is for cells that have intermediate CD11b but low SSC that fall outside of the CD11b+/SSC gating that are then positive for GR1.

3) In Figure 3N, the authors correlate the tumor size with infiltration of CD11b+Gr1+ events in various tumor models. However, the authors only use 2 ID mCCN4 + Dox tumors and the large YM1.7 KO tumors overlap with the small WT Yum1.7 tumors, making it difficult to interpret how tumor size influences MDSC recruitment.

AR: That figure is meant to show that, for similarly sized tumors, MDSC are more prevalent in tumors derived from WT compared to CCN4 KO cells (that is better shown

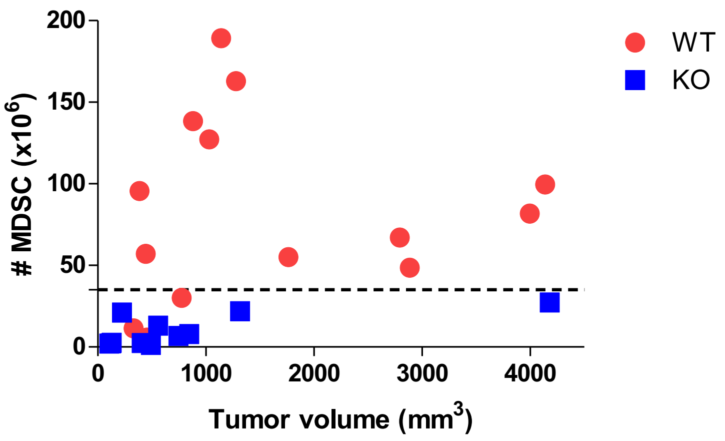


Figure C2: Number of live CD45+ CD11b+ Gr1+ (MDSC) isolated from the spleen of tumor bearing mice increased with a greater dependence on size of WT tumors compared with CCN4 KO tumors. Results aggregated for mice receiving WT (red circles) and CCN4 KO (blue squares) YUMM1.7 cells in a preliminary time-matched study where, following s.q. challenge with the indicated cell type, mice received an isotype control antibody or antibodies against PD1, CTLA4 or the combination starting at day 11. We redid this experiment using a size-matched experimental design, which is included in the manuscript.

in Figure 4A-B) and that the prevalence is even greater in ID mCCN4 + Dox tumors. As illustrated in Figure C2 (above), we have data where MDSCs were quantified in the spleen for a much broader range of tumor sizes that shows this dependence. In the interest of brevity, we didn't show that correlation as the dependence of MDSC on tumor size is well-established in the literature such that this figure doesn't really add much new.

4) In Figure 5, the authors chose to pursue expression of CXCL1 and CCL2 out of several of the differentially expressed cytokines in the Cytokine Array. Can they provide further rationale for choosing those specific two chemokines out of the entire chemokine set in the array? Further, it would be helpful to provide the ECAR and OCAR stress test profile provided by the Seahorse Assay to fully interpret the glycolytic flux.
AR: To clarify, the relative impact of CCN4 KO on the cytokines, chemokines, and growth factors assayed can be interpreted in terms of a z-score. A z-score for each orange dot in Figure 5A can be visually estimated by noting where a particular chemokine falls relative to the closest dotted line for the same x-value (i.e., the appropriate control that accounts for random chance). From that perspective, CCL2 is the most extreme (highest z-score) and CXCL1 is the third most extreme. IGFBP3 falls in between. Instead of showing a table of z-scores, we decided to present the data this way, as in addition to showing a z-score visually, the figure also shows the relative abundance of the assayed chemokines. As seen in Figure 5A, CXCL1 and CCL2 are more abundant than many of the other chemokines tested (i.e., they fall to the right side of the x-axis and thus have relatively high basal expression). We have edited the text to mention that both

differential and high basal expression motivated their selection. As mentioned in the text, characterizing the metabolic parameters in WT and KO YUMM1.7 cells was motivated by IGFBP3. We also assayed CCL5 in TCM and serum, which was assayed to have a similar basal abundance as CCL2 and CXCL1 but a lower z-score. CCL5 was not significantly different in TCM, serum from tumor bearing mice, and CD45- ex vivo tumor cells. In the interest of brevity, we did not include those data.

In terms of the Seahorse metabolic profiling, we have now included the timecourse ECAR time course profile that is used to estimate glycolysis and glycolytic capacity (new panels in supplemental Figure S8) shown in Fig C3 below:

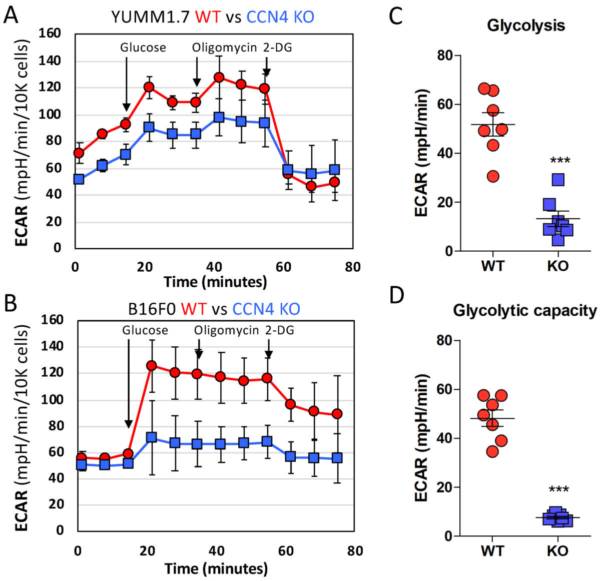


Figure C3 (new S8 in manuscript). Glycolysis and glycolytic capacity in YUMM1.7 and B16F0 cells were reduced upon CCN4 KO. Time course of ECAR analysis using XFe96 Seahorse Analyzer in WT and CCN4-KO tumors derived from YUMM1.7 (A) and B16F0 cells (B) after 36 hours ex vivo. (n = 3, results representative of one of three independent experiments). Curves represent average $\pm$ standard deviation. Time course profiles were analyzed to estimate (C) glycolysis and (D) glycolytic capacity for WT and CCN4-KO tumors derived from B16F0 cell variants. *** p<0.001. Error bars represent SEM.

5) In Figure 7, the authors seek to combine checkpoint blockade therapy with CCN4 knockout. They start treatment when tumors reach 100mm3 which happens to be later in the knockout model due to slowed growth kinetics. Given that PD-1 targeted monotherapy does not appear to be curative, if treatment was initiated earlier, might monotherapy be curative? Is CTLA-4/PD-1 combination synergistic?
AR: Of note, Hans Schreiber and coworkers published a meta-analysis suggesting that the earlier that therapy is started, the higher likelihood for getting a significant result (PMID: 22720238). Although, they question the clinical relevance of such early studies. That is why we focused on a size-matched experimental design where tumors were 100mm3 when treatment was initiated. We haven’t tested CTLA-4/PD-1 combination in YUMM1.7 in a size-matched study, which might be appropriate for a follow-on study.

6) Can the authors comment on effects of PD1 therapy in the knockout model? In the wild type tumor model, a clear difference in tumor size is seen between untreated and PD-1 treated mice. In the knockout model, the differences in tumor size are much smaller between the untreated and treated groups. In this case a significant difference is demonstrated 2 days after treatment, and while by day 3 the difference in tumor volume is still significant, the magnitude of the differences appears smaller than in the wild type setting.
AR: The experiment was designed to compare between +/- treatment for a given cell variant. We can say that the difference is significant. The study was not designed to make a statement about the difference in magnitude of the effect in WT versus KO. Note that because the treatment changes the slope of the curves midway through the experiment, we can't analyze the data in a similar way as shown in Figure 2.

Reviewer #2:
The current manuscript by Fernandez and colleagues describes a role for Cell Communication Network Factor 4 (CCN4) in promoting tumor-induced immunosuppression. This follows on from previous data from this group identifying this factor as a promotor EMT transition (Refs 11 and 12 from current manuscript). EMT transition has previously been shown to promote immunosuppression through a number of mechanisms and thus in the current study the authors explore the hypothesis that CCN4 directly regulates the immune response. The study is novel and of interest but there are technical limitations to the study that should be addressed to strengthen the conclusions.

7) Throughout the paper, the authors utilise a DNMT3a KO tumor as a control. What is the justification for this over a non-targeting control/ vector only control? Presuming the authors are referring to DNA (cytosine-5)-methyltransferase 3A, knockout of this gene would likely lead to significant transcriptional and epigenetic changes, thus making comparisons with this control irrelevant. This should be clearly explained and justified and experiments performed to ensure this "control" has no phenotype.

AR: In the interest of brevity, we only show these data for DNA (cytosine-5)-methyltransferase 3A-KO cells because the cells have also been successfully edited just at a different target. In terms of a difference in tumor growth in NSG versus C57BL/6, supplemental Figure S4 shows that it has no phenotype. Given that there is some overlapping biological activity with DNMT3B (e.g., PMIDs 10555141 and 12897133), the observation that DNMT3A KO alone has no phenotype in the context of B16F0 cells is maybe not so surprising. Of note in PMID 12897133, inactivation of both Dnmt3a and Dnmt3b and not single inactivation was required to get highly demethylated embryonic

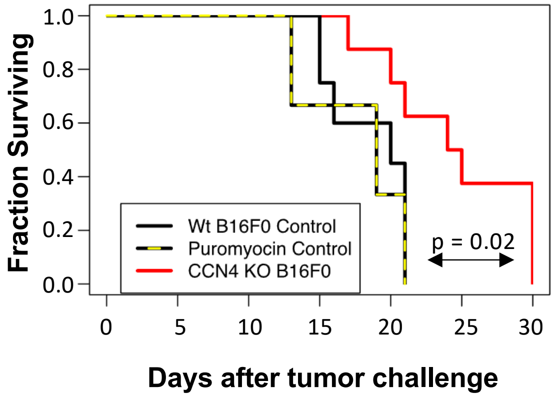


Figure C4. Controls for puromycin selection were created with pBabe-puro retrovirus (B16F0-Ctr) and injected subcutaneously in C57BL/6 mice (n=3, p-value vs WT (n = 8) = 0.6). Statistical significance for differences in overall survival were assessed using the Peto and Peto modification of a Gehan-Wilcoxon test.

stem cells, which could be restored by reintroducing isoforms of Dnmt3b. We have a follow-on manuscript that includes these DNMT3A data. In addition, we did a small pilot study using a non-targeting puromycin control, as shown in Figure C4. However, in the interest of brevity and the small size of this study, we decided to include only the DNMT3A data.

8) In Figure 2E none of the mice challenged with YUMM1.7 tumors achieve long-term survival but in Figure 4 this appears to be 20-40% (without or with Dox respectively). How do the authors explain this? One explanation is that the rtta element used for the inducible expression of CCN4 is immunogenic in wild-type mice (refer to PMID 20155386). If this is the case it would be a significant caveat for these experiments.
AR: To clarify, panels J-N in Figure 3 describe a rescue experiment where the base cell line used in panel K is Ym1.7-KO1 not WT YUMM1.7. Long-term survival of the ID vector control +/- dox is similar to KO1 shown in Figure 2E, where 7 of 23 mice injected were tumor free (30%) at the end of the experiment.

9) Several of the data are interesting but quite descriptive e.g. reduced expression of chemokines associated with immunosuppression, reduced MDSCs, reduced T cell activation in vitro. However, the mechanism by which CCN4 leads to enhanced tumor growth is not defined. As a minimum the author should present data with T cell

depletion/ targeting one of these factors to identify the key driver of the in vivo phenotype.

AR: We think the mechanism is multi-faceted. A manuscript that we have recently re-submitted to Nature Biomedical Engineering (see bioRxiv 2020.05.04.077107; doi: https://doi.org/10.1101/2020.05.04.077107) helps propose a mechanism based on human data that we think CCN4 has a direct effect on NK and CD8 T cell activity, skews the myeloid compartment, and alters the phenotype of fibroblasts. The data obtained using mouse melanoma models parallel these findings. Here KO eliminates the source from tumor cells while the rescue of CCN4 expression recovered the phenotype. Given the potential complexity that underpins this phenotype, we feel that parsing the different aspects of immunosuppression in these systems is best left for follow-on papers – as mentioned by Reviewer 3. From a practical perspective, the first author (Audry Fernandez) has left the lab and this manuscript largely reflects her contributions. Given the importance of first-author publications for the careers of junior scientists, asking a junior scientist to invest a significant effort to clarify mechanisms without the opportunity for being first author would not be fair to them.

10) In Figure 1 the authors present data suggesting that Knockout of CCN4 leads to greater tumor control. Whilst it is useful to have the individual growth kinetics from each mouse, this should be supported by data indicating the mean +/- SEM for these experiments to understand whether these differences are statistically significant. The presented formula attempts to do this, but is not clear why such an equation is needed to confirm this important point.

AR: Traditional statistical methods used in biological research today focus largely on the idea of rejecting a null hypothesis and eliminating the possibility that a result seen was the product of chance alone. Specifically, the proposed null hypothesis to be tested here is that the tumor size values observed at a single time point for mice challenged with WT cells are the same as for mice challenged with KO cells. Assuming that these observations can be represented by a normal distribution, a p-value of less than 0.05 implies that these observations for the two different conditions can not be explained by random chance. Implicitly one then accepts an alternative hypothesis but doesn't state what that hypothesis is. Here, one could come up with at least two competing alternative hypotheses: 1) that WT versus KO cells contain a different number of tumor initiating cells and that these cell variants grow at the same rate OR 2) that WT versus KO cells contain the same number of tumor initiating cells but grow at different rates once microscopic tumors are established. By presenting the data as mean +/- SEM plus a t-test to assess statistical significance (of note, that is shown in Figure 2I), one can conclude only that the results are not explained by random chance. In contrast, the Bayesian approach used here is designed to specifically focus on the observed data and testing a description of what we believe is different – that the growth rate is different. For a more in-depth explanation, refer to the previous manuscript: **In silico model-based inference: A contemporary approach for hypothesis testing in network biology (D. Klinke, Biotechnol Prog, 2014 DOI: 10.1002/btpr.1982)**. An explanation of conventional hypothesis testing and its pitfalls begins on page 5, while a brief explanation of the Bayesian approach that is illustrated by this and other works by Klinke Lab begins on page 9.

Specifically, in Figure 2, we are using multiple measures of tumor size obtained for each individual mouse with time to estimate both the initial tumor bolus and rate constant. We present these parameter values as posterior distributions – that is the distribution of parameter values that are consistent with the data, given our explicit statement of how we think this system can be described (i.e., the mathematical model) and the uncertainty associated with the experimental measurements. By approaching the problem in this way, we are not constraining the parameters in any way other than through the data collected in the mouse studies and we are allowing the parameters to best fit all data gathered across time points rather than being skewed by possible outliers. Statistical significance of the differences in the rates of growth between wildtype and knock out groups is calculated by a Chi-squared test with simulated p-value, which accounts for the shift in the posterior distributions between groups.

11) The authors suggest that targeting CCN4 enhances the effect of immune checkpoint blockade. However this is not borne out by the data where the magnitude of response is similar. I suggest this data is removed or conclusions modified.
AR: It seems that the effects are additive. We have modified the text such that we don't imply that these effects are synergistic (i.e., multiplicative).

Minor Points

12) Figure 1A is reproduced from one of the author's previous publications. Whilst this is not a major issue given it is a bioinformatics analysis, this should be acknowledged in the figure legend.
AR: While the particular graphic is unique to this manuscript, we have included a reference to this prior citation in the legend.

13) Several of the figure panels are called out non sequentially.
AR: We have relabeled and reordered some of the figures so that the panels are called out sequentially.

Reviewer #3:

This is an elegant paper on the effects of CCN4 in melanoma. The manuscript is clearly written, and the figures are clearly displayed to illustrate particular points made in the manuscript. The authors show that CCN4 acts as an immune-suppressor in melanoma. The authors propose that the main activity of CCN4 is through the inhibition of MDSC function. They demonstrate both direct effects, such as release by melanoma cells of chemokines to recruit MDSCs, and, inhibition of IFNg release by T cells. Checkpoint inhibitor therapy potentiates the effects of CCN4 knock-out.

14) Criticisms: the effect of knock-down in B16FO cells seems to be marginal at best, and is therefore not explored further. Does this really make a significant contribution to the

manuscript. If not, I suggest to remove these data, as they reduce the impact of all other data shown.
AR: Of note, there is a significant difference in tumor growth upon CCN4 KO in B16F0 cells. We have included the B16F0 data for multiple reasons. First, our prior work in JBC 2019 and Cell Mol Bioeng 2020 that focused on metastatic roles of CCN4 used both B16F0/F10 and YUMM1.7 cells. If we left out the B16F0 data, the obvious question would be why we didn't include those data (i.e., maybe it didn't work) and that we are selectively presenting data to spin the story in the most positive light. Second, the B16 model is thought to be the gold standard for testing cancer immunotherapies in mice because very few therapies work with it. We hazard to guess that one reason that very few therapies work is that there are multiple immunosuppressive mechanisms active in that model. The B16 model is of spontaneous origin which means that the random process associated with genetic mutation and selection favored the development of multiple suppressive mechanisms in these malignant cells. In contrast, the YUMM models originate from GEMM models, where oncogenic mutations are genetically introduced and shortcut the natural mutation/selection process. Third, finding something new that has an impact on tumor growth using the YUMM1.7 model may not be that difficult as this cell model was recently developed. In contrast, finding something new using the B16 model is much more difficult as it has been so widely studied. In the interest of providing a more complete story, we decided to include both the B16F0 and YUMM1.7 data.

15) The ultimate mechanism by which a matricellular protein affects the release of chemokines is not explored herein, and can safely be left for a future paper. Nevertheless, the authors are encouraged to speculate on this potential mechanism.
AR: Overall, we tried to be concise, given the word limits for manuscripts. As we mentioned in the discussion, we think that CCN4 induces EMT that is mediated by upregulation of SNAI1. These relationships are covered in greater detail in our recent JBC and Cell and Mol Bioeng papers. We also mentioned in the discussion that SNAI1 expression is associated with the production of CXCL1 and CCL2. Follow on studies will focus on elaborating these details.

**Subject:** Re: CANCER RESEARCH CAN-20-3478R1 -- Decision Rendered
**Date:** Tuesday, March 2, 2021 at 3:52:07 PM Eastern Standard Time
**From:** David Klinke
**To:**

Hi Dr. Van Dang,

Thank you for your response. So to clarify, Reviewer #3 that was supportive in the first round had no additional comments. Reviewer #1 felt that we addressed all of their comments. Reviewer #2 has the following concerns that we will address below:

**Reviewer #2 Comments to Author:**
The reviewers have addressed some of the points raised in the initial review but there are still concerning technical and mechanistic limitations as detailed in my responses below.

1. Reviewer comment and author responses in round 1 about DNMT3a KO tumor as a control. (see complete text at bottom of email).

   New Reviewer response: I believe that this is an unacceptable control for these studies as it would be impossible to be confident that DNMT3A deletion has no effect on the phenotype of B16F10 based upon these readouts alone. A number of studies have shown a phenotype in alternative cell types with a single DNMT3A knockout (PMID 25416277, 24589714). Therefore the non-targeting puromycin control data should be used to replace the DNMT3A data.

   New Author Response: To be clear here, we are saying that DNMT3A deletion has no effect on the in vivo tumor growth phenotype of B16F0 cells. That said, let's dig into this critique where "a number of studies" (2 references given) are cited related to DNMT3A. One of these citations is a review article from 2014 (PMID 24589714). This review covers general aspects of DNMT3A biology and correlative clinical studies. When it turns to discussing the role of DNMT3A in cancer, the authors state that the mechanism underlying the promotion of tumorigenesis by DNMT3A (based on clinical correlates) is still unknown. The only primary research cited using in vivo models is a paper from 2009 where they knocked down DNMT3A in B16 cells by transfecting in a shRNA plasmid (PMID 19632198). They found that tumors in C57Bl/6 mice generated from B16 clones that harbored the shRNA against DNMT3A were significantly smaller than WT or negative control (containing a "non-target" shRNA) cells. They also did a microarray analysis of the bulk tumors and found that the most significantly enriched pathways were associated with immunity and defense, as one would expect if an interferon pathway was triggered. One of the things with older papers is that, in time, we understand the limitations of some of the prior experimental techniques used. One of the limitations of shRNA is that it has off-target effects and transfecting in a shRNA plasmid can engage TLRs eliciting an interferon response. To get around that, the most rigorous approach is to do a rescue experiment with an RNAi-resistant target gene (see Curr Protoc Cell Biol 2010 – PMID 21154550). The 2009 paper did not do the rescue experiment, which raises doubts about how the results were interpreted.
   The other citation (PMID: 25416277) refers to a study where Dnmt3a is knocked out in mouse hematopoietic stem cells and then these mouse Dnmt3a-KO HSC are injected into sublethally irradiated host mice. They found that mice receiving Dnmt3a-KO cells had a median survival of 321 days and developed a range of myeloid and lymphoid diseases, where the malignancies contain a high frequency of secondary mutations. Interestingly, they show in Figure 4B that survival of mice receiving either WT HSC transduced with a GFP vector or Dnmt3a-KO HSC transduced with a GFP vector was the same (100% of the mice from both groups survived to 90 days). Putting aside concerns about whether a long-term study focusing on hematopoietic stem cells is relevant for malignant

melanocytes, these data support that Dnmt3a knockout alone has no phenotype in terms of in vivo growth differences. This is consistent with what we have observed.

2. Reviewer comment and Author Response in round 1 to critique about CCN4 rescue experiment using Ym1.7-KO1 cells. (see mail email for complete text).
   New Reviewer response: Thank you for clarifying the details of the tumor model. However, the data are still confusing. Doxycycline induced expression of CCN4 reduces the survival of mice, potentially consistent with an enhanced immunosuppressive phenotype. But why do the tumor types grow differently in the absence of doxycycline treatment.
   New Author Response: Of note, the rtta element is present in both Ym1.7-KO1 variants modified to contain the control ID vector or the ID mCCN4 vector. As to why do YUMM1.7-KO1-iCCN4 tumors grow differently than YUMM1.7-KO1-IV tumors both in the absence of dox could be explained by the observation that YUMM1.7-KO1-iCCN4 cells seem to be more immunogenic that YUMM1.7-KO1-IV cells (see ELISpot results in Figure 6). Of note, YUMM1.7 cells are thought to be not highly immunogenic, which led to developing YUMMER1.7 cells (PMID 28379630). That said, the 2x2 factorial experimental design coupled with multivariate Cox PH modeling in this experiment enables parsing the contribution of dox and the particular insert vector from the expression of CCN4 in YUMM1.7-KO1 cells.

3. Reviewer comment in round 1: Several of the data are interesting but quite descriptive e.g. reduced expression of chemokines associated with immunosuppression, reduced MDSCs, reduced T cell activation in vitro. However, the mechanism by which CCN4 leads to enhanced tumor growth is not defined. As a minimum the author should present data with T cell depletion/ targeting one of these factors to identify the key driver of the in vivo phenotype.
   Author Response in round 1: We think the mechanism is multi-faceted. A manuscript that we have recently re-submitted to Nature Biomedical Engineering (see bioRxiv 2020.05.04.077107; doi: https://doi.org/10.1101/2020.05.04.077107) helps propose a mechanism based on human data that we think CCN4 has a direct effect on NK and CD8 T cell activity, skews the myeloid compartment, and alters the phenotype of fibroblasts. The data obtained using mouse melanoma models parallel these findings. Here KO eliminates the source from tumor cells while the rescue of CCN4 expression recovered the phenotype. Given the potential complexity that underpins this phenotype, we feel that parsing the different aspects of immunosuppression in these systems is best left for follow-on papers – as mentioned by Reviewer 3. From a practical perspective … would not be fair to them.
   New Reviewer response: Whilst I sympathize that the mechanism may be complex to unravel, and on the restraints for junior scientists, I feel there are some simple experiments that could be performed to address this and provide important insight.

New Author Response: Couple of comments here. First, the comparison of WT versus CCN4 KO variants of both B16 and YUMM1.7 in C57Bl/6 and NSG mice establishes that CCN4 influences the host anti-tumor immune response. These results help establish bounds for what is possible – that myeloid cells, NK cells, CD4 T cells, and CD8 T cells play roles in controlling tumor growth in the absence of CCN4. So then, would depleting a T cell subset or targeting a chemokine using an antibody identify that this particular component is a key driver of the in vivo phenotype? In short, no. Let's take an example. Let's say we deplete CD8 T cells in mice challenged with YUMM1.7-KO1 cells. If a statistically significant result was obtained, all this would show is that the results would not be explained by random chance, that is to reject the null hypothesis. Appropriate interpretation would only be that reducing CD8 T cells has a significant impact on tumor growth. Would this mean that myeloid cells or NK cells are not important? No it wouldn't as one should do additional experiments that target those cells. If no statistically significant result was obtained, would this mean that CD8 T cells are not important. Not necessarily, it may mean that both NK and CD8 T cells contribute to cytotoxic function within the tumor microenvironment. Alternatively, say you neutralize a chemokine using an antibody. If you obtained a statistically significant result, would this mean that the other chemokines are not important? No, it wouldn't because that is not what was tested. Alternatively, if you didn't obtain a statistically significant result, would this mean that the targeted chemokine is not important? No, it wouldn't because maybe the dose of antibody wasn't appropriate or maybe the chemokines have overlapping functions. The point here is that there are no "simple experiments" that one could do here that would provide important insight. These immunologic details need to be addressed in a separate manuscript where alternative explanations can be explored and tested rigorously.

So in summary, there don't appear to be any meaningful critiques of this manuscript. The comment about a "simple experiment" to identify the key driver, in my view as a standing member of the Cancer Immunopathology and Immunotherapy study section at NIH, seems a bit naïve. Given these data, would you reconsider publishing this manuscript in Cancer Research?

Sincerely,
David Klinke
—

David J. Klinke II, Ph.D.
Professor - Department of Chemical and Biomedical Engineering
Adjunct Associate Professor - Department of Microbiology, Immunology & Cell Biology
West Virginia University, Morgantown, WV 26506-6102
(304) 293-9346 | ESB 517 | WVUCI 1838 | Website: http://klinke.openwetware.org

---

**From:** Cancer Research <>
**Reply-To:** "" <>
**Date:** Monday, March 1, 2021 at 9:40 AM
**To:** David Klinke <>
**Subject:** CANCER RESEARCH CAN-20-3478R1 -- Decision Rendered

Re: CAN-20-3478R1
Cell Communication Network factor 4 promotes tumor-induced immunosuppression in melanoma.

Dear Dr. Klinke,

Thank you for submitting your above-referenced manuscript to Cancer Research. We have now completed the review process and I regret to inform you that your manuscript will not be accepted for publication in Cancer Research. Comments from our Reviewers and/or Editors are offered for your information below.

Final decisions rendered on submissions to the journal are based on the Editor-in-Chief's or the Senior Editor's overall assessment of all of the information obtained on a manuscript during the review process. In addition to Reviewer comments, the Editorial Board member's recommendation and the priority scores assigned to the work are also critical. All submissions must meet stringent considerations of scientific priority, breadth, quality, and originality. They must also be judged to exert a broad impact of general interest to the diverse audience of the journal. Unfortunately, for this particular manuscript, it was felt that these criteria were not fulfilled to the extent needed for acceptance.

Although we cannot offer publication of your manuscript in *Cancer Research*, we approached in confidence the editors of another AACR journal, *Molecular Cancer Research*, to ascertain their level of interest. After careful consideration, those editors have agreed that your manuscript is of substantial interest to them, and they would like to encourage you to transfer it to *Molecular Cancer Research*. They are happy to consider it in the context of the *Cancer Research* reviews, and they are interested in publishing your manuscript without further major revision. If you would like to discuss this option further, please reach out to the Associate Editor of *Molecular Cancer Research*, Lucas Brand. To learn more about offers made through our transfer service, please access the instructions below. If you would like to transfer your manuscript and its reviews to *Molecular Cancer Research*, please use the link below.

Alternatively, you may use the transfer service to send your manuscript to any other AACR journal of your choice, we can only offer insight into *Molecular Cancer Research*'s interest in the work.

I regret that a more favorable decision could not be offered on this occasion. Nevertheless, on behalf of all the Editors of Cancer Research, I would like to express our true appreciation for the opportunity to receive and consider your manuscript, and our keen desire to welcome future submissions from your group to Cancer Research.

Sincerely,

Chi Van Dang, MD, PhD
Editor-in-Chief

**AACR Manuscript Transfer Service**
The AACR offers authors the option to easily transfer a declined Research Article to a second AACR journal of your choice for editorial consideration. At your request, all files, manuscript information, and reviews (if applicable) will be transferred immediately. For details of this service, please visit the AACR Manuscript Transfer FAQ page.

Click the link below to view your options and (if desired) begin the transfer process.

https://can.msubmit.net/cgi-bin/main.plex?el=DELETED

Comments from Reviewers:
Reviewer #1 (Reviewer Comments to the Author):

The authors have adequately addressed all of this reviewer's comments

Reviewer #2 (Reviewer Comments to the Author):

The reviewers have addressed some of the points raised in the initial review but there are still concerning technical and mechanistic limitations as detailed in my responses below.

Reviewer comment: Throughout the paper, the authors utilise a DNMT3a KO tumor as a control. What is the justification for this over a non-targeting control/ vector only control? Presuming the authors are referring to DNA (cytosine-5)-methyltransferase 3A, knockout of this gene would likely lead to significant transcriptional and epigenetic changes, thus making comparisons with this control irrelevant. This should be clearly explained and justified and experiments performed to ensure this "control" has no phenotype.

Author response: In the interest of brevity, we only show these data for DNA (cytosine-5)-methyltransferase 3A-KO cells because the cells have also been successfully edited just at a different target. In terms of a difference in tumor growth in NSG versus C57BL/6, supplemental Figure S4 shows that it has no phenotype. Given that there is some overlapping biological activity with DNMT3B (e.g., PMIDs 10555141 and 12897133), the observation that DNMT3A KO alone has no phenotype in the context of B16F0 cells is maybe not so surprising. Of note in PMID 12897133, inactivation of both Dnmt3a and Dnmt3b and not single inactivation was required to get highly demethylated embryonic stem cells, which could be restored by reintroducing isoforms of Dnmt3b. We have a follow-on manuscript that includes these DNMT3A data. In addition, we did a small pilot study using a non-targeting puromycin control, as shown in Figure C4. However, in the interest of brevity and the small size of this study, we decided to include only the DNMT3A data.

Reviewer response: I believe that this is an unacceptable control for these studies as it would be impossible to be confident that DNMT3A deletion has no effect on the phenotype of B16F10 based upon these readouts alone. A number of studies have shown a phenotype in alternative cell types with a single DNMT3A knockout (PMID 25416277, 24589714). Therefore the non-targeting puromycin control data should be used to replace the DNMT3A data.

Reviewer Comment: In Figure 2E none of the mice challenged with YUMM1.7 tumors achieve long-term survival but in Figure 4 this appears to be 20-40% (without or with Dox respectively). How do the authors explain this? One explanation is that the rtta element used for the inducible expression of CCN4 is immunogenic in wild-type mice (refer to PMID 20155386). If this is the case it would be a significant caveat for these experiments.

AR: To clarify, panels J-N in Figure 3 describe a rescue experiment where the base cell line used in panel K is Ym1.7-KO1 not WT YUMM1.7. Long-term survival of the ID vector control +/- dox is similar to KO1 shown in Figure 2E, where 7 of 23 mice injected were tumor free (30%) at the end of the experiment.

Reviewer response: Thank you for clarifying the details of the tumor model. However, the data are still confusing. Doxycycline induced expression of CCN4 reduces the survival of mice, potentially consistent with an enhanced immunosuppressive phenotype. But why do the tumor types grow differently in the absence of doxycycline treatment.

Reviewer Comment: Several of the data are interesting but quite descriptive e.g. reduced expression of chemokines associated with immunosuppression, reduced MDSCs, reduced T cell activation in vitro. However, the mechanism by which CCN4 leads to enhanced tumor growth is not defined. As a minimum the author should present data with T cell depletion/ targeting one of these factors to identify the key driver of the in vivo phenotype.

Author Response: We think the mechanism is multi-faceted. A manuscript that we have recently re-submitted to Nature Biomedical Engineering (see bioRxiv 2020.05.04.077107; doi: https://doi.org/10.1101/2020.05.04.077107) helps propose a mechanism based on human data that we think CCN4 has a direct effect on NK and CD8 T cell activity, skews the myeloid compartment, and alters the phenotype of fibroblasts. The data obtained using mouse melanoma models parallel these findings. Here KO eliminates the source from tumor cells while the rescue of CCN4 expression recovered the phenotype. Given the potential complexity that underpins this phenotype, we feel that parsing the different aspects of immunosuppression in these systems is best left for follow-on papers - as mentioned by Reviewer 3. From a practical perspective, the first author (Audry Fernandez) has left the lab and this manuscript largely reflects her contributions. Given the importance of first-author publications for the careers of junior scientists, asking a junior scientist to invest a significant effort to clarify mechanisms without the opportunity for being first author would not be fair to them.

Reviewer response: Whilst I sympathize that the mechanism may be complex to unravel, and on the restraints for junior scientists, I feel there are some simple experiments that could be performed to address this and provide important insight.

Reviewer Comment: In Figure 1 the authors present data suggesting that Knockout of CCN4 leads to greater tumor control. Whilst it is useful to have the individual growth kinetics from each mouse, this should be supported by data indicating the mean +/- SEM for these experiments to understand whether these differences are statistically significant. The presented formula attempts to do this, but is not clear why such an equation is needed to confirm this important point.

Author Response: Traditional statistical methods used in biological research today focus largely on the idea of rejecting a null hypothesis and eliminating the possibility that a result seen was the product of chance alone. Specifically, the proposed null hypothesis to be tested here is that the tumor size values observed at a single time point for mice challenged with WT cells are the same as for mice challenged with KO cells. Assuming that these observations can be represented by a normal distribution, a p-value of less than 0.05 implies that these observations for the two different conditions can not be explained by random chance. Implicitly one then accepts an alternative hypothesis but doesn't state what that hypothesis is. Here, one could come up with at least two competing alternative hypotheses: 1) that WT versus KO cells contain a different number of tumor initiating cells and that these cell variants grow at the same rate OR 2) that WT versus KO cells contain the same number of tumor initiating cells but grow at different rates once microscopic tumors are established. By presenting the data as mean +/- SEM plus a t-test to assess statistical significance (of note, that is shown in Figure 2I), one can conclude only that the results are not explained by random chance. In contrast, the Bayesian approach used here is designed to specifically focus on the observed data and testing a description of what we believe is different - that the growth rate is different. For a more in-depth explanation, refer to the previous manuscript: In silico model-based inference: A contemporary approach for hypothesis testing in network biology (D. Klinke, Biotechnol Prog, 2014 DOI: 10.1002/btpr.1982). An explanation of conventional hypothesis testing and its pitfalls begins on page 5, while a brief explanation of the Bayesian approach that is illustrated by this and other works by Klinke Lab begins on page 9.

Specifically, in Figure 2, we are using multiple measures of tumor size obtained for each individual mouse with time to estimate both the initial tumor bolus and rate constant. We present these parameter values as posterior distributions - that is the distribution of parameter values that are consistent with the data, given our explicit statement of how we think this system can be described (i.e., the mathematical model) and the uncertainty associated with the experimental measurements. By approaching the problem in this way, we are not constraining the parameters in any way other than through the data collected in the mouse studies and we are allowing the parameters to best fit all data gathered across time points rather than being skewed by possible outliers. Statistical significance of the differences in the rates of growth between wildtype and knock out groups is calculated by a Chi-squared test with simulated p-value, which accounts for the shift in the posterior distributions between groups.

Reviewer response: Thank you for this detailed explanation.

Reviewer question: The authors suggest that targeting CCN4 enhances the effect of immune checkpoint blockade. However this is not borne out by the data where the magnitude of response is similar. I suggest this data is removed or conclusions modified.

Author Response: It seems that the effects are additive. We have modified the text such that we don't imply that these effects are synergistic (i.e., multiplicative).

Reviewer response: Thank you for this amendment.
